# Immunity Depletion, Telomere Imbalance, and Cancer-associated Metabolism Pathway Aberrations in Intestinal Mucosa upon Caloric Restriction

**DOI:** 10.1101/2021.03.10.433216

**Authors:** Evan Maestri, Kalina Duszka, Vladimir A Kuznetsov

## Abstract

Systematic analysis of calorie restriction (CR) mechanisms and pathways in cancer biology has not been carried out, leaving therapeutic benefits unclear. Using a systems biology approach and metadata analysis, we studied gene expression changes in the response of normal mouse duodenum mucosa (DM) to short-term (2-weeks) 25% CR as a biological model. We found a high similarity of gene expression profiles in human and mouse DM tissues. Surprisingly, 26% of the 467 CR responding differential expressed genes (DEGs) in mice consist of cancer-associated genes—most never studied in CR contexts. The DEGs were enriched with over-expressed cell cycle, oncogenes, and metabolic reprogramming pathways (MRP) that determine tissue-specific tumorigenesis, cancer, and stem cell activation; tumor suppressors and apoptosis genes were under-expressed. DEG enrichments suggest a misbalance in telomere maintenance and activation of metabolic pathways playing dual (anti-cancer and pro-oncogenic) roles. Immune system genes (ISGs) consist of 37% of the total DEGs; the majority of ISGs are suppressed, including cell-autonomous immunity and tumor immune evasion controls. Thus, CR induces MRP suppressing multiple immune mechanics and activating oncogenic pathways, potentially driving pre-malignant and cancer states. These findings change the paradigm regarding the anti-cancer role of CR and may initiate specific treatment target development.

## Introduction

Calorie restriction (CR), where test animals receive a reduced energy diet, is one of the most broadly acting regimens for preventing or reversing weight gain and inhibiting cancer in experimental tumor models (1,2). Protocols typically involve 10 to 40% reductions in total energy intake compared to ad libitum-fed controls, but with adequate nutrition and a controlled physical environment (2). Chronic reduction of dietary energy intake without malnutrition decreases adiposity, inflammation and improves metabolic profiles (3,4). CR was shown to increase tumor latency and have protective effects in some experimental mammary carcinogenesis models (5,6). Upon CR, metabolic alterations foster health-promoting characteristics including increased insulin sensitivity, decreased blood glucose and growth factors (IGF-1), and angiogenesis (4). Reducing IGF-1 and glucose levels may decrease tumor progression (2–4,7). Large bodies of experimental preclinical models support CR as anti-tumorigenic (2–4,7). Yet, data has also demonstrated neutral and negative effects (8–10). For instance, mice small intestinal response had a highly-dispersal trend decreasing large polyps (> 2mm) but increasing small polyp (≤2 mm) numbers (8). CR started early in life reduced incidence/delayed progression in most rodent tumors, however, CR started in middle-aged mice had higher lifetime incidences of lymphatic neoplasms (10). Whether CR results in protective or deleterious effects on cancer risk and outcome depends on length and restriction severity (6,7). A dominating anti-cancer CR effect decreases tumor rate growth via inhibition of circulating systemic factors (e.g., hormones/growth factors) which stimulate cancer cell proliferation (5,11). However, many cancer subtypes lack hormone/growth factor sensitive cells (e.g., high aggressive basal-like cancers). Sensitive tumor clone(s) may be targets of pro-oncogenic metabolic reprogramming and/or replaced with more aggressive CR-resistant ones.

Recent randomized clinical trials have been explored for potential CR anti-cancer properties (12–14). These trials showed mixture effects of CR directly on tumor tissue growth, host immune cells response, and other tissue responses (e.g., adipocytes). However, these studies do not mechanistically support the anti-cancer role of CR on neoplastic processes as indicated by large bodies of empirical experimental model data. CR-mediated reduction in cancer cell proliferation is central to anti-cancer animal model studies. The clinical trial studies showed no negative energy/calorie restriction effects on cell proliferation marker Ki-67 in Barrett’s esophagus (13) and breast cancers (14). In men with prostate cancer, presurgical weight loss showed CR-mediated Ki-67 upregulation (12). Additionally, high overexpression of several periodic cell cycle genes, cancer-associated genes, and oncogenes have been found (12,14).

The duodenum mucosa (DM) is a useful biological model for the study of CR small intestine response (15). After epithelial cells, the most numerous cells in the lamina propria are immune cells, mainly duodenal intraepithelial lymphocytes (IEL) (16,17), present at 9-50 IELs per 100 epithelial cells that vary in different medical and bacterial conditions (17,18). The DM transcriptome model showed dichotomized differentially expressed genes (DEGs) response to CR, with metabolic genes upregulated and immune/inflammatory genes downregulated (15). However, DM tissue-specific transcriptional network profiles in CR-medicated cancer biology mechanics were not studied. The intestinal mucosa epithelium is the most highly proliferative mammalian tissue and CR further enhances epithelial regeneration (19,20). Precise balances control the quiescent (G0-phase) and active intestinal stem cells, progenitors, and stroma cells. In healthy mice, short-term CR reduced cellular mass resulting in 15% shorter villi, reduced numbers of more differentiated epithelial progenitor cells, while the proliferative rate and the number of Lgr5 (high)/Olfm4+ active intestinal stem cells were modestly increased (21). Isolated crypts from CR-restricted mice *in vitro* can form primary and secondary organoid bodies with increased proliferation rate and stem cell growth per crypt (21). Cycling intestinal stem cells exhibit high Wnt activity that sensitizes them to DNA damage after CR (21,22). This suggests CR initiates microenvironmental/stroma-mediated control loss, activates independent proliferation, and stimulates progression of stem cell organoid body formation—cancer hallmark factors.

The direct CR anti-cancer and pro-cancer mechanics in normally proliferated intestinal mucosa are poorly understood. Systematic analysis of CR mechanisms in cancer biology has not been carried out, making therapeutic benefits unclear (12,23,24). The contradictive tumor biology CR data motivated us to use hypothesis-testing and data-driven system biology analysis of gene expression profiles changes in small intestine mucosa tissue upon short-term CR.

Our major objective is to identify cancer driver genes and oncogenic pathways induced by CR-related perturbation of cellular homeostasis affecting normal proliferation of epithelium mucosa via metabolic reprogramming and depletion of immune system control. Our central hypothesis is that CR activates metabolic reprograming pathways in DM defined by cancer hallmarks (e.g., uncontrolled cell proliferation, tumor suppressors loss etc.), while suppressing immune system surveillance; this may initiate occurrence or preferential competition of abnormal proliferating cells leading to pre-cancer and cancer risks. To test this hypothesis, we developed unbiased and comprehensive data analysis approaches. We use the mouse mucosa response to 2-week 25% CR model in DM. Our results demonstrated CR induces drastic gene expression and pathway suppression of intracellular immunity and immune responses of T-, B-, NK-cells, macrophages and their precursors. We identify and characterize CR DEGs specifying regulatory networks modulating telomere stability and tumor suppressors, activating proliferative tissuespecific cancer-like stem cells, oncogenes, and chemical carcinogenesis. New CR response genes, perspective treatment onco-targets, and cancer risk factors are discussed. Our CR-induced metabolic reprogramming and multi-cellular competition models suggest plausible dysregulation of genes, networks, and pathways of pre-malignant and malignant states.

## Materials and methods

In this study, we carried out data analysis of differentially expressed genes (DEGs) of 26,966 probe sets (p.s.) from Affymetrix MoGene 1.0 ST microarray data from mouse mucosa scrapings produced in our previous study (15). Using statistical criteria reported in (15), 521 p.s. were selected. 16 p.s without gene annotations were excluded. We determined 505 probe sets (p.s.) representing CR mice DEGs, with 467 unique annotated gene symbols (Supplementary Data 1A-B). Adjusted p-values ≤ 0.05 and fold-changes ≥ 1.5 were the thresholds for statistical significance. The gene annotation available for mouse MGI DB (mm9/GRm38 assembly, http://www.informatics.jax.org/marker/) and microarray datasets were mapped using BLASTN and BLASTP to mouse genome and transcriptome respectively, manually curated, re-annotated and in some cases annotated de novo. We analysed gene sequences and annotated gene symbols associated with multiple p.s. We analysed normal human duodenum gene expression profiles (25) using statistical methods (Suppl. Methods) and studied correlations in the gene expression of mouse and human DM. We integrated other datasets using bioinformatics recourses (Suppl. Methods).

### Computational Resources

Gene enrichment subset analysis, networks, pathways, collections of DBs, list publications used during manual curation of genes, and our selection criteria and statistical methods are described in Suppl. Methods. In this study, we systematically used Ingenuity Pathway Analysis (IPA) tools (QIAGEN Inc., https://www.qiagenbioinformatics.com/products/ingenuity-pathway-analysis) (26). For the annotation, GO terms, and network connectivity enrichment analysis we also used: DAVID Bioinformatics 6.7 (https://david-d.ncifcrf.gov/), STRING v11 (https://string-db.org), Enrichr DB (https://amp.pharm.mssm.edu/Enrichr/).

## Results

### Analysis of differentially expressed genes in mouse duodenum mucosa

In our previous study, we carried out gene microarray expression profiling and its experimental validation for CR-responded DEGs in mouse DM (15). In this work, we reanalyzed the microarray dataset using advanced bioinformatics and systems biology approaches to test our central hypothesis. In CR mice compared to *ad libitum* mice a total of 240 p.s. were significantly upregulated and 265 p.s. were downregulated; there are 222 upregulated unique gene symbols and 246 downregulated unique symbols. Supplementary Data 1 indicates CR DEG categories.

Enrichment analysis of tissue-associated proteins was completed using the Uni-Prot datasets (Up_Tissue) annotation database (https://www.uniprot.org/). Through the selection of tissues with Benjamini < 0.05, the Enriched Immune System Gene (ISG) subset was represented by 171 genes (132 downregulated, 39 upregulated) that referred to the jejunal and colic lymph nodes (at 87.14-fold enrichment), spleen, activated spleen, and thymus. Epithelial Cell-Enriched Genes (ECG) were represented by 152 genes (52 downregulated, 100 upregulated) that referred to the colon, liver, kidney, and SI. Additional immune resources (27) and IPA’s Disease and Biological Function annotations strengthened the classification system. The DEGs not included in the previous categories were called “Other Genes”. GO Analysis via DAVID Bioinformatics of the CR DEGs (Supplementary Data 2) revealed top biological processes and pathways including immune response (p = 5.45 10^−7^) and glutathione transferase activity (p = 2.93 10^−9^).

### DM tissue expressed gene are translatable between mouse and human

The translatability between experimental models was analyzed by comparison of microarrays data from the CR mouse duodenum to normal human duodenum expression (**Figure 1A**) (25). We found 17,760 expressed genes in the normal human duodenum and 20,727 in the normal mouse duodenum (Methods). The number of shared orthologous expressed genes between normal human and mouse duodenum was 15,224; this represents 85.72% (15224/17760) of the entire human DM expressed genes and 73.45% (15224/20727) of the entire mouse DM expressed genes. Additionally, 382 of the 467 mouse CR DEGs had human orthologs. High levels of similarity exist between the duodenum gene lists for mouse and human. GO and pathway analysis via STRING v11 determined the functional associations between the human and mouse orthologous genes. **Figure 1B** and Supplementary Table S1 present the top enriched GO terms and pathways for the 382 CR DEGs with human orthologs. Gene set enrichment analysis (GSEA) indicated a dichotomization response in mouse CR microarray gene expression (metabolic and inflammatory) which we previously confirmed by qPCR (15) (**Figure 1C**).

**Figure 1.**
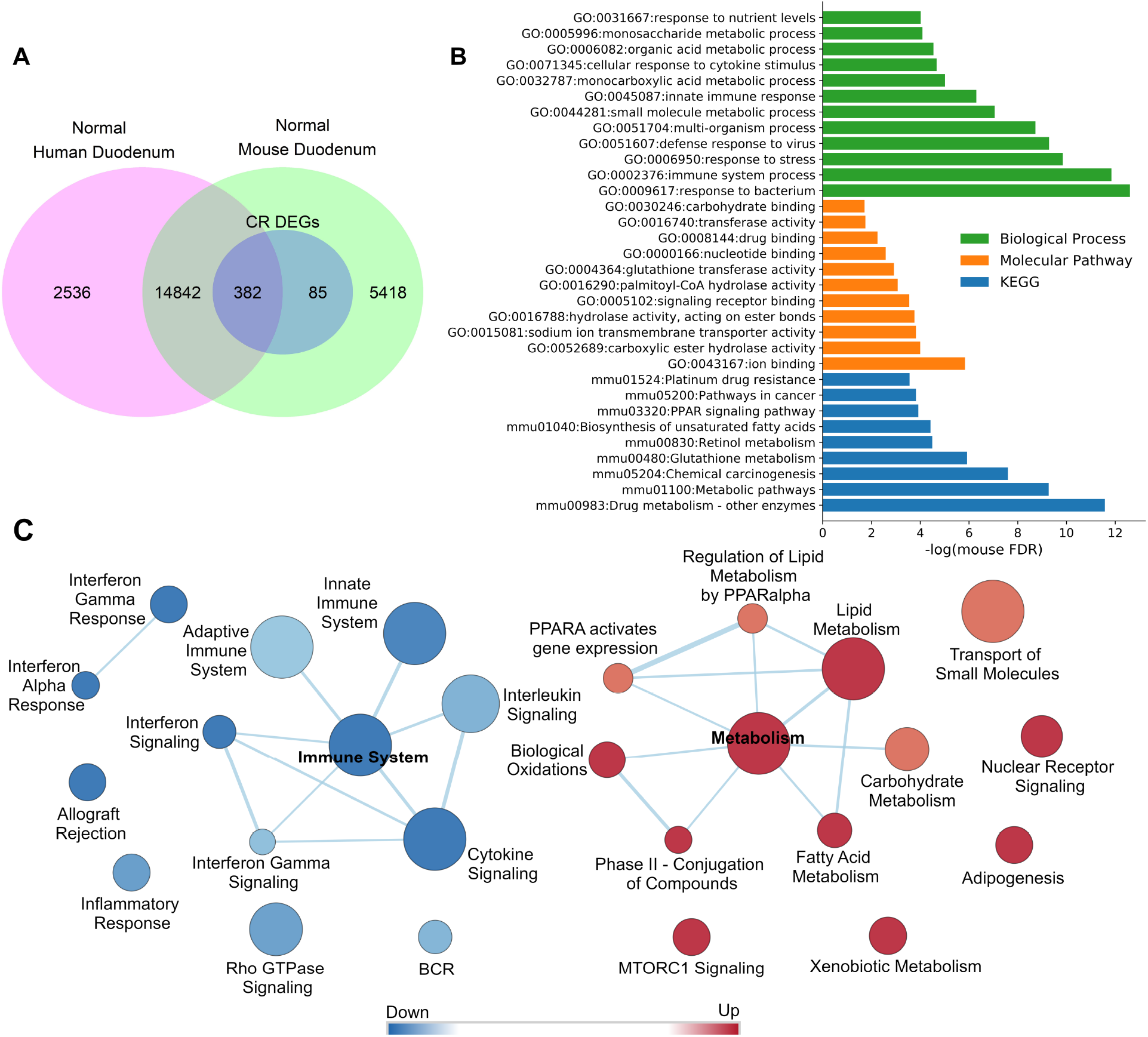
Association between mouse and human ortholog gene sets significantly expressed in normal mouse and normal human DM tissues and GO functional enrichment analysis of the mouse gene subset with human orthologs. **(A)** Venn diagram of mouse and human orthologous gene sets significantly expressed in normal mouse and normal human DM tissues. Normalized log10 transform microarray expression data for mouse and human DM tissues were analyzed (Suppl. Methods). A gene of human or mouse dataset was considered expressed with a mean log signal intensity value larger than cut-off value of three. **(B)** GO functional enrichment analysis of mouse DM DEGs with human orthologs responded to CR (genes selected from mouse significantly expressed gene subset of Panel A) were analyzed using the STRING v11 tools at enrichment FDR<0.05. CR induced DEGs at adj. p-value < 0.05 and |FC| > 1.5. GO analysis of the 382 annotated CR induced mouse DEGs with human orthologs yielded 255 significant Biological Process (BP) terms. Top terms included response to bacterium (GO:0009617, 44 genes), immune system process (GO:0002376, 78 genes), response to stress (GO:0006950, 103 genes), and defense response to virus (GO:0051607, 21 genes). The mouse CR DEGs were enriched in 67 Molecular Function (MF) terms and 34 KEGG Pathways including drug metabolism – other enzymes (mmu00983, 19 genes), metabolic pathways (mmu01100, 60 genes), chemical carcinogenesis (mmu05204, 15 genes), glutathione metabolism (mmu00480, 11 genes), and PPAR signaling pathway (mmu03320 10 genes). The same results with high confidence GO categories were found for the human orthologous genes in the common subset of Venn diagram Panel A. **(C)** Results from GSEA pre-ranked analysis (Suppl. Methods) of mouse CR DEGs (adjusted p-value < 0.05 at |FC| > 1.2). Gene sets (p<0.05, FDR < 0.25) were visualized by Cytoscape Enrichment Map.

### 26% of CR response genes are involved in mucosa normal-adenoma-carcinoma differential gene expression patterns

We carried out a detailed meta-analysis of several datasets including human duodenal adenoma/adenocarcinoma, colorectal adenoma, duodenal cancer (Familial Adenomatous Polyposis cases) transcriptome data, and cancer cell metabolism genes. Our analysis also included APC knockout data, Ingenuity Pathway Analysis (IPA) functional annotations, PubMed and Google literature search and manual literature curation. Supplementary Data 3 and Supplementary Methods describe the full curation processes. **Figure 2A** shows the metadata analysis of duodenal pre-cancerous adenoma/adenocarcinoma patterns as regulated in CR mice. We determined a collection of 121 oncogenes, tumor-associated and cancer-related genes (Supplementary Data 1R) which comprise 26% of our 467 mouse CR DEGs (adj. p-value <0.05 at |FC|>1.5), with 63 upregulated genes and 58 downregulated (Supplementary Data 1S-T). For instance, Supplementary Data 3I-L lists the 20 DEGs genes of CR DM that have been observed in the APC^Min/+^ mouse model of colorectal cancer (28,29). Of these 20 genes, 12 were upregulated (*Mgst2, Aadac, Ppara, Zbtb16, Rdh7, Ugt2b5, Akr1b7, G6pc, Aldh1a1, Mgst1, Ces1d, and Fbp1*) and 8 were downregulated (*Ifit2, Ly6a, Myadm, Cyba, Apobec2, Atf3, Anxa5,* and *Cfi*) in CR mice. These findings suggest a co-regulation pattern of CR DEGs and Apc knockout genes/the APC^Min/+^ mouse model of colorectal cancer (28,29). They are relevant to our hypothesized mucosal response to CR via aberrant tumorigenic regulatory processes.

**Figure 2.**
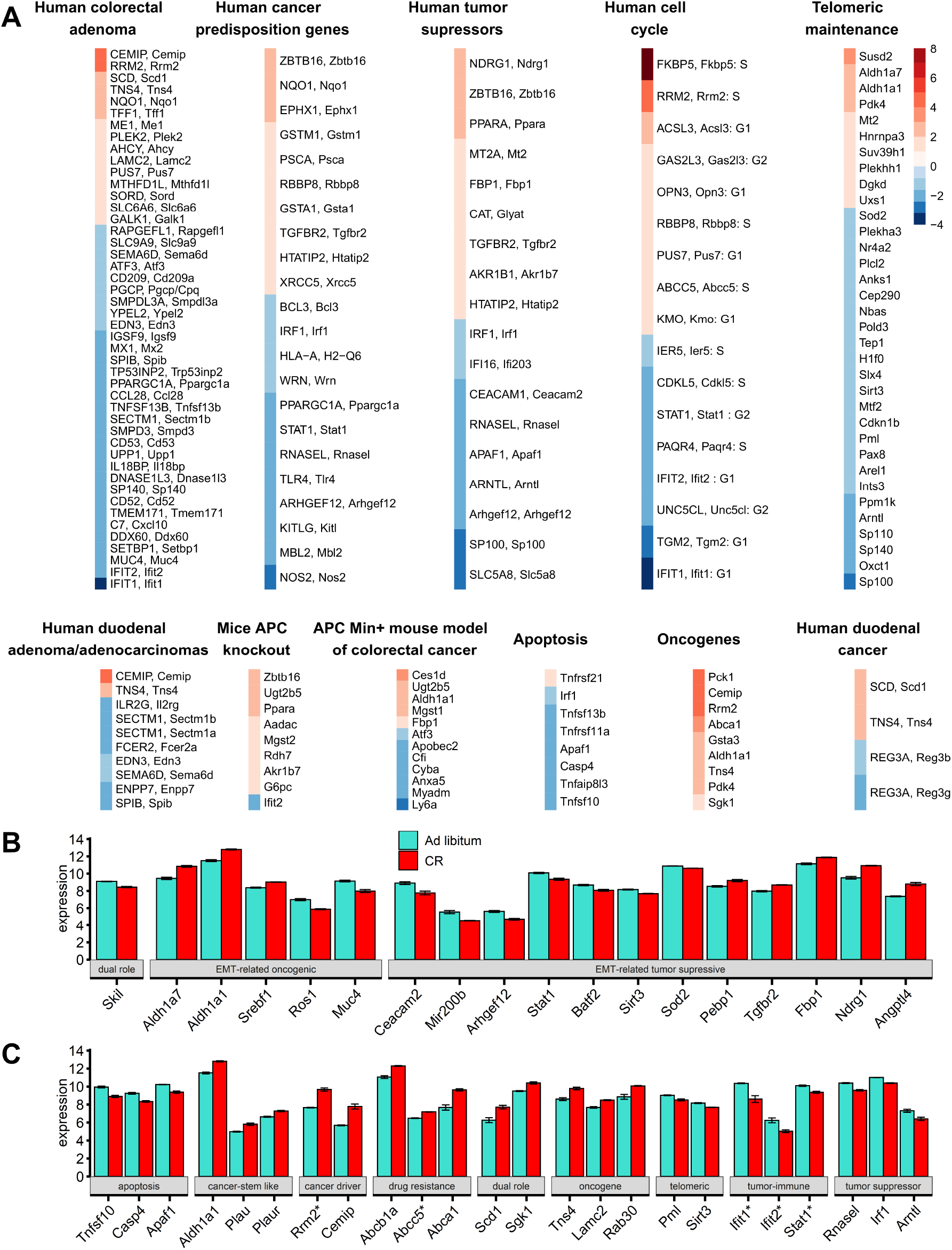
Categorization of CR induced DEG subsets and their heatmaps. **(A)** When human/mouse gene orthologs are both present as labels, this indicates the original meta dataset used was human. The heatmap is colored by the gene expression fold change (FC) in CR DM mice (orange up, blue downregulated gene). Only matching directionality gene expression changes were included when comparing to the 505 CR mice DEGs (upregulated in both mice CR model and human colorectal adenoma, duodenal cancer, and duodenal adenoma/adenocarcinoma or both models downregulated). Supplementary Data 3 lists the exact FC in previous models. Similarly, matched directionality gene expression changes for mice Apc knockout/APC^Min/+^ upregulated and CR mice downregulated or Apc knockout/APC^Min/+^ downregulated and CR upregulated was included. Human tumor suppressors were all suppressed in colon adenocarcinoma. Human cancer predisposition genes, telomeric maintenance, human cell cycle, apoptosis, and oncogene directionality in cancer models is not indicated. **(B)** Data are presented as the mean ± SEM (adjusted p-value < 0.05) for EMT-related CR DEGs with oncogenic, tumor suppressive, or dual roles. **(C)** Selected CR DEGs presented as the mean ± SEM (adj. p-value < 0.05) representing key proliferative, oncogene, tumor suppressor, apoptotic, stem-like epithelial, tumor-immune surveillance, and telomeric genes which may play negative roles in mucosa. *Sgk1* and *Scd1* (CR-upregulated) play dual roles in cancer progression, however, their upregulation promotes tumor growth and migration for colorectal carcinoma (114) and gastric cancer (115). Genes with asterisks indicate cell cycle classification.

Next, we compared CR response genes found in mice DM with the DEGs of normaladenoma-carcinoma patterns (25,30–32). Human duodenal adenoma/adenocarcinoma and colorectal adenoma transcription expression data (orthologs with corresponding mouse CR DEGs matched regulation directionality changes) indicated 53 significant cancer-associated genes of the 467 CR DEGs:15 upregulated genes, 38 suppressed (Supplementary Data 3A-F).

Using the Cancer Predisposition Gene Database (33), 24 human orthologues with corresponding mouse CR DEGs were identified as cancer predisposition genes (CPGs), genes in which inherited mutations in confer increased cancer risk: colon cancer CPGs (*GSTM1, MBL2, RBBP8*) and gastric cancer CPGs (*NOS2, PSCA, TLR4*).

The Tumor Suppressor Gene Database (https://bioinfo.uth.edu/TSGene/) includes 535 human tumor suppressor genes (TSGs) with lower expression in colon adenocarcinoma samples compared to normal tissue samples (34). Eight TSGs were downregulated upon CR (*Rnasel, Irf1, Ifi203, Sp100, Slc5a8, Apaf1, Arntl, Arhgef12*) and nine TSGs were upregulated upon CR (*Ppara, Tgfbr2, Ndrg1, Mt2, Htatip2, Zbtb16, Glyat, Akr1b7, Fbp1*). Using the CheEA3 transcription factor (TF) enrichment analysis tool, we observed that transcription of 7 (*Ppara, Tgfbr2, Ndrg1, Mt2, Htatip2, Akr1b7, Fbp1*) of the 9 up-regulated DEGs genes (called group 1 TSGs) could result in TF binding sites (BS) for liver X receptors (LXRs) (GEO-GSE35262, ChIP-Seq). In contrast, transcription of 6 (*Rnasel, Irf1, Ifi203, Sp100, Slc5a8 and Apaf1*) of the 8 upregulated DEGs (called group 2 TSGs) could result from TF-BS CDS2 interactions binding to promoters of mouse intestinal villus (GEO-GSE34566, ChIP-Seq). Upregulation of group 1 TSGs have alternative functions referring to activation of lipid metabolisms, while inhibition of group 2 TSGs is directly involved in villus morphology, cell organization, and developmental functions. Our DM villus model suggests CR-induced TSG function reduction/loss.

We also asked if there is a risk of highly aggressive and drug-resistant tumors associated with CR-responded DEGs in DM epithelial cells. **Figure 2B** shows that using the dbEMT, an epithelial-mesenchymal transition (EMT) associated gene resource (35), CR DEGs were classified as: oncogenic EMT-related genes (CR upregulated: Aldh1a1, Aldh1a7, Srebf1; CR downregulated: Ros1, Muc4) and tumor-suppressive EMT-related genes (CR downregulated: Ceacam2, Mir200b, Arhgef12, Stat1, Baft2, Sirt3, Sod2; CR upregulated: Angptl4, Ndrg1, Fbp1, Tgfbr2, Pebp1). Skil (suppressed upon CR), was classified as an EMT-related gene with dual roles (oncogenic and tumor-suppressive function). These findings suggest that CR may induce a risk of aberrant EMT to stemness, aggressiveness, and cancer state (36). Outcomes of such balance depend on chromosome instability, including telomeric and cell cycle pathways.

### Telomeric maintenance pathways and cell cycle gene responses in CR mice

Thirty-four telomeric CR mice DEGs (adj. p-value <0.05 at |FC|>1.2) were collected from the databases *TelNet* and IPA, literature searches, and manual curation of TERC, TERRA network, and DNA damage/repair gene sets (Supplementary Data 1Q, Supplementary Data 4, Supplementary Methods). Overall, 70.6% (24/34) telomeric DEGs were suppressed in CR mice, indicating telomeric maintenance dysregulation and chromosomal instability.

To identify CR DEGs directly involved in the cell cycle, we searched for mouse orthologues in CycleBase comprehensive human cell cycle periodic genes lists (Supplementary Table S2). Due to CR in mice, nine DEGs were upregulated (*Fkbp5, Rrm2, Acsl3, Gas2l3, Opn3, Rbbp8, Pus7, Abcc5, Kmo*). Enrichment analysis (STRING v11) showed that Rrm2 and Rbb8 form a functional network with Brca1 and Atm suggesting their involvement in homologous recombination. Additional associations: Rbb8 (G2/M DNA damage checkpoint) and Rrm2 (P53 signalling, glutathione pathways). Eight downregulated cell cycle CR mice DEGs (*Ifit1, Ifit2, Stat1, Tgm2, Unc5cl, Pagr4, Cdkl5, Ier5*) refer to immune system suppression, inflammatory, and immune cells development. Gene associations included positive cellular response to innate immune response (interferon-alpha, interferon beta) (*Ifit1, Ifit2, Stat1;* FDR<0.025) and I-kappa-B kinase/NF-kappa B signalling regulation (*Stat1, Unc5cl, Tgm2;* FDR<0.005), which dysregulation of occurs in chronic inflammatory diseases and certain cancers (37).

**Figure 2C** indicates genes with potential for substantial negative roles in CR DM mucosa.

### Mice mucosa cellular immune system compartments are deeply suppressed by CR and epithelial network interactions are activated

Upon CR, strong/global immune system suppression and epithelial activation occurs. The 171 ISGs consist 37% (171/467) of CR-responding DEGs. *Stat1* (immune) and *Ppara* (metabolic) drivers induce de novo abnormal regulation of DM pathways. The immune and epithelial network (**Figure 3**) contained 118 genes, 311 edges (protein interactions), average number of neighbors 3.64, clustering coefficient 0.14, network density 0.02, and PPI network enrichment p < 1.00 10^−16^. Subcellular ISG and ECG localization networks are in Supplementary Figure S1.

**Figure 3.**
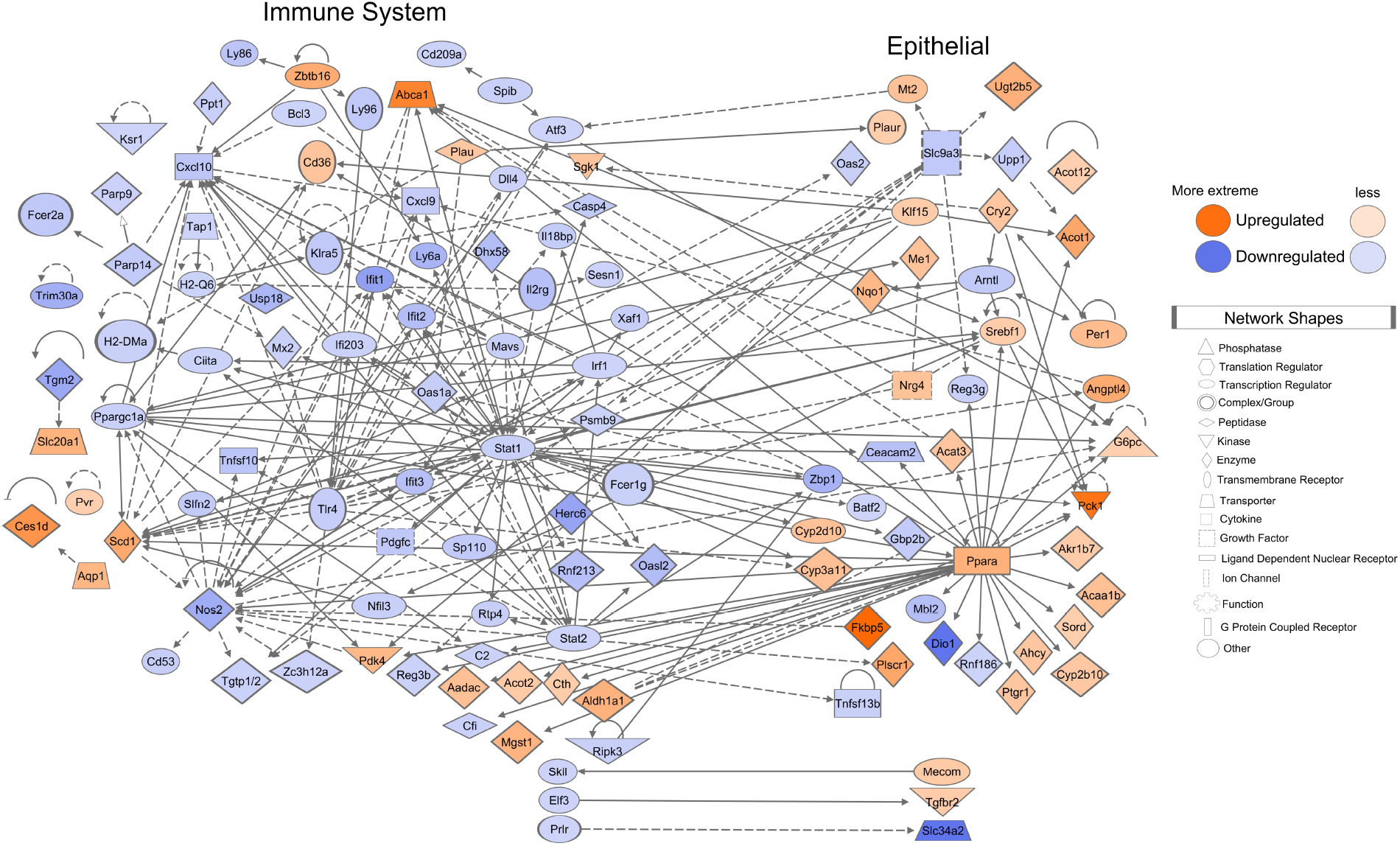
CR suppresses immune system and preferentially activates epithelial cell DEG networks and discriminates the networks interconnection. IPA DB and network graphical tool was used to construct the immune system and epithelial cell-specific subsets DEGs and their interactions. 174 edges (protein interactions) were found among high significantly connected immune genes network interactions with several super hubs (e.g., *Stat1* with 36 edges and *Tlr4* with 19 edges). CR reduces the expression of a vast majority of the immune genes. CR modulates expression of epithelium cells, activating metabolic genes, which had 58 edges within epithelial-specific genes (e.g., *Ppara* pathway directly connected with 26 edges). 34 edges were from immune to epithelium interactions (with hub of *Stat1,* 7 edges), and 45 edges were from epithelium to immune interactions (with hub *Ppara,* 16 edges, mostly linked with the upregulated immune system genes). Blue color node indicates downregulated genes and orange color node indicates upregulated genes. Color intensity is based on the gene expression fold changes. Bold lines indicate direct relationship. Dotted lines indicate indirect relationship.

The suppressed ISGs were characterized using the Disease and Biological Function annotation tool from IPA (Supplementary Table S3). The Lymphoid Tissue Structure and Development categorization (p-value dynamical range: 4.07 10^−19^ to 2.94 10^−3^) included 66 genes within its categories. The functional annotations proliferation of B lymphocytes (p = 9.52 10^−10^, *n* = 18), T cell development (p = 5.12 10^−9^, *n* = 24), development of antigen presenting cells (p = 7.18 10^−7^, *n* = 10), and NK cell proliferation (p = 7.31 10^−6^, *n* = 8), were displayed as a network with 31 genes (**Figure 4A**).

**Figure 4.**
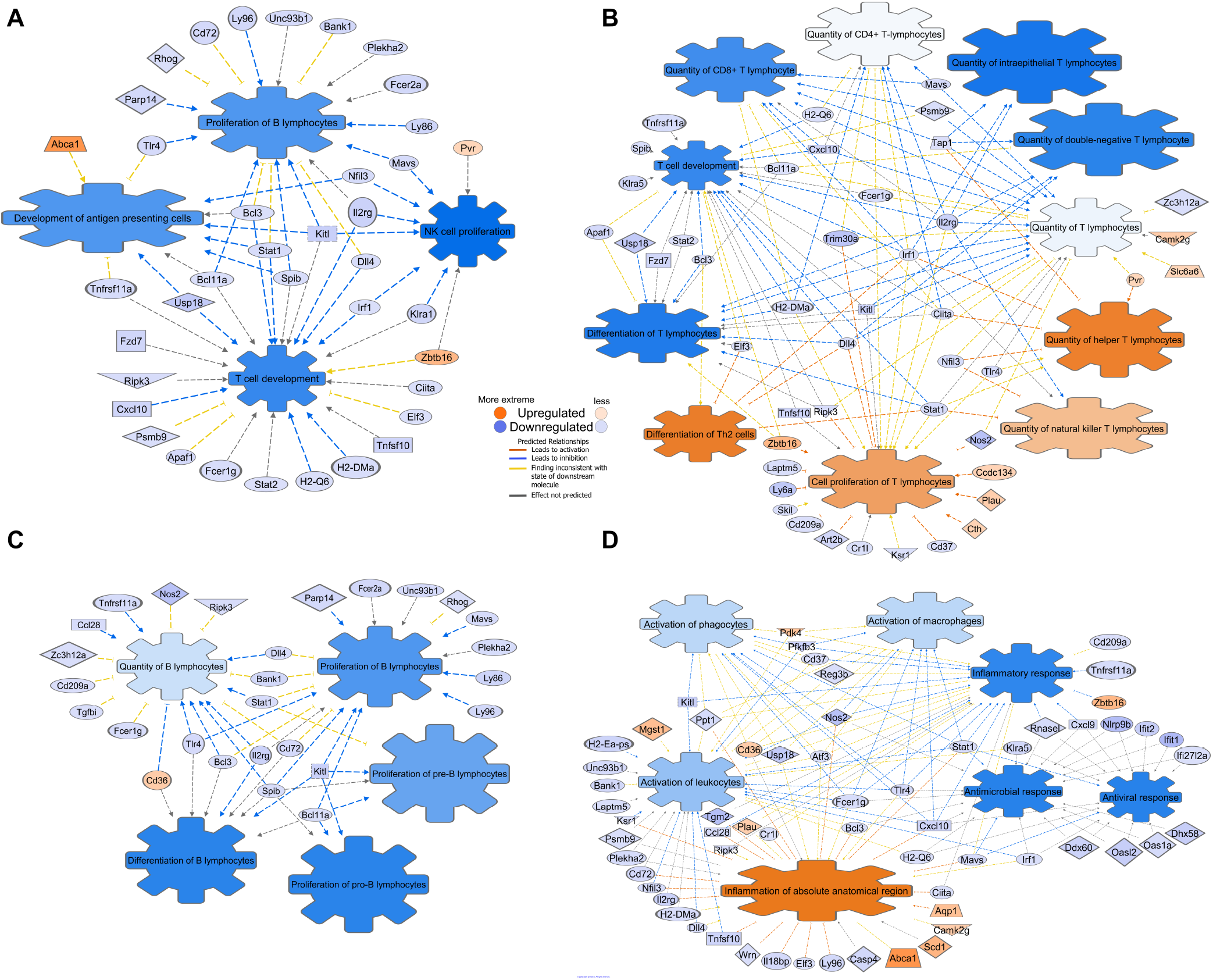
CR reduces T-, B-, NK- and antigen-presenting cells expression, their functional activity and suppresses interconnections between major immune cell type populations. **(A)** IPA network analysis reveals global suppression of the major immune cell types. **(B)** T and NK lymphocytes network. **(C)** B lymphocyte network. **(D)** Inflammatory cells network. The Path Designer in IPA colors the disease/functional annotations and edges based on their current predicted effects; orange is predicted activation, blue is predicted inhibition, yellow is findings inconsistent with state of downstream molecule, and grey color is effect not predicted.

Additionally, 45 T cell-associated functional annotations (**Figure 4B**) and 27 B cell associated functional annotations were displayed (**Figure 4C**). Sixty Inflammatory Response genes (p-value dynamical range: 2.47 10^−16^ to 2.94 10^−3^, n = 77) were selected (**Figure 4D**). An additional suppressed ISG functional annotation network is provided in Supplementary Figure S2. Our analysis determined a deep suppression of cellular immune system genes belonging to all tissue-associated lymphocyte populations and immune system regulatory cells, suggesting systemic immune-cell-specific quantity reduction in the mucosa.

### CR induces downregulation of embryonal/haematopoiesis/immune CSC genes, however upregulation of epithelial cell CSC genes

Cancer stem cells (CSCs) are a subpopulation of cancer cells possessing characteristics associated with normal stem cells, specifically self-renewal and differentiation giving rise to the cell types found in a particular cancer sample (38). CSCs are tumorigenic (tumor-forming), perhaps in contrast to other non-tumorigenic cancer cells. The CSCdb (http://bioinformatics.ustc.edu.cn/cscdb) provides the annotations of 74 marker genes of 40 different CSC lines and 1769 CSC-related genes (38). Supplementary Table S4 indicates 17 CSC-related genes and 1 marker CSC differentially expressed upon CR, with *Aldh1a1* identified as both. Overall, eight CSC DEGs were upregulated including epithelial cell-type associated (*Plau, Plaur, Aldh1a1, Bnip3, Mecom, Mgst1, Abcb1, Ndrg1*), and 10 were downregulated, mostly genes related to embryonic, haematopoiesis, and immune cells (*Ly6a, Kitlg, Il2rg, Dll4, Mecom, C2*).

### Gene ontology and pathway analysis revealed that 26% of CR responded DEGs are involved in malignancy and cancer–associated networks

The 26% (121/467) of DEGs were characterized in IPA (Supplementary Table S5). Within the Cancer category (p-value dynamical range: 7.03 10^−12^ to 2.64 10^−3^, *n* = 48) disease annotations with predicted activation included abdominal carcinoma (p = 7.96 10^−08^, *n* = 12), epithelial neoplasm (p = 2.40 10^−7^, *n* = 20), and tumorigenesis of epithelial neoplasm (p = 1.08 10^−4^, *n* = 13). Annotations with predicted inhibition included abdominal neoplasm (p = 7.03 10^−12^, *n* = 23), digestive organ tumor (p = 1.63 10^−10^, *n* = 21), abdominal cancer (p = 3.00 10^−10^, *n* = 17), and liver cancer (p = 7.57 10^−8^, *n* = 11) (**Figure 5A**). This indicates CR has downstream potential for both pro- and anti-cancerogenic effects. Upregulated pro-oncogenic genes associated with malignant condition and down regulated tumor suppressor genes are listed in Supplementary Table S6. We highlight future qPCR validation targets of CR in cancer-initiation mechanism contexts (Supplementary Table S7). For example, upregulation of CEMIP, RRM2, LAMC2, SCD, TNS4, and PLAU in humans is prognostically unfavorable for various cancers (http://www.proteinatlas.org)(39); these genes have CR-upregulated mouse orthologs.

**Figure 5.**
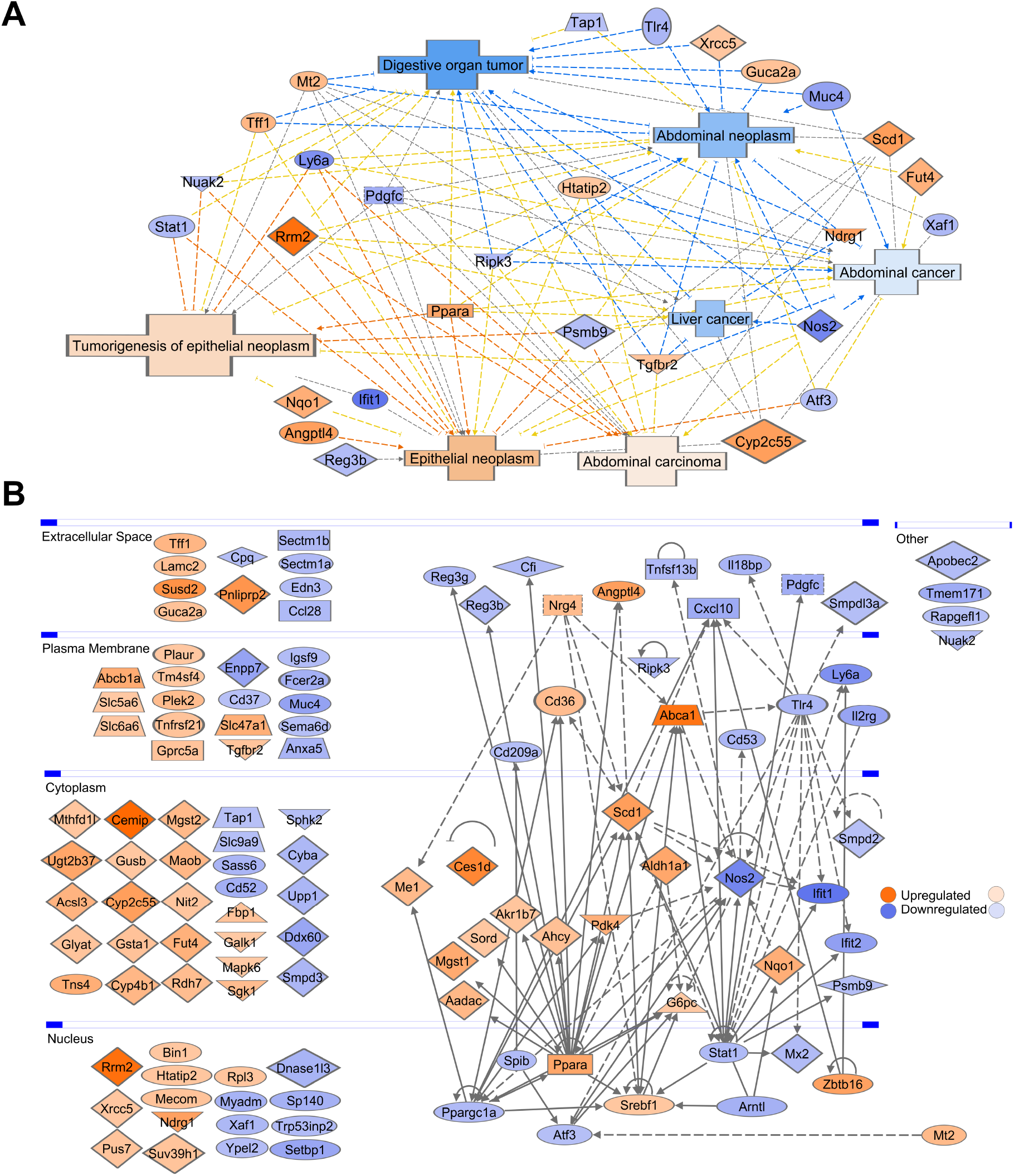
Cancer subcellular network and disease functional annotations. **(A)** A network of 28 genes based on the functional annotations of the cancer-associated genes. **(B)** The network includes 43 connected and 78 non-connected cancer-associated DM genes distributed across four cellular compartments: extracellular space, plasma membrane, cytoplasm, nucleus. Top hubs include *Ppara* (25 downstream targets, 6 upstream targets), *Stat1* (17 downstream, 12 upstream), *Ppargc1a* (12 downstream, 8 upstream), *Tlr4* (12 downstream, 1 upstream), *Scd* (10 downstream, 7 upstream), and *Nos2* (8 downstream, 13 upstream). These hubs account for 54.1% (131/242) of the edge connectivity.

Among the 121 DEGs, 43 genes have subcellular location forming interconnections (**Figure 5B**). The cancer-associated network contained 121 edges, average number of neighbors 3.67, clustering coefficient 0.15, network density 0.05, PPI network enrichment p < 1.00 10^−16^.

### Network of cancer-associated, immune system, epithelial, and telomere gene subsets

Enriched canonical pathways were generated through IPA for each gene subset (Supplementary Table S8). The Sirtuin Signaling Pathway was enriched in: 467 unique DEGs (p = 1.38 10^−3^, *n* = 15), cancer-associated (p = 1.32 10^−6^, *n* = 11), telomeres (p = 9.77 10^−5^, *n* = 5), ECGs (p = 8.51 10^−4^, *n* = 8). We highlight the Sirtuin Signaling Pathway because it had second highest significance for the cancer-associated network canonical pathways (Supplementary Table S9). Transcription of only one of the seven sirtuins, Sirtuin 3, was significantly reduced.

The immune system, cancer-associated, epithelial, and telomere subsets were displayed as a network with the Sirtuin Signaling Pathway overlaid (**Figure 6**). The network has 132 connected components, 227 non-connected components, 350 edges, average number of neighbors 3.74, clustering coefficient 0.14, network density 0.02, and PPI network enrichment p < 1.00 10^−16^. Supplementary Figure S3 shows the network interactions of the tumor-immune and tumor-epithelial microenvironments identifying possible mechanisms of CR-induced malignancy risks or anti-cancerogenic effects.

**Figure 6.**
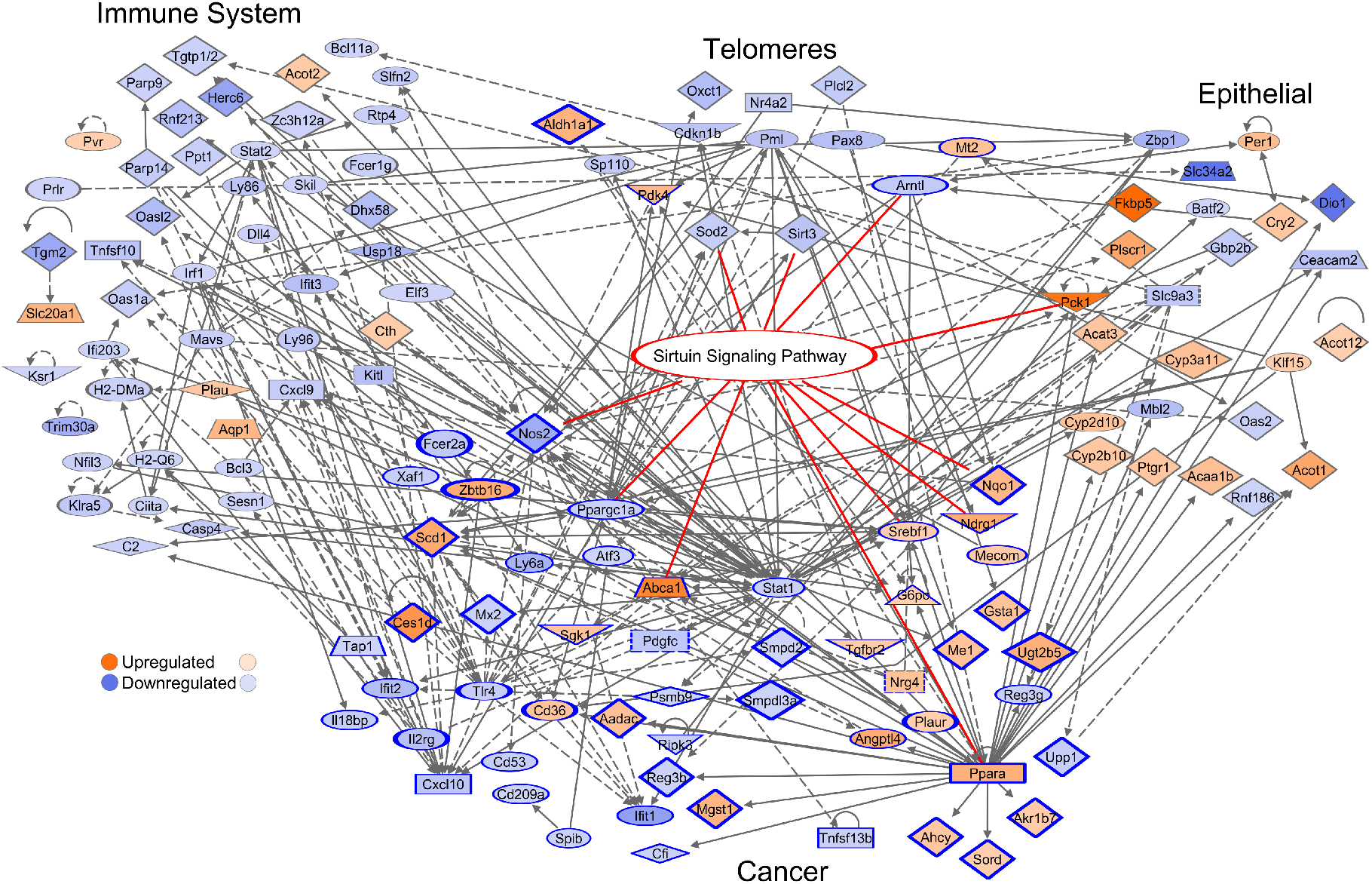
Cross-talk networks with Sirtuin Signaling Pathway. Cancer related genes have dark blue shape outline. Sirtuin Signaling Pathway genes have red edges. All genes are adjusted p-value < 0.05 at |FC| > 1.5, except the telomere subset with |FC| > 1.2. Top hubs include *Stat1* (44 downstream targets, 25 upstream targets), *Ppara* (42 downstream, 7 upstream), *Tlr4* (25 downstream, 4 upstream), *Ppargc1a* (19 downstream, 11 upstream), *Nos2* (14 downstream, 20 upstream), *Pml* (10 downstream, 4 upstream), and *Cxcl10* (1 downstream, 17 upstream). These hubs account for 34.7% (243/700) of the edge connectivity.

### CR induces a network between DEGs of glutathione, chemical carcinogenesis, and sirtuin signaling pathways

Glutathione metabolism (FDR = 1.20 10^−6^, *n* = 11), chemical carcinogenesis (FDR = 2.53 10^−8^, *n* = 15), and sirtuin signaling (p = 1.38 10^−3^, *n* = 15) gene subsets are highly enriched in our DEG dataset (40,41). Consistent with these findings, IPA enrichment analysis revealed significant functional interconnections between DEGs of glutathione, chemical carcinogenesis, and sirtuin signaling pathways (**Figure 7**). With one down-regulated gene exception (*Gpx2,* blue), the interconnected DEGs are upregulated (orange) in glutathione/chemical carcinogenesis pathways. This network model supports SIRT3 as a central coordinator of CR-mediated metabolic reprogramming of DM cellular composition. We included RRM2 in our model, a cell cycle protein (CycleBase-3) involved in the glutathione metabolism pathway (KEGG).

**Figure 7.**
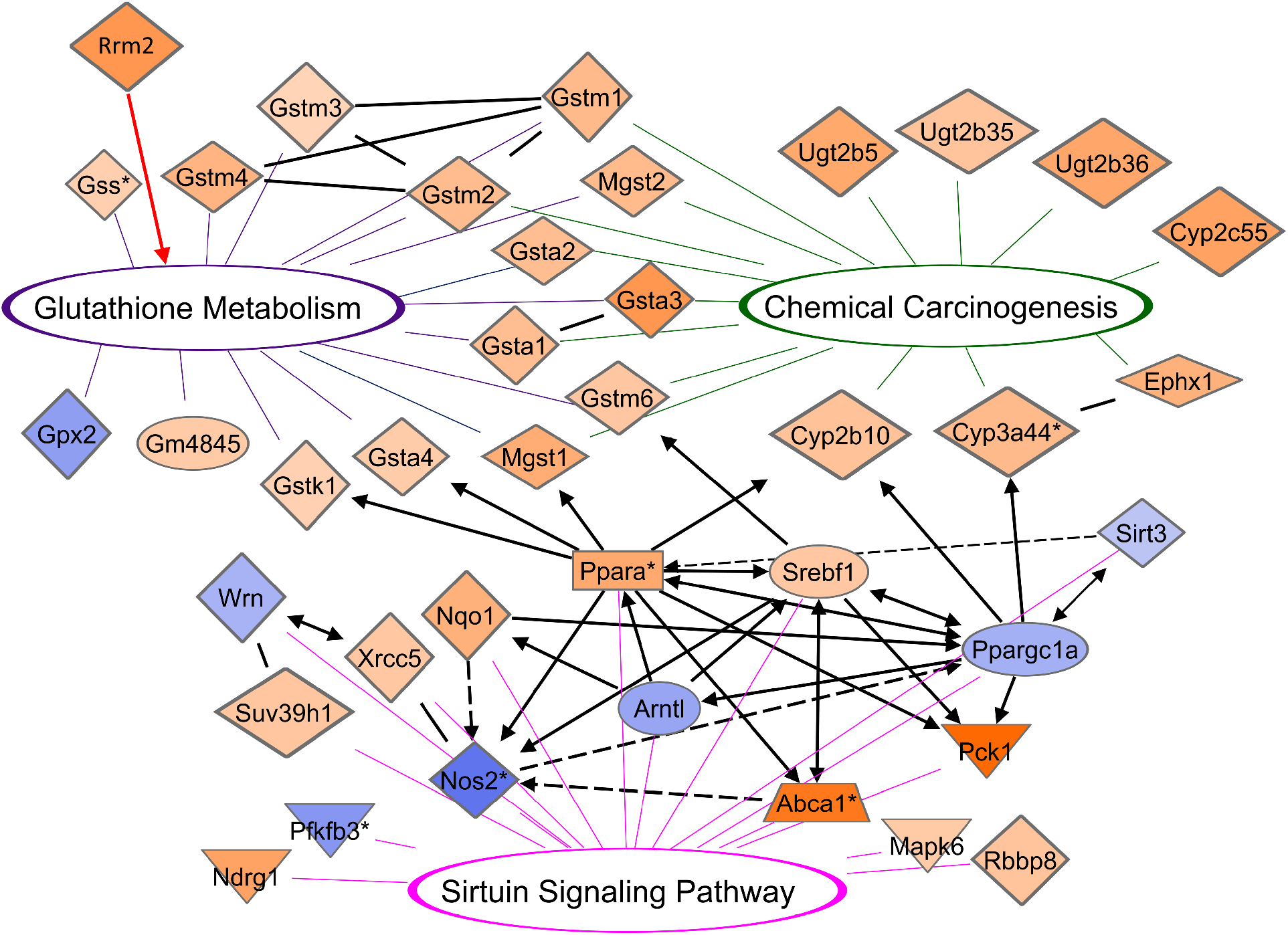
CR induces network between DEGs of glutathione, chemical carcinogenesis, and sirtuin signaling pathways. CR induces network between DEGs of glutathione metabolism (purple; mmu00480), chemical carcinogenesis (green; mmu05204), and sirtuin signaling pathways (magenta). Sirtuin signaling regulation has downstream targets in glutathione metabolism (*Gstk1, Gsta4, Mgst1, Gstm6*) and chemical carcinogenesis (*Cyp2b10, Cyp3a44*). Black lines indicate molecule relationships. Self-loops were removed for easier visualization of network interactions between molecules. Asterisks on molecules indicates druggable targets: *Ppara* (aleglitazar, atorvastatin/choline fenofibrate, bezafibrate, clofibrate, docosahexaenoic acid, ezetimibe/fenofibrate/simvastatin, gemfibrozil, NS-220, pemafibrate, TPST-1120, tesaglitazar), *Nos2* (GW 273629, N(G)-monomethyl-D-arginine, pimagedine, triflusal), *Cyp3a44* (atazanavir/cobicistat/darunavir), *Abca1* (probucol), *Gss* (N-acetyl-L-cysteine), *Pfkfb3* (PFK-158). All genes are adjusted p-value < 0.05 at |FC| > 1.5, except Sirt3 with FC-1.38.

### Interferon-inducible DNA-binding gene family members on Chr 11 are CR-suppressed

One of the top CR DEGs, predicted/uncharacterized protein-coding gene 5431 (*Gm5431*) localized in Chr 11qB(1.2), was strongly suppressed (FC =-6.76; p = 2.7 10^−17^). We observed that the gene belongs to Chr11qB(1.2) locus containing a family of paralog genes that encode interferon (IFN)-inducible GTPase proteins, also called immunity-related p47 GTPases (INF-I GTPases). Interferons induce intra-cellular programs functional in innate and adaptive immunity against infectious pathogens. Chr 11qB(1.2) locus comprises *Gm5431* with genes *Irgm1, Tgtp1, Tgtp2, Ifi47* and uncharacterized genes *9930111J21Rik1, 9930111J21Rik2, Gm12185,* Gm12186 and Gm12187 (and their transcribed isoforms) (**Figure 8A**, Supplementary Data 5). Nine of the ten genes are localized on the chromosome negative strand. Genes in this locus have high evolutionarily conserved sequences and paralogous in several other loci (for instances, Chr 11qB(1.3) (Irgm2, Igtp), Chr18 (Iigp1, F830016B08Rik)), in other chromosomes and orthologous in genomes of many species (**Figure 8A**, Supplementary Figure S4). IRGM in humans is an ortholog to this family, with links to Chron’s disease, inflammatory disorders, and non-alcoholic fatty liver disease (42). *Irgc1* (with RefSeq transcript NM_199013) located on Chr 7 is also an ortholog of the gene family to the human genome (IRGC) but its transcription was not significantly regulated by CR (Supplementary Figure S4).

**Figure 8.**
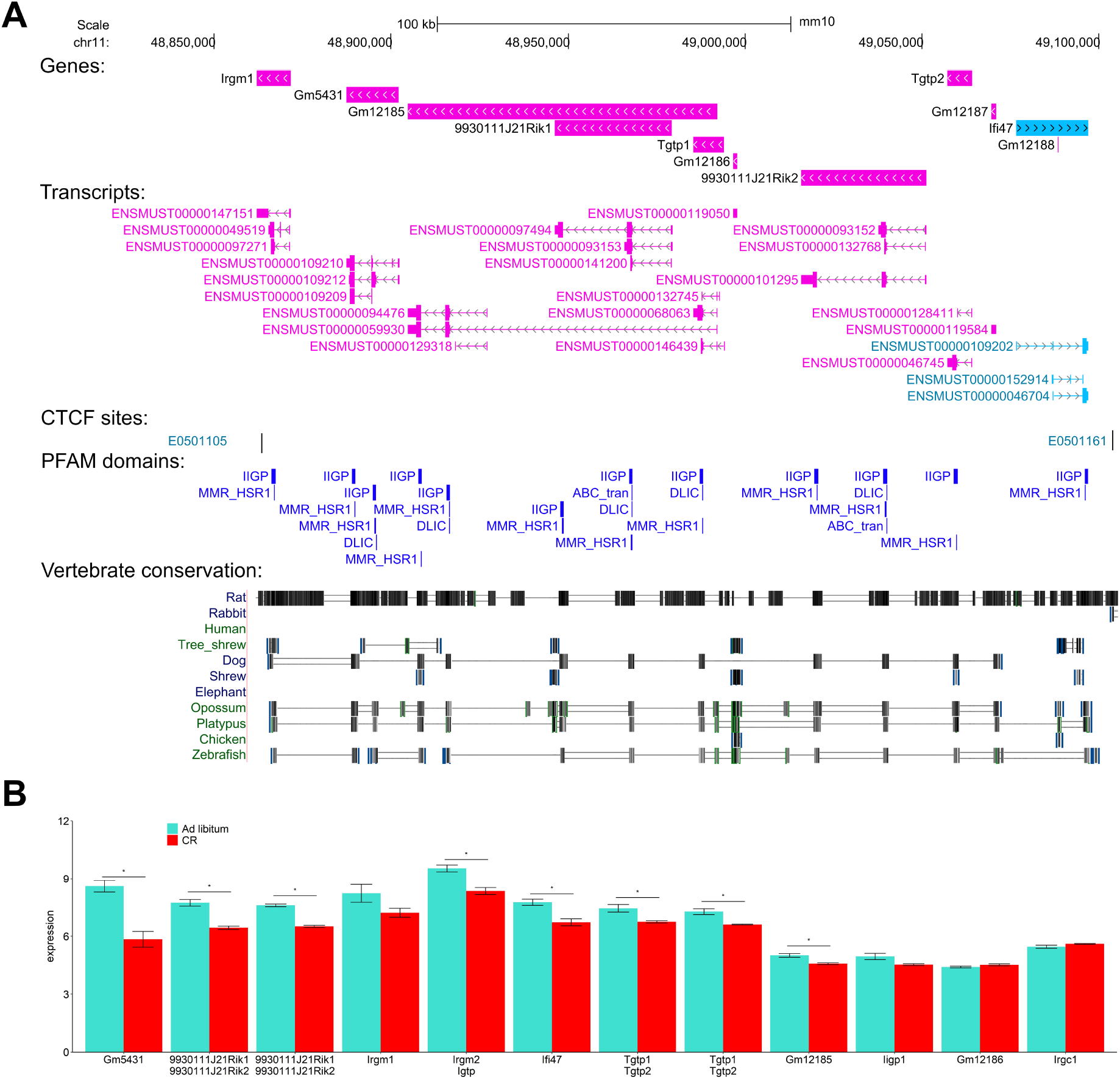
Chr11qB(1.2) locus comprising 10 genes belonging to the CR IFN-induced family. **(A)** Genes and their transcribed isoforms are annotated by the latest Ensembl (Build 75) currently supported by the UCSC genome browser (Build 100 annotations in Supplementary Data 5). All PFAM annotated proteins have one or more conserved functional domains. Pfam-A annotates transcript regions within the peptide recognizable as Pfam protein domains in characterized and non-characterized protein-coding genes for mouse assembly mm10 track of UCSC by the software HMMER3: IIGP (Interferon-inducible GTPase; PF05049), MMR_HSR1 (50S ribosome-binding GTPase; PF01926), DLIC (Dynein light intermediate chain; PF05783), ABC_tran (ABC transporter; PF00005). CTCF sites are displayed by the ENCODE Candidate Cis-Regulatory Elements (cCREs) track combined from all cell types. We used the UCSC “Multiz alignments” track, mouse assembly mm10, with multiple alignment data combining PhyloP and PhastCons methods. It shows evolutionary conservation from the PHAST package for 60 vertebrates and three subsets (Glires, Euarchontoglires, placental mammal). **(B)** Data are presented as the mean ± SEM. Asterisks indicate adj. p-values < 0.05 for statistical significance.

Using MAFFT-DASH - multiple alignment software integrated with structural search (43), we found high confidence similar sequences and domains between IIGP1 and all our proteins (Supplementary Data 5). Longer proteins encoded by *Gm5431, 9930111J21Rik1,* and *9930111J21Rik2* show the highest sequence similarity with IIGP1/1TQ4 and include two nonoverlapping IIGP1 sequence homologous. IIGP1, the member of INF-I GTPases encoded by gene *Iigp1* localized on Chr18, has known crystal structure and atom resolution 3D model 1TQ4. The IIGP1/1TQ4 is built of two domains, the G domain and a helical domain including five DNA-binding motifs (44). Due to high sequence similarity and presence of evolutionarily conserved sequence domains (**Figure 8A**), we suggest structural and functional similarity of IIGP1 and the members of the INF-I GTPase family analyzed in this section.

The transcription levels of the IFN-I-GTPases were commonly suppressed due to CR response (**Figure 8B**). Furthermore, the Chr. 11qB(1.2) loci is flanked by CTCF binding sites. Analysis of the IFN-inducible gene family members suggests that CR in DM induces global suppression of interferon-induced cellular functions in innate and adaptive immunity.

### Drug targets in energy-responsive metabolic pathways

A total of 36 drug targets were associated with the 467 mouse duodenum CR DEGs as identified by IPA (Supplementary Data 6). PPARa, NOS2, TLR4, and CXCL10 are the most relevant strongest hubs with the highest connectivity determined by network analysis. TLR4 is targetable by eritoran, GSK1795091, OM 174 lipid, and resatorvid. The activity of CXCL10 is modulated by MDX-1100. Anti-CXCL10 monoclonal antibodies have implications in infectious disease, chronic inflammatory, and autoimmune disease therapeutics and attenuate murine inflammatory bowel disease and murine AIDS colitis (45). Agents that deplete cellular glutathione metabolism in combination with arsenic trioxide are studied for the treatment of non-acute promyelocytic leukemia (46). Attempts to increase the efficacy of cancer treatments via lifestyle or diet changes would benefit from further examination of these top targets in nutrient-responsive pathways.

## Discussion

Because pre-malignant and cancer cells must reprogram their metabolic state in every step of progression to survive in nutrient-altered conditions, metabolic reprogramming is recognized as a cancer hallmark (47). In our study, we for the first time identified cancer driver genes and genes playing key roles in oncogenic pathways induced by CR perturbation of cellular composition, tissue homeostasis and metabolic reprogramming, leading to cell proliferation of epithelium mucosa and global depletion of immune system control.

We found that 26% of the CR responding differential expressed genes (DEGs) in mice DM consist of cancer-associated genes—most never studied in CR contexts. These responses may lead to perturbation of telomere maintaining mechanisms, and activation of the cell cycle, pro-oncogenic, and stem-like cancer cell pathways including EMT. CR-induces metabolic reprogramming, which affects the ISGs, consisting of 37% of the total DEGs; the majority of ISGs are suppressed, including cell-autonomous immunity and tumor immune evasion controls.

### Major findings of this study

**1)** CR induces tissue homeostasis dysregulation, shifting the transcriptome profile to pro-oncogenomic pathway patterns in DM associated with activation of metabolism and proliferative activity of <u>epithelial cells</u>, but depletion of intracellular immunity and functions of all <u>immune cell types</u>. **2)** CR induces transcription of <u>cell cycle</u> genes including a subset of key cancer-associated signaling genes directly involved in reprogramming pathways, premalignancy, malignancy states, and poor outcome in mice and humans. **3)** In CR response, <u>apoptotic</u> gene expression is reduced or not significantly varied. **4**) Tissue-specific proliferative epithelial stem DEGs are <u>activated</u>, however, immune-specific stem cell progenitor genes are <u>suppressed</u> upon CR. **5)** CR induces transcription activation of key tumor-susceptible and oncogenes gene sets and their networks with tissue-specific risk of carcinogenesis and down-regulates protective mechanisms mediated by key tumor suppressor genes. **6)** CR induces multiple transcription suppression effects in autophagy, tumor-immune surveillance mechanics and genes associated with induction and effector stages of NK, T-, B-cells and macrophages immune response. **7)** Detoxifying exogenous chemicals enzymes and drug metabolism networks with glutathione pathways are activated upon CR, but could be involved in anti-cancer and pro-cancer outcomes via mutagenesis and DNA damage/repair mechanisms.

Our network and pathway analysis provide interconnections of CR DEGs with biological functions, molecular mechanisms, cell types and cellular compartments. **Figure 9** shows the two-compartmental response models which hypothesise possible dysregulation mechanics of tissue/cell homeostasis upon CR in intestinal mucosa tissue leading to pro-oncogene pathway activation and increased potential risk of premalignant and cancer states. Supplementary Figure S5 shows key pathways and gene targets that may be involved in the multiple disbalances increasing risk of metabolic reprogramming leading to malignant states. Table 1 shows the selected list of the CR-response genes that expression is strongly supported by literature data as key genes of oncogenesis, immune suppression, and metabolic reprogramming and could be considered and tested as CR-induced cancer risk markers.

**Figure 9.**
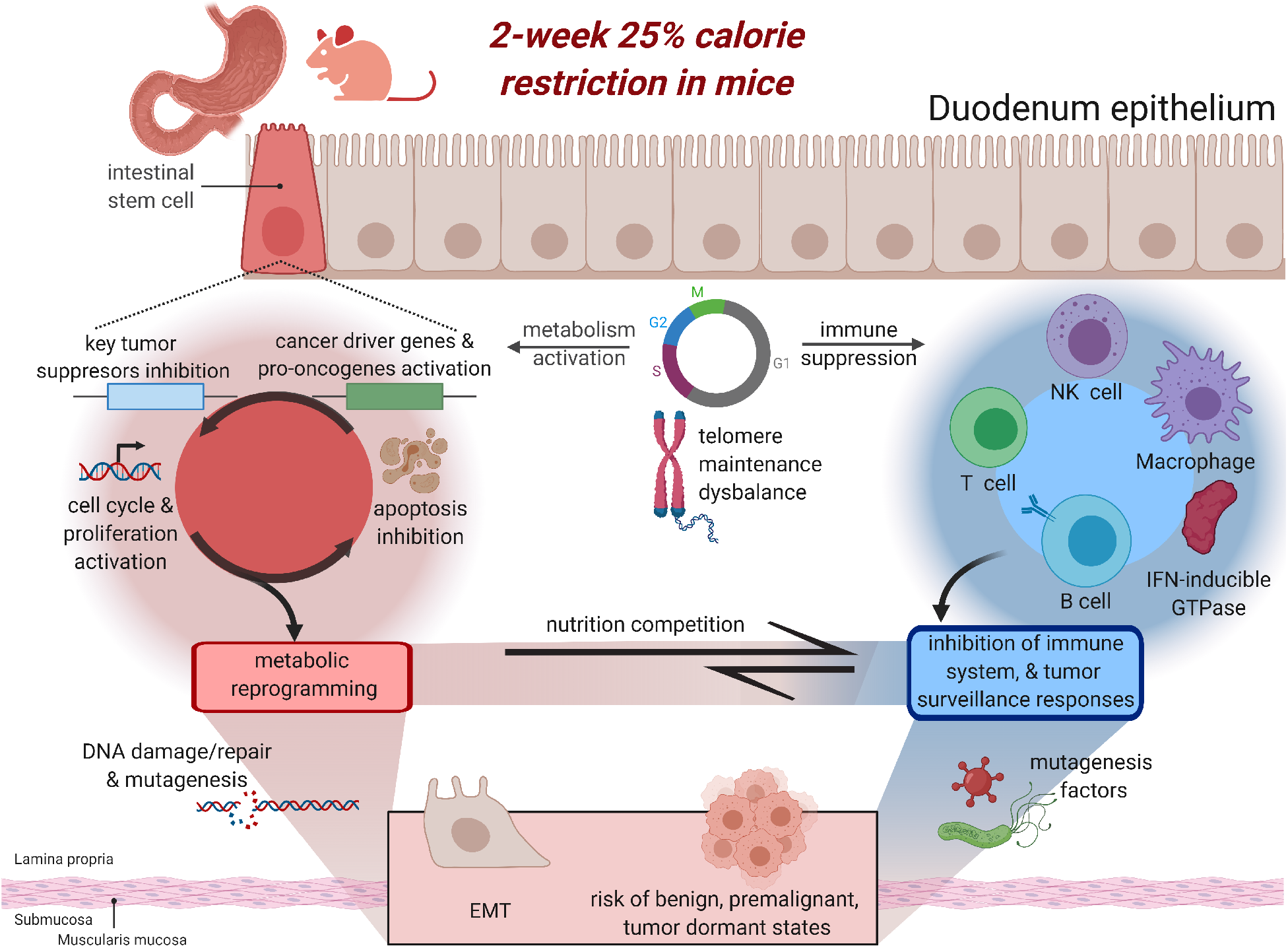
CR cellular disbalance models through immune and autophagy suppression. CR-induced metabolic reprogramming and immune suppression. DEGs and pathways analyses strongly support our hypothesis that CR dramatically changes the epithelial versus immune cells relationships. Functional interactions modulating the homeostasis cooperation between cell types may increase malignancy risk. Morphologically, mucosa enterocyte differentiation and their cell type density depends on the cell position along both the vertical axis (crypt-villus) and horizontal axis (proximal to distal) of the GI tract. At steady-state in normal conditions (homeostasis) of mucosa tissue, the cell types at differentiation states are position-dependent and relationships between cell types remains balanced. Our results suggest that CR induces a hypocellular and multiple cell-types disbalance by stimulation of cell proliferation and autonomation of this process. Our model suggests that CR response leads to metabolic activation, inducing cell cycle genes and proliferation of epithelial stem cells. CR reduces the total mass of epithelial cells (vertical axis) (21), autophagy, apoptosis mechanisms, and telomere stability. Immune cell populations including progenitors and effector cells are globally reduced, suggesting increased pathogen susceptibility. If the processes controlling DNA and RNA damage induced by chemical carcinogens (49,64) become dysregulated, it increases the risk of transformation and uncontrolled proliferation of abnormal stem-like cells.

**Table 1.**
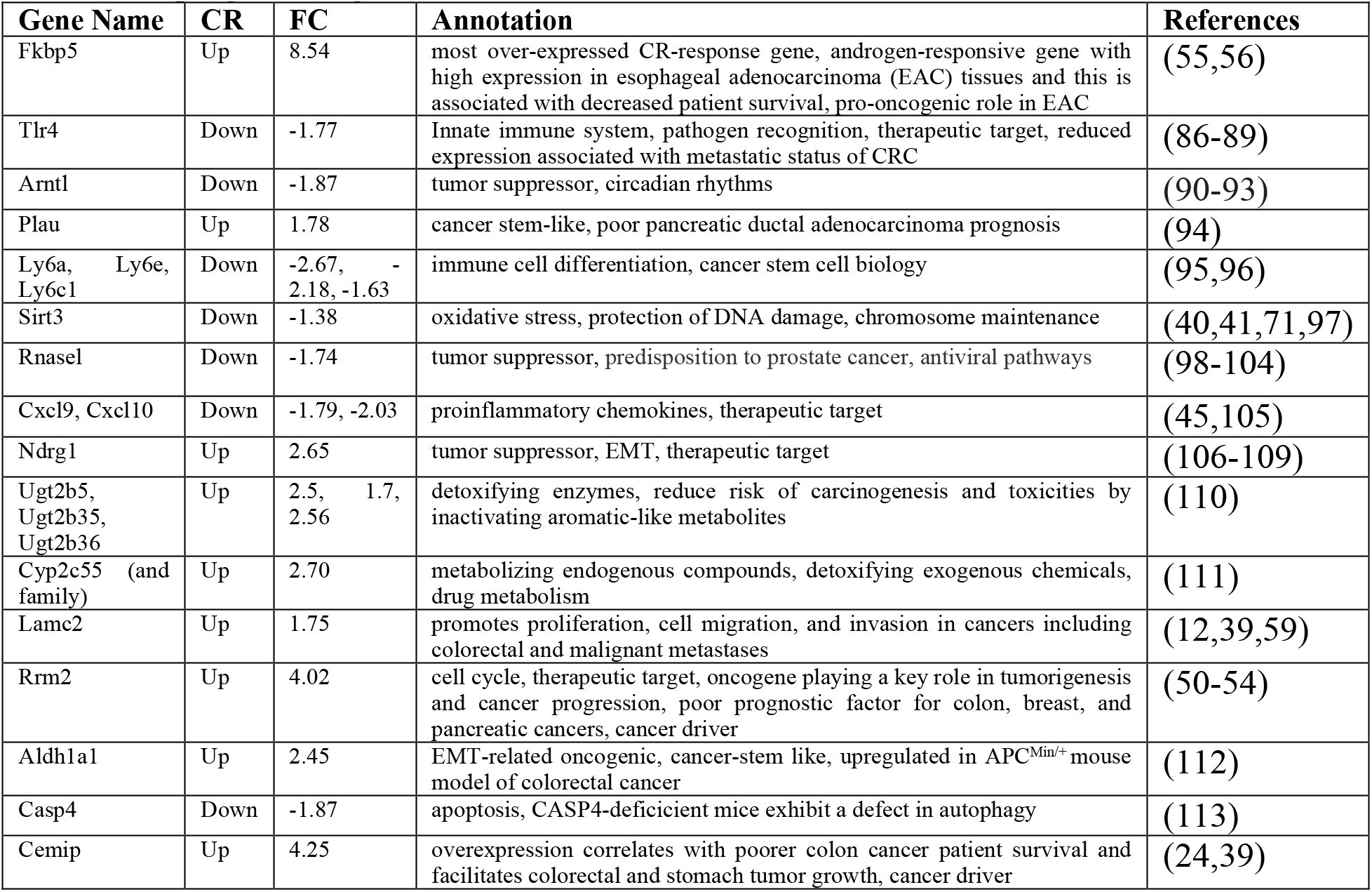
Selected CR DEGs in tumor suppressive, oncogenic, immune, epithelial stem-like, anticancer, and detoxifying pathways for future validation and consideration in pre-cancer and metabolic reprogramming states.

According to our model, long-term CR may confer increased tumorigenic risks if there are durable and sustained increases in stem mutations and abnormal proliferation, DNA damage, epigenetic modification, and cell numbers that can undergo mutagenesis. Activated stem cells of DM villi may exhibit reprogramming behavior if metabolic machinery is used at higher rates, in combination with aggressive chemical and pathogenic bacterial factors. While external to our analysis, bacteria (*H. Pylori*), parasites, and viruses may provide the mutagenesis factors initiating oncogenesis in mucosa epithelial cells. Pathogenic microbiota signals could also act on the intestinal stem cell niche and homeostasis upon CR (48).

The multiple gene dysregulation in mucosa upon CR response could be interpreted as an “early ischemia”-like syndrome observed in newborns, demonstrating marked reduction of villous height and increasing epithelium crypt-villus ratio (in the vertical axis) in combination with small numbers of immune cells in intestinal lymphoid structures along the lamina propria (horizontal axis) (49). These changes may resemble those seen in autolysis. Reduction or no change in apoptosis gene expression in epithelial cells occurs and no acute inflammatory cells are activated (49). This coincides with global INF-induced autophagy reduction in our study.

### CR induces epithelial cell cycle proliferation and suppresses apoptosis, increasing cancer risk

We observed suppressed apoptotic genes upon CR. Fasting, short-term, and long-term dietary restriction almost uniformly reduces cellular proliferation (liver, bladder, skin, heart, colorectum) explaining anticarcinogenic effects of CR (5). A potential pro-cancerogenic effect in fasted-refed animals resulted from proliferation increases with apoptosis decreases in response to refeeding (5). CR in some experimental models enhances cell death rate, however, proliferation/ apoptosis rates vary through the course of treatment (5,11,21). CR enhances the proliferation of Lgr5+ intestinal stem cells, the cell-of-origin for intestinal precancerous adenomas and leads to the first organoid and acceleration of the second organoid formations (11,21).

In normal physiological conditions, cell cycle and apoptosis gene activity in tissue-specific stem cells, progenitor, and differentiated epithelial cells maintains tissue homeostasis. However, short-term CR disturbs this balance and may switch-on pro-oncogenic pathways. For instance, *Rrm2* is high upregulated in DM CR response. RRM2 is an oncogene playing a key role in tumorigenesis and cancer progression, including colorectal and oesophageal cancers (oncoMX.org)(50–54). CR-activated cell cycle periodic genes in our study are directly or indirectly involved in tumorigenesis pathways, drive cancer cell proliferation and aggressiveness (RRM2, ACSL3, RBBP8, KMO, FKBP5), drug resistance (ABCC5) and modulate chemosensitivity (FKBP5) (references in Supplementary Table S2A). *Fkbp5* is the most over-expressed CR-response gene. This androgen-responsive gene has high expression in esophageal adenocarcinoma (EAC) tissues and is associated with decreased patient survival (55). FK506-binding protein 5 (FKBP5) plays a pro-oncogenic role in EAC (55,56). In tissue resection specimens from EAC patients, FKBP5 expression was positively associated with proliferation as measured by Ki-67 expression, which was also observed in the prostate cancer weight loss clinical trials (12,56). FKBP5 is a cis-trans prolyl isomerase that binds to the immunosuppressants FK506 and rapamycin. The number of activated CR mice cell cycle DEGs and their expression level increments (fold change) were stronger in the upregulated genes vs down-regulated DEGs. Our model showed high upregulation of FKBP5 and RRM2 (FC=8.5 and FC=4.0 respectively) compared to downregulated TGM2 and PAQR4 (FC=3.1 and FC=1.76 respectively). *Tgm2* and *Paqr4* may provide suppressive effects in some aggressive cancer cells (Supplementary Table S2B)(57,58). TGM2-siRNA knockdown attenuated colorectal cancer cell growth through the wnt3a/β-catenin/cyclin D1 pathway (58).

The pioneering NCI-funded (R21 CA161263) randomized clinical trial of prostate cancer patients undergoing presurgical CR-mediated body mass intervention manifested significantly grated Ki67 proliferation rate results (12). It determined activated DEGs related to tumor cell signaling associated with increased proliferation, transcription, oncogenesis, migration and invasion: *CBLC, POLB, ATF1, TFEB, ACVR1B, LAMC2, GSK3B, PHF6, MAP3K8, ARIDIA.* Our study also demonstrated CR-induced *Lamc2* overexpression. LAMC2 promotes proliferation, cell migration, and invasion in cancers including colorectal and malignant metastases (12,59). LAMC2 has been suggested as a therapeutic target because of its association with the Wnt/betta-catenin signaling pathway and effects on PI3K and ACT (12,59).

Our model includes potential gene regulatory switches modulating anti-cancer to pro-oncogenic CR response. CR induces specific mechanisms involved in pathogenic or adaptative EMT-driven stem cell response and proliferation of mucosa epithelial cells. The EMT pathway gene expression alteration (36) may or may not indicate markers solely, but silencing/dormant states could be formed. Due to CR-induced immune system depletion and mutagenesis/genotoxic events (leading to occurrence and accumulation of a driver mutation and genome instability), the risks of oncogene activation and tumor suppressor depletion required for conversion to tumor-initiating cells over long periods could increase (60). This suggests that tumor lesions may need to be studied in experimental systems over time with lower threshold detection level of small benign polyps and dormant malignant states (8). Time course systems biology analyses of CR severity and length should be conducted on pre-existing tumor dormant states (5,11).

### Oncogenes and pro-oncogenic genes are poorly studied in CR and carcinogenesis contexts

Searching in PubMed revealed only 32.2% (39/121) of our CR cancer-associated genes (Supplementary Data 7) are associated with terms “caloric” or “calorie” or “caloric restriction” or “calorie restriction” or “dietary restriction”. Supplementary Table S6 includes upregulated oncogenes upon CR: *Cemip, Tns4, Aldh1a1, Rab30, Rrm2*, and *Gsta3*. CEMIP mRNA overexpression correlates with poorer colon cancer patient survival and facilitates colorectal and stomach tumor growth (OncoMX) (24). TNS4 expression is transcriptionally regulated by MAP kinase signaling pathway and plays a critical role in tumorigenesis in several tissues including the colon. TNS4 and ALDH3A1 expression levels were increased in HCT-8 colon cancer cells and influence cancer cell migration, invasion, and proliferation (61). RAS oncogene family member 30 (RAB30) was upregulated in the microarray of epithelial colorectal adenocarcinoma and acute lymphoblastic leukemia-derived cell lines (62). PPARα-sensitive genes during starvation include *Cxcl10* and *Rab30* (63). *RRM2* plays oncogenic roles in tumorigenesis (50) and is a poor prognostic factor for colon, breast, and pancreatic cancers (51–54).

### Glutathione (GSH) plays a dual role in cancer progression

The Glutathione S-transferases (GSTs), phase II detoxification enzymes, were activated in CR mice: membrane-bound microsomal (*Mgst1, Mgst2*) and cytosolic family members (*Gsta1, Gsta2, Gsta3, Gsta4, Gstm1, Gstm2, Gstm3, Gstm4, Gstm6, Gstk1*). In healthy cells, it is crucial for removal and detoxification of carcinogens (64). However, GSH metabolism is associated with colorectal cancer pathogenesis (65). The GSH system regulates proliferation and survival by offering redox stability in a variety of cancers (66,67). MGST1 is crucial for stem differentiation (68). *MGST1* polymorphisms may contribute to CRC risk (69). GST-overexpressing phenotypes are present in many drug-resistant tumors (including breast, colon, and lung cancers) (66). We propose therapies targeting the GSH antioxidant system in tumors in combination with CR may sensitize cancer cells to treatment (64) and reduce the risk of cancerous effects in CR conditions. Mice treated with L-buthionine-sulfoximine depleted GSH levels in esophageal cancer and decreased tumor burden, inhibited cell proliferation, and activated cell apoptosis (70).

### The Sirtuin Signaling Pathway mediates CR-responded cancer-associated networks

This pathway regulates cancer cell metabolic reprogramming in glucose-poor environments (71). Pck1, a CR upregulated Sirtuin Signaling Pathway gene, is associated with cancer cell gluconeogenesis and increased PCK1 expression is crucial for cancer growth in the absence of glucose (72). In our study, upon CR in mice DM, expression of *Sirt3* isoforms (probe set 10568997, NM_022433) was downregulated. Sirtuin 3 (SIRT3) is a major NAD(+)-dependent mitochondrial deacetylase and the key regulator of fundamental processes frequently dysregulated in aging and other diseases (40,41). New experimental designs may be pivotal to validate SIRT3 as a cancer regulator in CR.

### CR induces multiple transcription suppression effects in the genes of autophagy and tumor immune surveillance mechanics

In contrast to activation of the metabolic genes associated with epithelial cells, we found immune system transcription suppression in 37% of DEGs induced by CR. Restricted cell cycle genes in interferon-inducible pathways include antiviral and anti-cancer activity (73) (*Ifit1* and *Ifit2*) and pro-inflammatory reactions recruiting immune cells to target cells (74) (genes *Stat1* and *Unc5cl*) (Supplementary Table S2B).

### Negative role of CR in tumor-immune surveillance

The immune system classified CR-responding DEGs were majorly suppressed (Supplementary Table S10). An intact immune system is essential to prevent neoplastic cell development and progression (75,76). Mice with a homozygous deletion of the Rag-2 alleles completely lack NK-like T, T and B cells and have increased incidence and growth of spontaneous tumors and chemically induced cancer lesions (77). IFN-g forms the basis of an extrinsic tumor-suppressor mechanism in immunocompetent hosts (75,76,78). STAT1 can suppress tumor formation (75). The elimination of interferon IFN-g or STAT1 genes in mice resulted in increased incidence and growth of spontaneous and chemically induced tumors (75,77,78). CR may impact IFN-mediated tumor elimination (79).

### CR induces suppression of INF pathways in DM

Type II interferon signaling (IFN-g) is suppressed by CR. IFN-inducible GTPases play central roles in defending the mammalian cell’s interior against a diverse group of invading pathogens. The IFN-inducible GTPases ability to control infection at the level of an individual cell—a process termed cell-autonomous immunity— is responsible for microbial killing (80). We identified IFN-inducible GTPase genes on Chr11q B(1.2) which were downregulated upon CR. *Ifit1, Ifit2, Ifit3, Ifi203, Ifi44, Ifi27l2a,* and *Irf1* (all CR-suppressed) function in antiviral and antimicrobial response. Ifit1 knockout promoted viral replication in murine norovirus infected cells (73). Loss of interferon regulatory factor 1 (IRF1) function causes severe susceptibility to infections in mice and humans (81). IRF1 suppresses tumor cell growth, stimulating immune responses against tumor cells (81–84). Defects in *IRF1* are associated with gastric and lung cancer, and myelogenous leukemia (82). The *Stat1/2* and *Irf1/9* CR-network interactions mediate immunity and gut inflammation (74).

Further discussion of key modulators of CR-response in metabolic reprogramming and immune suppression in Table 1 can be found in the Suppl. Discussion including: tumor suppressor Rnasel (CR-downregulated), proinflammatory chemokines Cxcl9/10 (CR-downregulated), immune cell differentiation Ly6 gene family (CR-downregulated), predicted therapeutic target Ndrg1 (CR-upregulated), and detoxification enzymes (CR-upregulated).

In summary, our results suggest that CR-mediated metabolic reprogramming suppresses multiple host tumor surveillance prevention mechanics including cell-autonomous immunity and activates signaling pathways of pro-oncogenes, tissue-specific cycling and silenced stem cells, and cancer predisposition genes driving pre-malignant and cancer states. This study’s limitations refer to meta-data sets available for analysis, data type (e.g., microarrays), DM expression profile composition (cellular mixture), CR experimental time design, and pre-clinical models of CR effects on tumor pathobiology. However, our systems biology, data-driven, and hypothesis-testing approach to metadata provides feasibility by identifying DM DEGs subsets according to cell-type specificity functions: IFN-inducible GTPases genes, CR-induced cancer- and immune system cross-talks. This provides multiple determinants of aberrant signalling, cell-cell interaction networks leading dynamically to cell transformation, clonal selection, immune response delay, immune surveillance suppression and tumorigenesis (85).

These findings may change the paradigm regarding the anti-cancer role of CR and initiate new unbiased CR cancer biology studies. Clearly, no single cell type/subtype or unique pathway accounts for all anti-cancer or pro-oncogenic effects of CR in normal, but complex mucosa tissue. As with most chronic disease intervention strategies, combination approaches improving lifestyle (CR, diet, physical activity), prophylactic strategies and reproducible pharmacological interventions that target specific cells and their pathways are needed to prevent pre-cancer and cancer states. Future (bioengineering) directions to implement our results is to identify novel drugs and CR mimetics, compounds that mimic the specific CR mechanism. This will allow protection against cancerous metabolic tissue reprogramming pathways, chromosome stability, provide normal stem cell differentiation and proliferation, ensure physiologic cellular composition balances in target the tissue, and protect the immune system from suppression or hyper-activation.

Our computationally predicted approaches provide new platforms and resources for the formulation and analysis of testable hypothesis for CR in pro-cancer and cancer prevention mechanisms. Our models could be useful for further modelling and experimental validations of CR-response protecting metabolic reprogramming pathways. Further study is encouraging for the perspective utilization of CR-associated mechanisms in clinical oncology strategies.

## Conclusion

- CR dramatically reduces immune responses, dysregulates tissue-specific epithelial stem cell developmental processes and telomere maintaining processes
- CR induces metabolic reprogramming processes, oncogenic and cell cycle pathways, and may increase the risk of malignancy
- CR-induced Rrm2, Lamc2, Fkbp5, and aberrant glutathione gene family activation coupled with Sirtuin3 and RNaseL suppression could play tumorigenic roles in mucosa pathophysiology
- Interferon-inducible gene family including novel members on Chr11qB1.2 are suppressed by CR

## Supporting information

Supplementary Methods and Discussion

## Data Availability

The microarray datasets analyzed during the current study are available in the ArrayExpress repository, https://www.ebi.ac.uk/arrayexpress/experiments/E-MTAB-6248.

## Authors’ contributions

Concept, design and study supervision: V.A.K., Development of methodology, analysis and interpretation of results and writing of manuscript: E.M, V.A.K. Critical review and discussion of the manuscript, generation of biological samples: K.D.

## Acknowledgements

Anonymous referees for their careful review, constructive criticism and useful comments. BioRender for Figure 9.

## References

1. Colman, R.J., Anderson, R.M., Johnson, S.C., Kastman, E.K., Kosmatka, K.J., Beasley, T.M., Allison, D.B., Cruzen, C., Simmons, H.A., Kemnitz, J.W. et al. (2009) Caloric restriction delays disease onset and mortality in rhesus monkeys. Science, 325, 201–204.

2. Hursting, S.D., Smith, S.M., Lashinger, L.M., Harvey, A.E. and Perkins, S.N. (2010) Calories and carcinogenesis: lessons learned from 30 years of calorie restriction research. Carcinogenesis, 31, 83–89.

3. Kritchevsky, D. (2001) Caloric restriction and cancer. J Nutr Sci Vitaminol (Tokyo), 47, 13–19.

4. Hursting, S.D., Dunlap, S.M., Ford, N.A., Hursting, M.J. and Lashinger, L.M. (2013) Calorie restriction and cancer prevention: a mechanistic perspective. Cancer Metab, 1, 10.

5. Klebanov, S. (2007) Can short-term dietary restriction and fasting have a long-term anticarcinogenic effect? Interdiscip Top Gerontol, 35, 176–192.

6. Harvie, M. and Howell, A. (2012) Energy restriction and the prevention of breast cancer. Proc Nutr Soc, 71, 263–275.

7. O’Flanagan, C.H., Smith, L.A., McDonell, S.B. and Hursting, S.D. (2017) When less may be more: calorie restriction and response to cancer therapy. BMC Med, 15, 106.

8. Kakuni, M., Morimura, K., Wanibuchi, H., Ogawa, M., Min, W., Hayashi, S. and Fukushima, S. (2002) Food restriction inhibits the growth of intestinal polyps in multiple intestinal neoplasia mouse. Jpn J Cancer Res, 93, 236–241.

9. Tsao, J.L., Dudley, S., Kwok, B., Nickel, A.E., Laird, P.W., Siegmund, K.D., Liskay, R.M. and Shibata, D. (2002) Diet, cancer and aging in DNA mismatch repair deficient mice. Carcinogenesis, 23, 1807–1810.

10. Pugh, T.D., Oberley, T.D. and Weindruch, R. (1999) Dietary intervention at middle age: caloric restriction but not dehydroepiandrosterone sulfate increases lifespan and lifetime cancer incidence in mice. Cancer Res, 59, 1642–1648.

11. Mihaylova, M.M., Sabatini, D.M. and Yilmaz, O.H. (2014) Dietary and metabolic control of stem cell function in physiology and cancer. Cell Stem Cell, 14, 292–305.

12. Demark-Wahnefried, W., Rais-Bahrami, S., Desmond, R.A., Gordetsky, J.B., Hunter, G.R., Yang, E.S., Azrad, M., Fruge, A.D., Tsuruta, Y., Norian, L.A. et al. (2017) Presurgical weight loss affects tumour traits and circulating biomarkers in men with prostate cancer. Br J Cancer, 117, 1303–1313.

13. Kristal, A.R., Blount, P.L., Schenk, J.M., Sanchez, C.A., Rabinovitch, P.S., Odze, R.D., Standley, J., Vaughan, T.L. and Reid, B.J. (2005) Low-fat, high fruit and vegetable diets and weight loss do not affect biomarkers of cellular proliferation in Barrett esophagus. Cancer Epidemiol Biomarkers Prev, 14, 2377–2383.

14. Demark-Wahnefried, W., Rogers, L.Q., Gibson, J.T., Harada, S., Fruge, A.D., Oster, R.A., Grizzle, W.E., Norian, L.A., Yang, E.S., Della Manna, D. et al. (2020) Randomized trial of weight loss in primary breast cancer: Impact on body composition, circulating biomarkers and tumor characteristics. Int J Cancer, 146, 2784–2796.

15. Duszka, K., Ellero-Simatos, S., Ow, G.S., Defernez, M., Paramalingam, E., Tett, A., Ying, S., Konig, J., Narbad, A., Kuznetsov, V.A. et al. (2018) Complementary intestinal mucosa and microbiota responses to caloric restriction. Sci Rep, 8, 11338.

16. Hammer, S.T. and Greenson, J.K. (2013) The clinical significance of duodenal lymphocytosis with normal villus architecture. Arch Pathol Lab Med, 137, 1216–1219.

17. Marsh, M.N. and Rostami, K. (2016) What Is A Normal Intestinal Mucosa? Gastroenterology, 151, 784–788.

18. Rostami, K., Marsh, M.N., Johnson, M.W., Mohaghegh, H., Heal, C., Holmes, G., Ensari, A., Aldulaimi, D., Bancel, B., Bassotti, G. et al. (2017) ROC-king onwards: intraepithelial lymphocyte counts, distribution & role in coeliac disease mucosal interpretation. Gut, 66, 2080–2086.

19. Yousefi, M., Nakauka-Ddamba, A., Berry, C.T., Li, N., Schoenberger, J., Simeonov, K.P., Cedeno, R.J., Yu, Z. and Lengner, C.J. (2018) Calorie Restriction Governs Intestinal Epithelial Regeneration through Cell-Autonomous Regulation of mTORC1 in Reserve Stem Cells. Stem Cell Reports, 10, 703–711.

20. Pena-Villalobos, I., Casanova-Maldonado, I., Lois, P., Sabat, P. and Palma, V. (2018) Adaptive Physiological and Morphological Adjustments Mediated by Intestinal Stem Cells in Response to Food Availability in Mice. Front Physiol, 9, 1821.

21. Yilmaz, O.H., Katajisto, P., Lamming, D.W., Gultekin, Y., Bauer-Rowe, K.E., Sengupta, S., Birsoy, K., Dursun, A., Yilmaz, V.O., Selig, M. et al. (2012) mTORC1 in the Paneth cell niche couples intestinal stem-cell function to calorie intake. Nature, 486, 490–495.

22. Huels, D.J., Bruens, L., Hodder, M.C., Cammareri, P., Campbell, A.D., Ridgway, R.A., Gay, D.M., Solar-Abboud, M., Faller, W.J., Nixon, C. et al. (2018) Wnt ligands influence tumour initiation by controlling the number of intestinal stem cells. Nat Commun, 9, 1132.

23. Bultman, S.J. (2017) Interplay between diet, gut microbiota, epigenetic events, and colorectal cancer. Mol Nutr Food Res, 61.

24. Fink, S.P., Myeroff, L.L., Kariv, R., Platzer, P., Xin, B., Mikkola, D., Lawrence, E., Morris, N., Nosrati, A., Willson, J.K. et al. (2015) Induction of KIAA1199/CEMIP is associated with colon cancer phenotype and poor patient survival. Oncotarget, 6, 30500–30515.

25. Thiruvengadam, S.S., O’Malley, M., LaGuardia, L., Lopez, R., Wang, Z., Shadrach, B.L., Chen, Y., Li, C., Veigl, M.L., Barnholtz-Sloan, J.S. et al. (2019) Gene Expression Changes Accompanying the Duodenal Adenoma-Carcinoma Sequence in Familial Adenomatous Polyposis. Clin Transl Gastroenterol, 10, e00053.

26. Kramer, A., Green, J., Pollard, J., Jr. and Tugendreich, S. (2014) Causal analysis approaches in Ingenuity Pathway Analysis. Bioinformatics, 30, 523–530.

27. Lyons, Y.A., Wu, S.Y., Overwijk, W.W., Baggerly, K.A. and Sood, A.K. (2017) Immune cell profiling in cancer: molecular approaches to cell-specific identification. NPJ Precis Oncol, 1, 26.

28. Paoni, N.F., Feldman, M.W., Gutierrez, L.S., Ploplis, V.A. and Castellino, F.J. (2003) Transcriptional profiling of the transition from normal intestinal epithelia to adenomas and carcinomas in the APCMin/+ mouse. Physiol Genomics, 15, 228–235.

29. Sansom, O.J., Reed, K.R., Hayes, A.J., Ireland, H., Brinkmann, H., Newton, I.P., Batlle, E., Simon-Assmann, P., Clevers, H., Nathke, I.S. et al. (2004) Loss of Apc in vivo immediately perturbs Wnt signaling, differentiation, and migration. Genes Dev, 18, 1385–1390.

30. Sakaguchi, Y., Yamamichi, N., Tomida, S., Takeuchi, C., Kageyama-Yahara, N., Takahashi, Y., Shiogama, K., Inada, K.I., Ichinose, M., Fujishiro, M. et al. (2018) Identification of marker genes and pathways specific to precancerous duodenal adenomas and early stage adenocarcinomas. J Gastroenterol.

31. Pesson, M., Volant, A., Uguen, A., Trillet, K., De La Grange, P., Aubry, M., Daoulas, M., Robaszkiewicz, M., Le Gac, G., Morel, A. et al. (2014) A gene expression and pre-mRNA splicing signature that marks the adenoma-adenocarcinoma progression in colorectal cancer. PLoS One, 9, e87761.

32. Sabates-Bellver, J., Van der Flier, L.G., de Palo, M., Cattaneo, E., Maake, C., Rehrauer, H., Laczko, E., Kurowski, M.A., Bujnicki, J.M., Menigatti, M. et al. (2007) Transcriptome profile of human colorectal adenomas. Mol Cancer Res, 5, 1263–1275.

33. Wei, R., Yao, Y., Yang, W., Zheng, C.H., Zhao, M. and Xia, J. (2016) dbCPG: A web resource for cancer predisposition genes. Oncotarget, 7, 37803–37811.

34. Zhao, M., Kim, P., Mitra, R., Zhao, J. and Zhao, Z. (2016) TSGene 2.0: an updated literature-based knowledgebase for tumor suppressor genes. Nucleic Acids Res, 44, D1023–1031.

35. Zhao, M., Liu, Y., Zheng, C. and Qu, H. (2019) dbEMT 2.0: An updated database for epithelial-mesenchymal transition genes with experimentally verified information and precalculated regulation information for cancer metastasis. J Genet Genomics, 46, 595–597.

36. Nieto, M.A., Huang, R.Y., Jackson, R.A. and Thiery, J.P. (2016) Emt: 2016. Cell, 166, 21–45.

37. Hoesel, B. and Schmid, J.A. (2013) The complexity of NF-kappaB signaling in inflammation and cancer. Mol Cancer, 12, 86.

38. Shen, Y., Yao, H., Li, A. and Wang, M. (2016) CSCdb: a cancer stem cells portal for markers, related genes and functional information. Database (Oxford), 2016.

39. Uhlen, M., Zhang, C., Lee, S., Sjostedt, E., Fagerberg, L., Bidkhori, G., Benfeitas, R., Arif, M., Liu, Z., Edfors, F. et al. (2017) A pathology atlas of the human cancer transcriptome. Science, 357.

40. Haigis, M.C., Deng, C.X., Finley, L.W., Kim, H.S. and Gius, D. (2012) SIRT3 is a mitochondrial tumor suppressor: a scientific tale that connects aberrant cellular ROS, the Warburg effect, and carcinogenesis. Cancer Res, 72, 2468–2472.

41. Chen, Y., Fu, L.L., Wen, X., Wang, X.Y., Liu, J., Cheng, Y. and Huang, J. (2014) Sirtuin-3 (SIRT3), a therapeutic target with oncogenic and tumor-suppressive function in cancer. Cell Death Dis, 5, e1047.

42. Mehto, S., Jena, K.K., Nath, P., Chauhan, S., Kolapalli, S.P., Das, S.K., Sahoo, P.K., Jain, A., Taylor, G.A. and Chauhan, S. (2019) The Crohn’s Disease Risk Factor IRGM Limits NLRP3 Inflammasome Activation by Impeding Its Assembly and by Mediating Its Selective Autophagy. Mol Cell, 73, 429–445 e427.

43. Rozewicki, J., Li, S., Amada, K.M., Standley, D.M. and Katoh, K. (2019) MAFFT-DASH: integrated protein sequence and structural alignment. Nucleic Acids Res, 47, W5–W10.

44. Ghosh, A., Uthaiah, R., Howard, J., Herrmann, C. and Wolf, E. (2004) Crystal structure of IIGP1: a paradigm for interferon-inducible p47 resistance GTPases. Mol Cell, 15, 727–739.

45. Liu, M., Guo, S., Hibbert, J.M., Jain, V., Singh, N., Wilson, N.O. and Stiles, J.K. (2011) CXCL10/IP-10 in infectious diseases pathogenesis and potential therapeutic implications. Cytokine Growth Factor Rev, 22, 121–130.

46. Lee, C., Lin, Y., Huang, M., Lin, C., Liu, C., Chow, J. and Liu, H.E. (2006) Increased cellular glutathione and protection by bone marrow stromal cells account for the resistance of non-acute promylocytic leukemia acute myeloid leukemia cells to arsenic trioxide in vivo. Leuk Lymphoma, 47, 521–529.

47. Ohshima, K. and Morii, E. (2021) Metabolic Reprogramming of Cancer Cells during Tumor Progression and Metastasis. Metabolites, 11.

48. Xing, P.Y., Pettersson, S. and Kundu, P. (2020) Microbial Metabolites and Intestinal Stem Cells Tune Intestinal Homeostasis. Proteomics, 20, e1800419.

49. Glickman, J.N. (2018) The Nonneoplastic Small Intestine.

50. Furuta, E., Okuda, H., Kobayashi, A. and Watabe, K. (2010) Metabolic genes in cancer: their roles in tumor progression and clinical implications. Biochim Biophys Acta, 1805, 141–152.

51. Liu, X., Zhang, H., Lai, L., Wang, X., Loera, S., Xue, L., He, H., Zhang, K., Hu, S., Huang, Y. et al. (2013) Ribonucleotide reductase small subunit M2 serves as a prognostic biomarker and predicts poor survival of colorectal cancers. Clin Sci (Lond), 124, 567–578.

52. Fujita, H., Ohuchida, K., Mizumoto, K., Itaba, S., Ito, T., Nakata, K., Yu, J., Kayashima, T., Souzaki, R., Tajiri, T. et al. (2010) Gene expression levels as predictive markers of outcome in pancreatic cancer after gemcitabine-based adjuvant chemotherapy. Neoplasia, 12, 807–817.

53. Hsieh, Y.Y., Chou, C.J., Lo, H.L. and Yang, P.M. (2016) Repositioning of a cyclin-dependent kinase inhibitor GW8510 as a ribonucleotide reductase M2 inhibitor to treat human colorectal cancer. Cell Death Discov, 2, 16027.

54. Zhang, C., Aldrees, M., Arif, M., Li, X., Mardinoglu, A. and Aziz, M.A. (2019) Elucidating the Reprograming of Colorectal Cancer Metabolism Using Genome-Scale Metabolic Modeling. Front Oncol, 9, 681.

55. Smith, E., Palethorpe, H.M., Ruszkiewicz, A.R., Edwards, S., Leach, D.A., Underwood, T.J., Need, E.F. and Drew, P.A. (2016) Androgen Receptor and Androgen-Responsive Gene FKBP5 Are Independent Prognostic Indicators for Esophageal Adenocarcinoma. Dig Dis Sci, 61, 433–443.

56. Palethorpe, H.M., Drew, P.A. and Smith, E. (2017) Androgen Signaling in Esophageal Adenocarcinoma Cell Lines In Vitro. Dig Dis Sci, 62, 3402–3414.

57. Feng, Y., Sun, T., Yu, Y., Gao, Y., Wang, X. and Chen, Z. (2018) MicroRNA-370 inhibits the proliferation, invasion and EMT of gastric cancer cells by directly targeting PAQR4. J Pharmacol Sci, 138, 96–106.

58. Gu, C., Cai, J., Xu, Z., Zhou, S., Ye, L., Yan, Q., Zhang, Y., Fang, Y., Liu, Y., Tu, C. et al. (2019) MiR-532-3p suppresses colorectal cancer progression by disrupting the ETS1/TGM2 axis-mediated Wnt/beta-catenin signaling. Cell Death Dis, 10, 739.

59. Huang, D., Du, C., Ji, D., Xi, J. and Gu, J. (2017) Overexpression of LAMC2 predicts poor prognosis in colorectal cancer patients and promotes cancer cell proliferation, migration, and invasion. Tumour Biol, 39, 1010428317705849.

60. Yousefi, M., Li, L. and Lengner, C.J. (2017) Hierarchy and Plasticity in the Intestinal Stem Cell Compartment. Trends Cell Biol, 27, 753–764.

61. Tang, X., Kuhlenschmidt, T.B., Li, Q., Ali, S., Lezmi, S., Chen, H., Pires-Alves, M., Laegreid, W.W., Saif, T.A. and Kuhlenschmidt, M.S. (2014) A mechanically-induced colon cancer cell population shows increased metastatic potential. Mol Cancer, 13, 131.

62. Vidyasekar, P., Shyamsunder, P., Arun, R., Santhakumar, R., Kapadia, N.K., Kumar, R. and Verma, R.S. (2015) Genome Wide Expression Profiling of Cancer Cell Lines Cultured in Microgravity Reveals Significant Dysregulation of Cell Cycle and MicroRNA Gene Networks. PLoS One, 10, e0135958.

63. Regnier, M., Polizzi, A., Lippi, Y., Fouche, E., Michel, G., Lukowicz, C., Smati, S., Marrot, A., Lasserre, F., Naylies, C. et al. (2018) Insights into the role of hepatocyte PPARalpha activity in response to fasting. Mol Cell Endocrinol, 471, 75–88.

64. Kennedy, L., Sandhu, J.K., Harper, M.E. and Cuperlovic-Culf, M. (2020) Role of Glutathione in Cancer: From Mechanisms to Therapies. Biomolecules, 10.

65. Kim, A.D., Zhang, R., Han, X., Kang, K.A., Piao, M.J., Maeng, Y.H., Chang, W.Y. and Hyun, J.W. (2015) Involvement of glutathione and glutathione metabolizing enzymes in human colorectal cancer cell lines and tissues. Mol Med Rep, 12, 4314–4319.

66. Bansal, A. and Simon, M.C. (2018) Glutathione metabolism in cancer progression and treatment resistance. J Cell Biol, 217, 2291–2298.

67. Narayanankutty, A., Job, J.T. and Narayanankutty, V. (2019) Glutathione, an Antioxidant Tripeptide: Dual Roles in Carcinogenesis and Chemoprevention. Curr Protein Pept Sci, 20, 907–917.

68. Brautigam, L., Zhang, J., Dreij, K., Spahiu, L., Holmgren, A., Abe, H., Tew, K.D., Townsend, D.M., Kelner, M.J., Morgenstern, R. et al. (2018) MGST1, a GSH transferase/peroxidase essential for development and hematopoietic stem cell differentiation. Redox Biol, 17, 171–179.

69. Zhang, H., Liao, L.H., Liu, S.M., Lau, K.W., Lai, A.K., Zhang, J.H., Wang, Q., Chen, X.Q., Wei, W., Liu, H. et al. (2007) Microsomal glutathione S-transferase gene polymorphisms and colorectal cancer risk in a Han Chinese population. Int J Colorectal Dis, 22, 1185–1194.

70. Peng, L., Linghu, R., Chen, D., Yang, J., Kou, X., Wang, X.Z., Hu, Y., Jiang, Y.Z. and Yang, J. (2017) Inhibition of glutathione metabolism attenuates esophageal cancer progression. Exp Mol Med, 49, e318.

71. Yang, H., Zhu, R., Zhao, X., Liu, L., Zhou, Z., Zhao, L., Liang, B., Ma, W., Zhao, J., Liu, J. et al. (2019) Sirtuin-mediated deacetylation of hnRNP A1 suppresses glycolysis and growth in hepatocellular carcinoma. Oncogene, 38, 4915–4931.

72. Grasmann, G., Smolle, E., Olschewski, H. and Leithner, K. (2019) Gluconeogenesis in cancer cells -Repurposing of a starvation-induced metabolic pathway? Biochim Biophys Acta Rev Cancer, 1872, 24–36.

73. Mears, H.V., Emmott, E., Chaudhry, Y., Hosmillo, M., Goodfellow, I.G. and Sweeney, T.R. (2019) Ifit1 regulates norovirus infection and enhances the interferon response in murine macrophage-like cells. Wellcome Open Res, 4, 82.

74. Leon-Cabrera, S., Vazquez-Sandoval, A., Molina-Guzman, E., Delgado-Ramirez, Y., Delgado-Buenrostro, N.L., Callejas, B.E., Chirino, Y.I., Perez-Plasencia, C., Rodriguez-Sosa, M., Olguin, J.E. et al. (2018) Deficiency in STAT1 Signaling Predisposes Gut Inflammation and Prompts Colorectal Cancer Development. Cancers (Basel), 10.

75. Topfer, K., Kempe, S., Muller, N., Schmitz, M., Bachmann, M., Cartellieri, M., Schackert, G. and Temme, A. (2011) Tumor evasion from T cell surveillance. J Biomed Biotechnol, 2011, 918471.

76. Kaplan, D.H., Shankaran, V., Dighe, A.S., Stockert, E., Aguet, M., Old, L.J. and Schreiber, R.D. (1998) Demonstration of an interferon gamma-dependent tumor surveillance system in immunocompetent mice. Proc Natl Acad Sci U S A, 95, 7556–7561.

77. Shankaran, V., Ikeda, H., Bruce, A.T., White, J.M., Swanson, P.E., Old, L.J. and Schreiber, R.D. (2001) IFNgamma and lymphocytes prevent primary tumour development and shape tumour immunogenicity. Nature, 410, 1107–1111.

78. Street, S.E., Trapani, J.A., MacGregor, D. and Smyth, M.J. (2002) Suppression of lymphoma and epithelial malignancies effected by interferon gamma. J Exp Med, 196, 129–134.

79. Gao, J., Shi, L.Z., Zhao, H., Chen, J., Xiong, L., He, Q., Chen, T., Roszik, J., Bernatchez, C., Woodman, S.E. et al. (2016) Loss of IFN-gamma Pathway Genes in Tumor Cells as a Mechanism of Resistance to Anti-CTLA-4 Therapy. Cell, 167, 397–404 e399.

80. Meunier, E. and Broz, P. (2016) Interferon-inducible GTPases in cell autonomous and innate immunity. Cell Microbiol, 18, 168–180.

81. Komatsu, Y., Christian, S.L., Ho, N., Pongnopparat, T., Licursi, M. and Hirasawa, K. (2015) Oncogenic Ras inhibits IRF1 to promote viral oncolysis. Oncogene, 34, 3985–3993.

82. Hong, M., Zhang, Z., Chen, Q., Lu, Y., Zhang, J., Lin, C., Zhang, F., Zhang, W., Li, X., Zhang, W. et al. (2019) IRF1 inhibits the proliferation and metastasis of colorectal cancer by suppressing the RAS-RAC1 pathway. Cancer Manag Res, 11, 369–378.

83. Ohsugi, T., Yamaguchi, K., Zhu, C., Ikenoue, T., Takane, K., Shinozaki, M., Tsurita, G., Yano, H. and Furukawa, Y. (2019) Anti-apoptotic effect by the suppression of IRF1 as a downstream of Wnt/beta-catenin signaling in colorectal cancer cells. Oncogene, 38, 6051–6064.

84. Shao, L., Hou, W., Scharping, N.E., Vendetti, F.P., Srivastava, R., Roy, C.N., Menk, A.V., Wang, Y., Chauvin, J.M., Karukonda, P. et al. (2019) IRF1 Inhibits Antitumor Immunity through the Upregulation of PD-L1 in the Tumor Cell. Cancer Immunol Res, 7, 1258–1266.

85. Kuznetsov, V.A., Makalkin, I.A., Taylor, M.A. and Perelson, A.S. (1994) Nonlinear dynamics of immunogenic tumors: parameter estimation and global bifurcation analysis. Bull Math Biol, 56, 295–321.

86. Santaolalla, R., Sussman, D.A., Ruiz, J.R., Davies, J.M., Pastorini, C., Espana, C.L., Sotolongo, J., Burlingame, O., Bejarano, P.A., Philip, S. et al. (2013) TLR4 activates the beta-catenin pathway to cause intestinal neoplasia. PLoS One, 8, e63298.

87. Fukata, M., Chen, A., Vamadevan, A.S., Cohen, J., Breglio, K., Krishnareddy, S., Hsu, D., Xu, R., Harpaz, N., Dannenberg, A.J. et al. (2007) Toll-like receptor-4 promotes the development of colitis-associated colorectal tumors. Gastroenterology, 133, 1869–1881.

88. Niedzielska, I., Niedzielski, Z., Tkacz, M., Orawczyk, T., Ziaja, K., Starzewski, J., Mazurek, U. and Markowski, J. (2009) Toll-like receptors and the tendency of normal mucous membrane to transform to polyp or colorectal cancer. J Physiol Pharmacol, 60 Suppl 1, 65–71.

89. Simiantonaki, N., Kurzik-Dumke, U., Karyofylli, G., Jayasinghe, C., Michel-Schmidt, R. and Kirkpatrick, C.J. (2007) Reduced expression of TLR4 is associated with the metastatic status of human colorectal cancer. Int J Mol Med, 20, 21–29.

90. Gwon, D.H., Lee, W.Y., Shin, N., Kim, S.I., Jeong, K., Lee, W.H., Kim, D.W., Hong, J. and Lee, S.Y. (2020) BMAL1 Suppresses Proliferation, Migration, and Invasion of U87MG Cells by Downregulating Cyclin B1, Phospho-AKT, and Metalloproteinase-9. Int J Mol Sci, 21.

91. Tang, Q., Cheng, B., Xie, M., Chen, Y., Zhao, J., Zhou, X. and Chen, L. (2017) Circadian Clock Gene Bmal1 Inhibits Tumorigenesis and Increases Paclitaxel Sensitivity in Tongue Squamous Cell Carcinoma. Cancer Res, 77, 532–544.

92. Fekry, B., Ribas-Latre, A., Baumgartner, C., Deans, J.R., Kwok, C., Patel, P., Fu, L., Berdeaux, R., Sun, K., Kolonin, M.G. et al. (2018) Incompatibility of the circadian protein BMAL1 and HNF4alpha in hepatocellular carcinoma. Nat Commun, 9, 4349.

93. Yeh, C.M., Shay, J., Zeng, T.C., Chou, J.L., Huang, T.H., Lai, H.C. and Chan, M.W. (2014) Epigenetic silencing of ARNTL, a circadian gene and potential tumor suppressor in ovarian cancer. Int J Oncol, 45, 2101–2107.

94. Asuthkar, S., Stepanova, V., Lebedeva, T., Holterman, A.L., Estes, N., Cines, D.B., Rao, J.S. and Gondi, C.S. (2013) Multifunctional roles of urokinase plasminogen activator (uPA) in cancer stemness and chemoresistance of pancreatic cancer. Mol Biol Cell, 24, 2620–2632.

95. AlHossiny, M., Luo, L., Frazier, W.R., Steiner, N., Gusev, Y., Kallakury, B., Glasgow, E., Creswell, K., Madhavan, S., Kumar, R. et al. (2016) Ly6E/K Signaling to TGFβ Promotes Breast Cancer Progression, Immune Escape, and Drug Resistance. Cancer research, 76, 3376–3386.

96. Park, J.W., Park, J.M., Park, D.M., Kim, D.Y. and Kim, H.K. (2016) Stem Cells Antigen-1 Enriches for a Cancer Stem Cell-Like Subpopulation in Mouse Gastric Cancer. Stem Cells, 34, 1177–1187.

97. Amano, H., Chaudhury, A., Rodriguez-Aguayo, C., Lu, L., Akhanov, V., Catic, A., Popov, Y.V., Verdin, E., Johnson, H., Stossi, F. et al. (2019) Telomere Dysfunction Induces Sirtuin Repression that Drives Telomere-Dependent Disease. Cell Metab.

98. Long, T.M., Chakrabarti, A., Ezelle, H.J., Brennan-Laun, S.E., Raufman, J.P., Polyakova, I., Silverman, R.H. and Hassel, B.A. (2013) RNase-L deficiency exacerbates experimental colitis and colitis-associated cancer. Inflamm Bowel Dis, 19, 1295–1305.

99. Burke, J.M., Moon, S.L., Matheny, T. and Parker, R. (2019) RNase L Reprograms Translation by Widespread mRNA Turnover Escaped by Antiviral mRNAs. Mol Cell, 75, 1203–1217 e1205.

100. Mullan, P.B., Hosey, A.M., Buckley, N.E., Quinn, J.E., Kennedy, R.D., Johnston, P.G. and Harkin, D.P. (2005) The 2,5 oligoadenylate synthetase/RNaseL pathway is a novel effector of BRCA1-and interferon-gamma-mediated apoptosis. Oncogene, 24, 5492–5501.

101. Kruger, S., Silber, A.S., Engel, C., Gorgens, H., Mangold, E., Pagenstecher, C., Holinski-Feder, E., von Knebel Doeberitz, M., Moeslein, G., Dietmaier, W. et al. (2005) Arg462Gln sequence variation in the prostate-cancer-susceptibility gene RNASEL and age of onset of hereditary non-polyposis colorectal cancer: a casecontrol study. Lancet Oncol, 6, 566–572.

102. Kruger, S., Engel, C., Bier, A., Silber, A.S., Gorgens, H., Mangold, E., Pagenstecher, C., Holinski-Feder, E., von Knebel Doeberitz, M., Royer-Pokora, B. et al. (2007) The additive effect of p53 Arg72Pro and RNASEL Arg462Gln genotypes on age of disease onset in Lynch syndrome patients with pathogenic germline mutations in MSH2 or MLH1. Cancer Lett, 252, 55–64.

103. Batman, G., Oliver, A.W., Zehbe, I., Richard, C., Hampson, L. and Hampson, I.N. (2011) Lopinavir up-regulates expression of the antiviral protein ribonuclease L in human papillomavirus-positive cervical carcinoma cells. Antivir Ther, 16, 515–525.

104. Banerjee, S., Li, G., Li, Y., Gaughan, C., Baskar, D., Parker, Y., Lindner, D.J., Weiss, S.R. and Silverman, R.H. (2015) RNase L is a negative regulator of cell migration. Oncotarget, 6, 44360–44372.

105. Kikuchi, N., Ye, J., Hirakawa, J. and Kawashima, H. (2019) Forced Expression of CXCL10 Prevents Liver Metastasis of Colon Carcinoma Cells by the Recruitment of Natural Killer Cells. Biol Pharm Bull, 42, 57–65.

106. Wangpu, X., Yang, X., Zhao, J., Lu, J., Guan, S., Lu, J., Kovacevic, Z., Liu, W., Mi, L., Jin, R. et al. (2015) The metastasis suppressor, NDRG1, inhibits “stemness” of colorectal cancer via down-regulation of nuclear beta-catenin and CD44. Oncotarget, 6, 33893–33911.

107. Sahni, S., Bae, D.H., Lane, D.J., Kovacevic, Z., Kalinowski, D.S., Jansson, P.J. and Richardson, D.R. (2014) The metastasis suppressor, N-myc downstream-regulated gene 1 (NDRG1), inhibits stress-induced autophagy in cancer cells. J Biol Chem, 289, 9692–9709.

108. Strzelczyk, B., Szulc, A., Rzepko, R., Kitowska, A., Skokowski, J., Szutowicz, A. and Pawelczyk, T. (2009) Identification of high-risk stage II colorectal tumors by combined analysis of the NDRG1 gene expression and the depth of tumor invasion. Ann Surg Oncol, 16, 1287–1294.

109. Park, K.C., Paluncic, J., Kovacevic, Z. and Richardson, D.R. (2020) Pharmacological targeting and the diverse functions of the metastasis suppressor, NDRG1, in cancer. Free Radic Biol Med, 157, 154–175.

110. Sundararaghavan, V.L., Sindhwani, P. and Hinds, T.D., Jr. (2017) Glucuronidation and UGT isozymes in bladder: new targets for the treatment of uroepithelial carcinomas? Oncotarget, 8, 3640–3648.

111. Guengerich, F.P. (2019) Cytochrome P450 research and The Journal of Biological Chemistry. J Biol Chem, 294, 1671–1680.

112. Tomita, H., Tanaka, K., Tanaka, T. and Hara, A. (2016) Aldehyde dehydrogenase 1A1 in stem cells and cancer. Oncotarget, 7, 11018–11032.

113. Krause, K., Caution, K., Badr, A., Hamilton, K., Saleh, A., Patel, K., Seveau, S., Hall-Stoodley, L., Hegazi, R., Zhang, X. et al. (2018) CASP4/caspase-11 promotes autophagosome formation in response to bacterial infection. Autophagy, 14, 1928–1942.

114. Lang, F., Perrotti, N. and Stournaras, C. (2010) Colorectal carcinoma cells--regulation of survival and growth by SGK1. Int J Biochem Cell Biol, 42, 1571–1575.

115. Wang, C., Shi, M., Ji, J., Cai, Q., Zhao, Q., Jiang, J., Liu, J., Zhang, H., Zhu, Z. and Zhang, J. (2020) Stearoyl-CoA desaturase 1 (SCD1) facilitates the growth and anti-ferroptosis of gastric cancer cells and predicts poor prognosis of gastric cancer. Aging (Albany NY), 12, 15374–15391.

