## Supplementary Methods and Discussion for "Immunity Depletion, Telomere Imbalance, and Cancer-associated Metabolism Pathway Aberrations in Intestinal Mucosa upon Caloric Restriction"

### Database usage

The collection of databases used to carefully manually curate and re-annotate differentially expressed duodenum mucosa CR-responded genes.

| Database | Reference |
|---|---|
| UniProtKB/UniProt Tissue Annotation (https://www.uniprot.org/)<br>DAVID Bioinformatics 6.7 Functional Annotation Tool (https://david-d.ncifcrf.gov/) | (1) |
| Ingenuity Pathway Analysis (IPA) Disease and Biological Function Annotations (https://www.qiagenbioinformatics.com/products/ingenuity-pathway-analysis) | (2) |
| STRING v11 Gene Ontology (GO) (https://string-db.org/) | (3) |
| Cancer Cell Metabolism Gene (CCM) Database (https://bioinfo.uth.edu/ccmGDB/) | (4) |
| Cancer Predisposition Gene (CPG) Database | (5) |
| The Tumor Suppressor Gene (TSG) Database (https://bioinfo.uth.edu/TSGene1.0/) | (6) |
| TelNet Database (www.cancertelsys.org/telnet/) | (7) |
| Cancer Stem Cell Database (CSCdb) | (8) |
| Enrichr Database (https://amp.pharm.mssm.edu/Enrichr/) | (9) |
| Mouse Genome Informatics (MGI) Batch Query tool (http://www.informatics.jax.org/batch) | (10) |
| Genome Browser (http://genome.ucsc.edu/) with mouse Dec. 2011 (GRCm38/mm10) | (11) |
| The telomerase RNA component (TERC) Path Designer via IPA | (2) |
| NCBI Reference Sequence (RefSeq) (https://www.ncbi.nlm.nih.gov/refseq/) | (12) |
| OncoMX (http://oncomx.org/) | (13) |
| dbEMT (http://dbemt.bioinfo-minzhao.org/) Epithelial-Mesenchymal Transition Gene Database | (14) |

### Gene symbols associated with multiple probesets (p.s.)

If more than one p.s. was associated with a gene symbol, the directionality of the regulation was checked. When the p.s. associated with a specific gene symbol had either all upregulation or all downregulation expression data, the p.s. with the lowest adj. p-value was chosen (Supplementary Table S11). Only one gene associated with multiple p.s. had both upregulation and downregulation data. This led to the distinction of *Armc2* having isoform A (upregulated) and isoform B (downregulated). Isoform A is associated with p.s. 10368881, is located at chr10:42008653-42008725, and is *Armc2's* second longest exon isoform. Isoform B is associated with p.s. 10368859, is located at chr10:41914993-42007709, and is *Armc2's* longest isoform. The UCSC Genome Browser (http://genome.ucsc.edu/) with mouse Dec. 2011 (GRCm38/mm10) assembly was utilized to annotate Armc2's isoforms (11).

**Gene enrichment analysis**

Enrichment analysis of tissue-associated proteins in mice was completed using the Uniprot tissue (UP_tissue) annotation database through DAVID Bioinformatics 6.7. and its Functional Annotation Tool (1). Through selection of tissues with Benjamini < 0.05, Enriched Immune System Gene (ISG) subset was represented by the proteins connected to the jejunal and colic lymph nodes (at 87.14-fold enrichment), spleen, activated spleen and thymus. Additionally, Epithelial Cell-Enriched Genes (ECG) were represented by proteins that referred to the colon, liver, kidney, and SI. Next, both the ISGs and ECGs were excluded from the 505 DEGs and returned the "Other DEGs" category. Genes related to the immune system were identified in the Other DEGs list and ECG list; the classification system was strengthened using immune resources and IPA's Disease and Biological Function annotations (15).

**Gene subset recourses**

The Mouse Genome Informatics (MGI) Batch Query tool (http://www.informatics.jax.org/batch) was used for ortholog conversions (10). Additionally, gene BLAST (https://blast.ncbi.nlm.nih.gov/Blast.cgi) determined the best homologous genes in the mouse genome for lost or re-solve conflict annotations (16).

**Expression data analysis**

Transcriptional profiling using Affymetrix Human Transcriptome Array 2.0 of duodenal biopsies from normal tissue of 12 familial adenomatous polyposis (FAP) patients who developed cancer was assessed (17). The NCBI GEO accession for this dataset is #GSE111156. This microarray was normal human duodenum expression and had 26,869 p.s. after removal of control p.s. and p.s. without annotation data from Affymetrix. For each p.s. across the 12 samples, the median expression value was utilized to represent the p.s. signal. We determined the overlap with the microarray from the CR DM scraping; the mean value was used to represent each p.s. signal. The cutoff for both datasets was 3 for log microarray signal intensity as defined from the frequency distribution expression functions; this separated expressed genes for both datasets. MGI batch query converted human gene symbols to mouse orthologs. Of the 467 mouse CR DEGs, 382 had human orthologs. When one mouse gene symbol was associated with multiple human orthologs, the protein sequences were aligned with the Clustal Omega program through UniProt (18).

**Gene ontology, network analysis, and gene set enrichment**

STRING v11 (https://string-db.org/) generated networks for mouse and orthologous human genes (3). Interaction sources were experiments and databases, with the minimum required interaction score at high confidence (0.700). The statistical background was the whole mouse or human genome. Gene Ontology (GO) information was generated by STRING for each network as defined by the COG database. Functional GO enrichments (FDR <0.05) were compared between human and mouse.

DAVID Bioinformatics 6.7 was utilized for additional GO of all mouse CR DEGs (1). The background set was the whole mouse genome. Functionally enriched DAVID terms were defined as those with a Benjamini-corrected *p*-value of less than 0.05.

Ranked Gene Set Enrichment Analysis (GSEA) was performed with software v4.1.0 (19). The gene set annotations were downloaded from: http://download.baderlab.org/EM_Genesets/current_release/Mouse/symbol/Mouse_GO_AllPathways_no_GO_iea_September_01_2020_symbol.gmt. The file excludes GO annotation evidence codes 'IEA' (inferred from electronic annotation), 'ND' (No biological data available), 'RCA' (inferred from reviewed computational analysis). Default settings were used. "Collapse dataset to gene symbols" was set to "false". The ranked list prior to analysis was calculated by the p-value and direction of the log fold-change for mouse CR DEGs with adjusted p-value < 0.05 at |FC| > 1.2.. Significant gene sets (with $p<0.05$ and FDR < 0.25) were visualized in Cytoscape v3.8.0 EnrichmentMap (20).

**Immune cell type enriched genes**

Enriched genes of immune cells types (>2-fold difference) from human and mouse expression profiles were identified from Table 1 (15). Human genes were converted to mouse orthologs and compared against the 505 CR DEGs.

**Curation of cancer-associated genes and selection criteria**

We curated genes lists from colorectal adenoma and duodenal adenoma/adenocarcinomas, the Cancer Cell Metabolism Gene Database, Apc knockout and APCMin/+ mouse model of colorectal cancer studies, and literature searches. Next, these gene lists were compared to the 505 CR DEGs for significance (adj. p-value <0.05 at |FC|>1.5).

Three supplementary files were utilized that indicated gene transcript under/over expression in human adenoma versus normal mucosa (21-23). These included duodenal adenoma/adenocarcinomas tumor-normal tissue pairs (FC ≥1.25 compared with matched normal mucosa) from Supplementary Table 2 (23), colorectal adenoma samples of 44k Whole Human Genome microarrays (Agilent) (≥ 2.0 FC, P-value ≤ 0.01 by *t*-test with FDR) from Supplementary Table 2 (21), and colorectal adenomas (FC ≥ 4.0 compared with normal mucosa and Mann-Whitney test using a 1% FDR) from Supplementary Table 4 (22). Additionally, DEGs from human duodenal cancer-normal comparisons in familial adenomatous polyposis (FAP) cases ($p<0.05$ and FC >2) were used; FAP is caused by loss of *APC* function (17). Human transcripts were converted to mouse orthologs with MGI batch query (10). Only matching directionality gene expression changes were included when comparing to the 505 CR DEGs (upregulated in both CR and adenomas or downregulated in both CR and adenomas).

The Cancer Cell Metabolism Gene Database maintains a comprehensive resource of human cancer cell metabolism (CCM) genes (https://bioinfo.uth.edu/ccmGDB/) (4). Their 514 CCM gene list (curated from five other cancer gene databases) was converted to mouse orthologs using MGI batch query and compared to our 505 CR DEGs (10). The directionality of the CC gene expression changes were not indicated.

Top regulated genes following Apc loss at days four and five (FDR of 5%) from Supplementary Table 1a-c (24) were compared to our 505 CR DEGs. Additionally, transcription profiles of $APC^{Min/+}$ adenomas and carcinomas compared to wild-type epithelial and $APC^{Min/+}$ normal epithelial cells (Mann-Whitney pairwise comparison test, $p \leq 0.05$) were overlapped with our 505 CR DEGs (25). Only matching directionality gene expression changes were included (Apc knockout/$APC^{Min/+}$ upregulated and CR downregulated or Apc knockout/$APC^{Min/+}$ downregulated and CR upregulated).

**Curation of telomeric maintenance genes and selection criteria**

After preliminary network analysis of the upregulated ECGs using STRING v11, we hypothesized that CR may dysregulate telomere maintenance (3). We curated genes from databases *TelNet* and

Ingenuity Pathway Analysis (IPA), literature searches, and manual curation. Human genes were converted to mouse orthologs using MGI Batch Query (10). Next, these gene lists were compared to the 26,966 p.s. and were compiled using statistical criteria of adjusted p-value <0.05 at |FC|>1.2.

The *TelNet* database included a list of 132 significant human genes in Table S1 involved in telomere maintenance after pan-cancer analysis of tumor compared to normal cells (p-value < 0.01, log2 ratio below − 0.782 or above 0.852) or an (anti-) correlation of gene expression and telomere length ratios (p-value < 0.01, Rho below − 0. 295 or above 0.186) (7). The TERC Path Designer via IPA (QIAGEN Inc., https://www.qiagenbioinformatics.com/products/ingenuity-pathway-analysis) generated a list of 45 human genes involved in the telomerase RNA component (TERC) pathway (2). RNA-seq analysis of TERRA knockdown in mouse embryonic stem cell gene expression (log 2 (FC) > +/- 1 and p<0.05) identified 199 genes in Table S1 (26). In addition, the protein interactome of TERRA was characterized with 134 protein partners, which are involved in chromatin modification/transcription in Table S2 (26). PubMed was used to search for articles relating to telomeres, their maintenance, and CR gene expression. Lastly, genes were curated via manual search in the 26,966 p.s. for Telo*.

**Interferon-inducible GTPase family annotations**

Multiple alignment and blat/blast mapping of the Affymetrix micro-array individual probe sets (p.s.) (designed in UTR and exons) showed many p.s. have multiple alignment across the IFN-inducible GTPase genes. It is almost impossible to unambiguously assign to one gene or the other certain gene pairs (*9930111J21Rik1, 9930111J21Rik2* and *Tgt1, Tgtp2*) based on sum and individual p.s. expression signals that is used as endpoint mRNA abundance signal. However, for other genes, microarray p.s. design allows us to develop bioinformatics strategies that reduces or fully excludes such uncertainty.

Chromosome coordinates of the Chr11 qB(1.2) locus IFN-inducible GTPase family are provided in Supplementary Data 5 Table B. Exon sequence data was exported from the UCSC genome browser (https://genome.ucsc.edu/) track NCBI RefSeq for the genes of interest (selecting for 5'UTR exons, CDS exons, 3' UTR exons): *Irgm1, Gm5431, Gm12185, Gm12186, 9930111J21Rik1, 9930111J21Rik2, Tgtp1, Tgtp2.* Next, we built a custom BLAST database using the makeblastdb application. Within the NCBI Genome Workbench, all probe set sequences (ProbeIDs) associated with a Transcript Cluster ID were ran against the local exon sequence database with BLASTn (using default parameters wordsize=11, E-value=-10.00). For each Transcript Cluster ID, only ProbeIDs with greater than 90% in query coverage and identity match were included in the 8x8 matrix in Supplementary Data 5 Table F. Supplementary Figure 4 contains UCSC genome browser tracks defining paralogs of the interferon-inducible GTPase family located at other loci than Chr11 including F830016B08Rik, Iigp1, Irgm2, Igtp, and Irgc1.

**Network Analysis**

A core expression analysis was run in IPA (QIAGEN Inc., https://www.qiagenbioinformatics.com/products/ingenuity-pathway-analysis) based on the expression fold changes of our p.s. for the 467 mouse CR DEGs, (2). The reference set was the Mouse Gene 1.0 ST Array and the confidence was experimentally observed. Fischer's exact test in IPA determined enriched canonical pathways for each subset. The Disease and Biological Function annotations within IPA strengthened the curation of gene category subsets for the cancer-associated and ISGs and identified additional immune genes in the Other category. Lastly, core expression analyses were run in IPA separately on each mouse subset (tumor/cancer, ISG, ECG, telomere) using the same criteria as the 467 CR DEGs. After manual curation and inspection, the

IPA Networks were imported into Cytoscape v3.8.0 (27). Unconnected nodes were removed from the analysis. The network analyzer tool calculated Cytoscape network statistics using directed analysis. This identified the number of proteins, edges (interactions), average number of neighbors, clustering coefficient, and network density. Lastly, network data were imported into STRING v11 to determine protein-protein interaction (PPI) enrichment p-values.

**Supplementary Discussion**

Selected CR DEGs in tumor suppressive, oncogenic, immune, epithelial stem-like, anti-cancer, and detoxifying pathways for future validation and consideration in pre-cancer and metabolic reprogramming states.

| Gene Name | CR | FC | Annotation | References |
|---|---|---|---|---|
| Fkbp5 | Up | 8.54 | most over-expressed CR-response gene, androgen-responsive gene with high expression in esophageal adenocarcinoma (EAC) tissues and this is associated with decreased patient survival, pro-oncogenic role in EAC | (28-30) |
| Tlr4 | Down | -1.77 | Innate immune system, pathogen recognition, therapeutic target, reduced expression associated with metastatic status of CRC | (31-34) |
| Arntl | Down | -1.87 | tumor suppressor, circadian rhythms | (35-38) |
| Plau | Up | 1.78 | cancer stem-like, poor pancreatic ductal adenocarcinoma prognosis | (39) |
| Ly6a, Ly6e, Ly6c1 | Down | -2.67, -2.18, -1.63 | immune cell differentiation, cancer stem cell biology | (40,41) |
| Sirt3 | Down | -1.38 | oxidative stress, protection of DNA damage, chromosome maintenance | (42-45) |
| Rnasel | Down | -1.74 | tumor suppressor, predisposition to prostate cancer, antiviral pathways | (46-52) |
| Cxcl9, Cxcl10 | Down | -1.79, -2.03 | proinflammatory chemokines, therapeutic target | (53,54) |
| Ndrg1 | Up | 2.65 | tumor suppressor, EMT, therapeutic target | (55-58) |
| Ugt2b5, Ugt2b35, Ugt2b36 | Up | 2.5, 1.7, 2.56 | detoxifying enzymes, reduce risk of carcinogenesis and toxicities by inactivating aromatic-like metabolites | (59) |
| Cyp2c55 (and family) | Up | 2.70 | metabolizing endogenous compounds, detoxifying exogenous chemicals, drug metabolism | (60) |
| Lamc2 | Up | 1.75 | promotes proliferation, cell migration, and invasion in cancers including colorectal and malignant metastases | (61-63) |
| Rrm2 | Up | 4.02 | cell cycle, therapeutic target, oncogene playing a key role in tumorigenesis and cancer progression, poor prognostic factor for colon, breast, and pancreatic cancers, cancer driver | (64-68) |
| Aldh1a1 | Up | 2.45 | EMT-related oncogenic, cancer-stem like, upregulated in $APC^{Min/+}$ mouse model of colorectal cancer | (69) |
| Casp4 | Down | -1.87 | apoptosis, CASP4-deficicient mice exhibit a defect in autophagy | (70) |
| Cemip | Up | 4.25 | overexpression correlates with poorer colon cancer patient survival and facilitates colorectal and stomach tumor growth, cancer driver | (61,71) |

*Tlr4 was downregulated upon CR. Tlr4* interacted with the hubs *Stat1, Stat2, Cxcl10, Irf1, Nos2,* and *Pml. Tlr1/3* were also downregulated. Toll-like receptor 4 (TLR4) promotes neoplasia through activation of the β-catenin pathway and 40% of sporadic CRC and 20% of colon adenomas over-express TLR4 (31). TLR4 signalling in the colon induces epithelial proliferation and blocking TLR4 may reduce tumor development (32). TLRs are involved in the progression from precancerous polyps to tumors (33). However, studies have also demonstrated strong reduced expression of TLR4 is associated with increased metastatic potential of CRC (34).

CR-regulated DM DEGs overlapped with TERRA knockdown and protein interactome data indicating chromosomal instability and telomere maintenance imbalance was likely. In single telomere depletion of telomeric repeat-containing RNA (TERRA) transcripts in cancer cells, DNA damage was induced at telomeres and extratelomeric sites in the genome (72). TERRA molecules bind chromatin throughout the genome (26). Thus, in CR response DM DNA damage due to TERRA depletion may alter the genomic integrity of chromatin sites normally bound to TERRA (72).

*Arntl was suppressed upon CR.* The *BMAL1/ARNTL* gene acts as a TSG. Brain and muscle aryl hydrocarbon receptor nuclear translocator-like 1 (*BMAL1/ARNTL)* maintains circadian rhythms and inhibits growth and metastasis of tumor cells in lung, ovarian, and breast cancer and tongue squamous cell carcinoma (35-38). *BMAL1* alters the proliferation, migration, and invasion of cancer cells (35). Therefore, *BMAL1* could be a potential therapeutic target for cancer treatment. Significant *Arntl* suppression found in our work suggests that treatment of *Arntl* expression may be a perspective strategy for reducing risk of cancerous effects in CR conditions.

*Irf1 was suppressed upon CR.* IRF1 expression was lower in CRC than in normal mucosa and inversely associated with CRC proliferation rate and metastasis (73). Ras/MEK activation in cancer cells downregulates IFN-inducible gene transcription by targeting IRF1 expression, increasing susceptibility to viral oncolysis (74). Wnt signaling pathway impairment is involved in CRC development via activating anti-apoptotic properties of tumors associated with IRF1 degradation (75). However, pro-oncogenic function(s) of IRF1 is possible; IRF1-deficient tumor cells lost the ability to upregulate PD-L1 expression in vivo in cancer that enhances tumor-infiltrating T-cell cytotoxicity (76).

All CSC genes are multifunctional and context-dependent; this allows alternative pathways. For instance, *Aldh1a1, Ll2rg*, and *Dll4* are also involved in telomerase maintaining and signalling. CSCs have been implicated in chemoresistance properties and ALDH1A1 expression is involved in resistance to gemcitabine chemotherapy for pancreatic cancer treatment (77). Urokinase plasminogen activator (PLAU) was upregulated upon CR in our study; poor pancreatic ductal adenocarcinoma prognosis has been correlated with increased expression of PLAU and following its suppression, tumorigenicity, and gemcitabine resistance decreases (39). The Lymphocyte antigen-6 (Ly6) gene family members *Ly6a, Ly6e, Ly6c1*—which are involved in immune cell differentiation—were suppressed upon CR. Expression of human LY6E is required for tumor immune escape and its knockdown reduced PDl1 (immune checkpoint molecule) expression after IFN-γ treatment (40). Upregulation of Stem Cells Antigen-1 (Sca-1) encoded by the Ly6a gene enriches tumorigenicity, metastasis, and chemoresistance in mouse gastric cancer (41).

*Sirt3 was suppressed upon CR.* Current data highlights recent advances and controversies regarding the yin and yang functions of SIRT3 in cancer. Sirt3 over-expression is associated with the reduction of oxidative stress, protection of DNA damage, chromosome maintenance, acetylation of histones, tumor suppression, and metabolic control of cellular homeostasis (42,43). SIRT1, SIRT3, and SIRT6 mediate hnRNP A1 deacetylation and inhibit both proliferation and tumorigenesis (44). However, murine tumors lacking Sirt3 induce genome instability and SIRT3 can act as a tumor suppressor by reducing reactive oxygen species (ROS) and regulating HIF-1α (42,43). SIRT3 protein levels are decreased in human breast cancers, and its knockout in mice has led to age-related diseases (78). Sirt3 expression downregulation is functionally connected to telomere dysfunction. Telomere shortening in livers of telomerase knockout mice repressed all seven sirtuins (45).

*A key tumor suppressor, RNase L, is strongly suppressed in response to CR.* The RNase L gene (RNASEL) encodes a component of the interferon-regulated 2-5A system that functions through antiviral, antibacterial, and anti-proliferative activities (46,47). Interferon-activated 2', 5'-oligoadenylate synthetase (OAS) induces RNaseL to cleave cellular and viral mRNA resulting in apoptosis, autophagy and inflammation (48). *Oas1a, Oas1b, Oas1g, Oas2*, and *Oasl2* were downregulated upon CR, explaining a mechanism for increased pathogen susceptibility. RNase L induces STAT1 and BRAC1 dependent interferon-gamma (IFN-γ) responses (48) and promotes innate immune responses to intestinal damage, ameliorating murine colitis and colitis-associated cancer (46). RNase L is a candidate for the hereditary prostate cancer 1 (HPC1) allele and its gene mutations are associated with predisposition to prostate cancer (48-52). Functionally different variants of SNP in RNASEL are associated with the age of disease onset of hereditary non-polyposis CRC in Lynch syndrome patients (49,50). RNase L suppression (or mutation) upon CR may play negative roles in the mucosa and therapeutic expression control may be essential.

*Proinflammatory chemokines Cxcl9/10 were CR-downregulated.* The type II interferon signalling molecule *Cxcl10* had enriched interactions with immunity network hubs (*Stat1/2, Tlr4, Irf*). The analysis noted an asymmetry in input/output connections for *Cxcl10* with 1 downstream and 16 upstream targets. Changes in CXCL10 expression are associated with infectious disease, immune dysfunction, and tumor development (53). *Cxcl10* mediates leukocyte trafficking by activating T lymphocytes (Th1), NK cells, macrophages, dendritic and B cells (53). CXCL9/10 aortic concentrations increase due to aging; however, CR prevented age-related increases in CXCL10 (79-82). CXCL10 stable expression in vivo growing and metastasized colon carcinoma cells is controlled by recruiting and cytolytic functions of NK cells (54). CR-mediated CXCL10 reduction in mucosa epithelial cells and NK cell numbers may increase cancer development risks. CXCL10 therapeutic implications have been identified (53).

*Predicted CR modulated therapeutic target, Ndrg1.* Our results showed CR induces the expression of N-myc downstream regulated gene-1 (NDRG1), a marker of normal epithelial cells. Ndrg1 pharmacological targeting is a promising cancer therapeutic strategy because it plays potent metastasis suppressing roles by inducing apoptosis. More tumorigenic and stem-cell like properties were found in NDRG1-silenced CRC cells (55). NDRG1 suppresses the stress-induced pro-survival autophagic pathway (56). In colon cancer, NDRG1 expression declines as normal colonic epithelium progresses to carcinoma (57); patients with lowered NDRG1 mRNA had a shorter 5-year survival rate. However, pro-oncogenic pleiotropic roles have been reported in other cancer-types (58). NDRG1 anti-tumor activities may be crucial in CR conditions.

*Regulation of detoxification enzymes upon CR.* UDP-glucuronosyltransferases (UGTs) reduce the risk of mutagenesis, carcinogenesis, and toxicities by inactivating aromatic-like metabolites (83). This detoxifying system (*Ugt2b5, Ugt2b35, Ugt2b36)* is upregulated upon CR. UGT enzymes are suppressed in several cancers, including bladder cancer by allowing carcinogen accumulation (59). The negative modifications in tumor suppressors and pro-oncogenic upregulation in CR mice may be combatted by the increases in detoxifying enzymes. The cytochrome P450 (CYP) superfamily is involved in metabolizing endogenous compounds, detoxifying exogenous chemicals, and drug metabolism; furthermore, deficiencies in several P450s have been demonstrated in human diseases (60). CYP genes upregulated in CR mice were *Cyp2b10, Cyp2c55, Cyp2d10, Cyp2d9, Cyp2j6, Cyp3a44,* and *Cyp4b1*. According to CR-induced expression profiles and network analysis, the glutathione pathway and chemical carcinogenesis rewiring processes are interconnected, but in some contexts, their balance might be lost.

### References


1. Huang da, W., Sherman, B.T. and Lempicki, R.A. (2009) Systematic and integrative analysis of large gene lists using DAVID bioinformatics resources. *Nat Protoc*, **4**, 44-57.
2. Kramer, A., Green, J., Pollard, J., Jr. and Tugendreich, S. (2014) Causal analysis approaches in Ingenuity Pathway Analysis. *Bioinformatics*, **30**, 523-530.
3. Szklarczyk, D., Gable, A.L., Lyon, D., Junge, A., Wyder, S., Huerta-Cepas, J., Simonovic, M., Doncheva, N.T., Morris, J.H., Bork, P. *et al.* (2019) STRING v11: protein-protein

association networks with increased coverage, supporting functional discovery in genome-wide experimental datasets. *Nucleic Acids Res*, **47**, D607-D613.
4. Kim, P., Cheng, F., Zhao, J. and Zhao, Z. (2016) ccmGDB: a database for cancer cell metabolism genes. *Nucleic Acids Res*, **44**, D959-968.
5. Wei, R., Yao, Y., Yang, W., Zheng, C.H., Zhao, M. and Xia, J. (2016) dbCPG: A web resource for cancer predisposition genes. *Oncotarget*, **7**, 37803-37811.
6. Zhao, M., Kim, P., Mitra, R., Zhao, J. and Zhao, Z. (2016) TSGene 2.0: an updated literature-based knowledgebase for tumor suppressor genes. *Nucleic Acids Res*, **44**, D1023-1031.
7. Braun, D.M., Chung, I., Kepper, N., Deeg, K.I. and Rippe, K. (2018) TelNet - a database for human and yeast genes involved in telomere maintenance. *BMC Genet*, **19**, 32.
8. Shen, Y., Yao, H., Li, A. and Wang, M. (2016) CSCdb: a cancer stem cells portal for markers, related genes and functional information. *Database (Oxford)*, **2016**.
9. Kuleshov, M.V., Jones, M.R., Rouillard, A.D., Fernandez, N.F., Duan, Q., Wang, Z., Koplev, S., Jenkins, S.L., Jagodnik, K.M., Lachmann, A. *et al.* (2016) Enrichr: a comprehensive gene set enrichment analysis web server 2016 update. *Nucleic Acids Res*, **44**, W90-97.
10. Bult, C.J., Blake, J.A., Smith, C.L., Kadin, J.A., Richardson, J.E. and Mouse Genome Database, G. (2019) Mouse Genome Database (MGD) 2019. *Nucleic Acids Res*, **47**, D801-D806.
11. Kent, W.J., Sugnet, C.W., Furey, T.S., Roskin, K.M., Pringle, T.H., Zahler, A.M. and Haussler, D. (2002) The human genome browser at UCSC. *Genome Res*, **12**, 996-1006.
12. O'Leary, N.A., Wright, M.W., Brister, J.R., Ciufo, S., Haddad, D., McVeigh, R., Rajput, B., Robbertse, B., Smith-White, B., Ako-Adjei, D. *et al.* (2016) Reference sequence (RefSeq) database at NCBI: current status, taxonomic expansion, and functional annotation. *Nucleic Acids Res*, **44**, D733-745.
13. Dingerdissen, H.M., Bastian, F., Vijay-Shanker, K., Robinson-Rechavi, M., Bell, A., Gogate, N., Gupta, S., Holmes, E., Kahsay, R., Keeney, J. *et al.* (2020) OncoMX: A Knowledgebase for Exploring Cancer Biomarkers in the Context of Related Cancer and Healthy Data. *JCO Clin Cancer Inform*, **4**, 210-220.
14. Zhao, M., Liu, Y., Zheng, C. and Qu, H. (2019) dbEMT 2.0: An updated database for epithelial-mesenchymal transition genes with experimentally verified information and precalculated regulation information for cancer metastasis. *J Genet Genomics*, **46**, 595-597.
15. Lyons, Y.A., Wu, S.Y., Overwijk, W.W., Baggerly, K.A. and Sood, A.K. (2017) Immune cell profiling in cancer: molecular approaches to cell-specific identification. *NPJ Precis Oncol*, **1**, 26.
16. Boratyn, G.M., Camacho, C., Cooper, P.S., Coulouris, G., Fong, A., Ma, N., Madden, T.L., Matten, W.T., McGinnis, S.D., Merezhuk, Y. *et al.* (2013) BLAST: a more efficient report with usability improvements. *Nucleic Acids Res*, **41**, W29-33.
17. Thiruvengadam, S.S., O'Malley, M., LaGuardia, L., Lopez, R., Wang, Z., Shadrach, B.L., Chen, Y., Li, C., Veigl, M.L., Barnholtz-Sloan, J.S. *et al.* (2019) Gene Expression Changes Accompanying the Duodenal Adenoma-Carcinoma Sequence in Familial Adenomatous Polyposis. *Clin Transl Gastroenterol*, **10**, e00053.
18. Sievers, F. and Higgins, D.G. (2018) Clustal Omega for making accurate alignments of many protein sequences. *Protein Sci*, **27**, 135-145.
19. Subramanian, A., Tamayo, P., Mootha, V.K., Mukherjee, S., Ebert, B.L., Gillette, M.A., Paulovich, A., Pomeroy, S.L., Golub, T.R., Lander, E.S. *et al.* (2005) Gene set enrichment

analysis: a knowledge-based approach for interpreting genome-wide expression profiles. *Proc Natl Acad Sci U S A*, **102**, 15545-15550.
20. Merico, D., Isserlin, R., Stueker, O., Emili, A. and Bader, G.D. (2010) Enrichment map: a network-based method for gene-set enrichment visualization and interpretation. *PLoS One*, **5**, e13984.
21. Pesson, M., Volant, A., Uguen, A., Trillet, K., De La Grange, P., Aubry, M., Daoulas, M., Robaszkiewicz, M., Le Gac, G., Morel, A. *et al.* (2014) A gene expression and pre-mRNA splicing signature that marks the adenoma-adenocarcinoma progression in colorectal cancer. *PLoS One*, **9**, e87761.
22. Sabates-Bellver, J., Van der Flier, L.G., de Palo, M., Cattaneo, E., Maake, C., Rehrauer, H., Laczko, E., Kurowski, M.A., Bujnicki, J.M., Menigatti, M. *et al.* (2007) Transcriptome profile of human colorectal adenomas. *Mol Cancer Res*, **5**, 1263-1275.
23. Sakaguchi, Y., Yamamichi, N., Tomida, S., Takeuchi, C., Kageyama-Yahara, N., Takahashi, Y., Shiogama, K., Inada, K.I., Ichinose, M., Fujishiro, M. *et al.* (2018) Identification of marker genes and pathways specific to precancerous duodenal adenomas and early stage adenocarcinomas. *J Gastroenterol*.
24. Sansom, O.J., Reed, K.R., Hayes, A.J., Ireland, H., Brinkmann, H., Newton, I.P., Batlle, E., Simon-Assmann, P., Clevers, H., Nathke, I.S. *et al.* (2004) Loss of Apc in vivo immediately perturbs Wnt signaling, differentiation, and migration. *Genes Dev*, **18**, 1385-1390.
25. Paoni, N.F., Feldman, M.W., Gutierrez, L.S., Ploplis, V.A. and Castellino, F.J. (2003) Transcriptional profiling of the transition from normal intestinal epithelia to adenomas and carcinomas in the APCMin/+ mouse. *Physiol Genomics*, **15**, 228-235.
26. Chu, H.P., Cifuentes-Rojas, C., Kesner, B., Aeby, E., Lee, H.G., Wei, C., Oh, H.J., Boukhali, M., Haas, W. and Lee, J.T. (2017) TERRA RNA Antagonizes ATRX and Protects Telomeres. *Cell*, **170**, 86-101 e116.
27. Shannon, P., Markiel, A., Ozier, O., Baliga, N.S., Wang, J.T., Ramage, D., Amin, N., Schwikowski, B. and Ideker, T. (2003) Cytoscape: a software environment for integrated models of biomolecular interaction networks. *Genome Res*, **13**, 2498-2504.
28. Ellsworth, K.A., Eckloff, B.W., Li, L., Moon, I., Fridley, B.L., Jenkins, G.D., Carlson, E., Brisbin, A., Abo, R., Bamlet, W. *et al.* (2013) Contribution of FKBP5 genetic variation to gemcitabine treatment and survival in pancreatic adenocarcinoma. *PLoS One*, **8**, e70216.
29. Palethorpe, H.M., Drew, P.A. and Smith, E. (2017) Androgen Signaling in Esophageal Adenocarcinoma Cell Lines In Vitro. *Dig Dis Sci*, **62**, 3402-3414.
30. Smith, E., Palethorpe, H.M., Ruszkiewicz, A.R., Edwards, S., Leach, D.A., Underwood, T.J., Need, E.F. and Drew, P.A. (2016) Androgen Receptor and Androgen-Responsive Gene FKBP5 Are Independent Prognostic Indicators for Esophageal Adenocarcinoma. *Dig Dis Sci*, **61**, 433-443.
31. Santaolalla, R., Sussman, D.A., Ruiz, J.R., Davies, J.M., Pastorini, C., Espana, C.L., Sotolongo, J., Burlingame, O., Bejarano, P.A., Philip, S. *et al.* (2013) TLR4 activates the beta-catenin pathway to cause intestinal neoplasia. *PLoS One*, **8**, e63298.
32. Fukata, M., Chen, A., Vamadevan, A.S., Cohen, J., Breglio, K., Krishnareddy, S., Hsu, D., Xu, R., Harpaz, N., Dannenberg, A.J. *et al.* (2007) Toll-like receptor-4 promotes the development of colitis-associated colorectal tumors. *Gastroenterology*, **133**, 1869-1881.
33. Niedzielska, I., Niedzielski, Z., Tkacz, M., Orawczyk, T., Ziaja, K., Starzewski, J., Mazurek, U. and Markowski, J. (2009) Toll-like receptors and the tendency of normal mucous membrane to transform to polyp or colorectal cancer. *J Physiol Pharmacol*, **60 Suppl 1**, 65-71.

34. Simiantonaki, N., Kurzik-Dumke, U., Karyofylli, G., Jayasinghe, C., Michel-Schmidt, R. and Kirkpatrick, C.J. (2007) Reduced expression of TLR4 is associated with the metastatic status of human colorectal cancer. *Int J Mol Med*, **20**, 21-29.
35. Gwon, D.H., Lee, W.Y., Shin, N., Kim, S.I., Jeong, K., Lee, W.H., Kim, D.W., Hong, J. and Lee, S.Y. (2020) BMAL1 Suppresses Proliferation, Migration, and Invasion of U87MG Cells by Downregulating Cyclin B1, Phospho-AKT, and Metalloproteinase-9. *Int J Mol Sci*, **21**.
36. Tang, Q., Cheng, B., Xie, M., Chen, Y., Zhao, J., Zhou, X. and Chen, L. (2017) Circadian Clock Gene Bmal1 Inhibits Tumorigenesis and Increases Paclitaxel Sensitivity in Tongue Squamous Cell Carcinoma. *Cancer Res*, **77**, 532-544.
37. Fekry, B., Ribas-Latre, A., Baumgartner, C., Deans, J.R., Kwok, C., Patel, P., Fu, L., Berdeaux, R., Sun, K., Kolonin, M.G. *et al.* (2018) Incompatibility of the circadian protein BMAL1 and HNF4alpha in hepatocellular carcinoma. *Nat Commun*, **9**, 4349.
38. Yeh, C.M., Shay, J., Zeng, T.C., Chou, J.L., Huang, T.H., Lai, H.C. and Chan, M.W. (2014) Epigenetic silencing of ARNTL, a circadian gene and potential tumor suppressor in ovarian cancer. *Int J Oncol*, **45**, 2101-2107.
39. Asuthkar, S., Stepanova, V., Lebedeva, T., Holterman, A.L., Estes, N., Cines, D.B., Rao, J.S. and Gondi, C.S. (2013) Multifunctional roles of urokinase plasminogen activator (uPA) in cancer stemness and chemoresistance of pancreatic cancer. *Mol Biol Cell*, **24**, 2620-2632.
40. AlHossiny, M., Luo, L., Frazier, W.R., Steiner, N., Gusev, Y., Kallakury, B., Glasgow, E., Creswell, K., Madhavan, S., Kumar, R. *et al.* (2016) Ly6E/K Signaling to TGFβ Promotes Breast Cancer Progression, Immune Escape, and Drug Resistance. *Cancer research*, **76**, 3376-3386.
41. Park, J.W., Park, J.M., Park, D.M., Kim, D.Y. and Kim, H.K. (2016) Stem Cells Antigen-1 Enriches for a Cancer Stem Cell-Like Subpopulation in Mouse Gastric Cancer. *Stem Cells*, **34**, 1177-1187.
42. Chen, Y., Fu, L.L., Wen, X., Wang, X.Y., Liu, J., Cheng, Y. and Huang, J. (2014) Sirtuin-3 (SIRT3), a therapeutic target with oncogenic and tumor-suppressive function in cancer. *Cell Death Dis*, **5**, e1047.
43. Haigis, M.C., Deng, C.X., Finley, L.W., Kim, H.S. and Gius, D. (2012) SIRT3 is a mitochondrial tumor suppressor: a scientific tale that connects aberrant cellular ROS, the Warburg effect, and carcinogenesis. *Cancer Res*, **72**, 2468-2472.
44. Yang, H., Zhu, R., Zhao, X., Liu, L., Zhou, Z., Zhao, L., Liang, B., Ma, W., Zhao, J., Liu, J. *et al.* (2019) Sirtuin-mediated deacetylation of hnRNP A1 suppresses glycolysis and growth in hepatocellular carcinoma. *Oncogene*, **38**, 4915-4931.
45. Amano, H., Chaudhury, A., Rodriguez-Aguayo, C., Lu, L., Akhanov, V., Catic, A., Popov, Y.V., Verdin, E., Johnson, H., Stossi, F. *et al.* (2019) Telomere Dysfunction Induces Sirtuin Repression that Drives Telomere-Dependent Disease. *Cell Metab*.
46. Long, T.M., Chakrabarti, A., Ezelle, H.J., Brennan-Laun, S.E., Raufman, J.P., Polyakova, I., Silverman, R.H. and Hassel, B.A. (2013) RNase-L deficiency exacerbates experimental colitis and colitis-associated cancer. *Inflamm Bowel Dis*, **19**, 1295-1305.
47. Burke, J.M., Moon, S.L., Matheny, T. and Parker, R. (2019) RNase L Reprograms Translation by Widespread mRNA Turnover Escaped by Antiviral mRNAs. *Mol Cell*, **75**, 1203-1217 e1205.
48. Mullan, P.B., Hosey, A.M., Buckley, N.E., Quinn, J.E., Kennedy, R.D., Johnston, P.G. and Harkin, D.P. (2005) The 2,5 oligoadenylate synthetase/RNaseL pathway is a novel effector of BRCA1- and interferon-gamma-mediated apoptosis. *Oncogene*, **24**, 5492-5501.

49. Kruger, S., Silber, A.S., Engel, C., Gorgens, H., Mangold, E., Pagenstecher, C., Holinski-Feder, E., von Knebel Doeberitz, M., Moeslein, G., Dietmaier, W. *et al.* (2005) Arg462Gln sequence variation in the prostate-cancer-susceptibility gene RNASEL and age of onset of hereditary non-polyposis colorectal cancer: a case-control study. *Lancet Oncol*, **6**, 566-572.
50. Kruger, S., Engel, C., Bier, A., Silber, A.S., Gorgens, H., Mangold, E., Pagenstecher, C., Holinski-Feder, E., von Knebel Doeberitz, M., Royer-Pokora, B. *et al.* (2007) The additive effect of p53 Arg72Pro and RNASEL Arg462Gln genotypes on age of disease onset in Lynch syndrome patients with pathogenic germline mutations in MSH2 or MLH1. *Cancer Lett*, **252**, 55-64.
51. Batman, G., Oliver, A.W., Zehbe, I., Richard, C., Hampson, L. and Hampson, I.N. (2011) Lopinavir up-regulates expression of the antiviral protein ribonuclease L in human papillomavirus-positive cervical carcinoma cells. *Antivir Ther*, **16**, 515-525.
52. Banerjee, S., Li, G., Li, Y., Gaughan, C., Baskar, D., Parker, Y., Lindner, D.J., Weiss, S.R. and Silverman, R.H. (2015) RNase L is a negative regulator of cell migration. *Oncotarget*, **6**, 44360-44372.
53. Liu, M., Guo, S., Hibbert, J.M., Jain, V., Singh, N., Wilson, N.O. and Stiles, J.K. (2011) CXCL10/IP-10 in infectious diseases pathogenesis and potential therapeutic implications. *Cytokine Growth Factor Rev*, **22**, 121-130.
54. Kikuchi, N., Ye, J., Hirakawa, J. and Kawashima, H. (2019) Forced Expression of CXCL10 Prevents Liver Metastasis of Colon Carcinoma Cells by the Recruitment of Natural Killer Cells. *Biol Pharm Bull*, **42**, 57-65.
55. Wangpu, X., Yang, X., Zhao, J., Lu, J., Guan, S., Lu, J., Kovacevic, Z., Liu, W., Mi, L., Jin, R. *et al.* (2015) The metastasis suppressor, NDRG1, inhibits "stemness" of colorectal cancer via down-regulation of nuclear beta-catenin and CD44. *Oncotarget*, **6**, 33893-33911.
56. Sahni, S., Bae, D.H., Lane, D.J., Kovacevic, Z., Kalinowski, D.S., Jansson, P.J. and Richardson, D.R. (2014) The metastasis suppressor, N-myc downstream-regulated gene 1 (NDRG1), inhibits stress-induced autophagy in cancer cells. *J Biol Chem*, **289**, 9692-9709.
57. Strzelczyk, B., Szulc, A., Rzepko, R., Kitowska, A., Skokowski, J., Szutowicz, A. and Pawelczyk, T. (2009) Identification of high-risk stage II colorectal tumors by combined analysis of the NDRG1 gene expression and the depth of tumor invasion. *Ann Surg Oncol*, **16**, 1287-1294.
58. Park, K.C., Paluncic, J., Kovacevic, Z. and Richardson, D.R. (2020) Pharmacological targeting and the diverse functions of the metastasis suppressor, NDRG1, in cancer. *Free Radic Biol Med*, **157**, 154-175.
59. Sundararaghavan, V.L., Sindhwani, P. and Hinds, T.D., Jr. (2017) Glucuronidation and UGT isozymes in bladder: new targets for the treatment of uroepithelial carcinomas? *Oncotarget*, **8**, 3640-3648.
60. Guengerich, F.P. (2019) Cytochrome P450 research and The Journal of Biological Chemistry. *J Biol Chem*, **294**, 1671-1680.
61. Uhlen, M., Zhang, C., Lee, S., Sjostedt, E., Fagerberg, L., Bidkhori, G., Benfeitas, R., Arif, M., Liu, Z., Edfors, F. *et al.* (2017) A pathology atlas of the human cancer transcriptome. *Science*, **357**.
62. Demark-Wahnefried, W., Rais-Bahrami, S., Desmond, R.A., Gordetsky, J.B., Hunter, G.R., Yang, E.S., Azrad, M., Fruge, A.D., Tsuruta, Y., Norian, L.A. *et al.* (2017) Presurgical weight loss affects tumour traits and circulating biomarkers in men with prostate cancer. *Br J Cancer*, **117**, 1303-1313.

63. Huang, D., Du, C., Ji, D., Xi, J. and Gu, J. (2017) Overexpression of LAMC2 predicts poor prognosis in colorectal cancer patients and promotes cancer cell proliferation, migration, and invasion. *Tumour Biol*, **39**, 1010428317705849.
64. Furuta, E., Okuda, H., Kobayashi, A. and Watabe, K. (2010) Metabolic genes in cancer: their roles in tumor progression and clinical implications. *Biochim Biophys Acta*, **1805**, 141-152.
65. Liu, X., Zhang, H., Lai, L., Wang, X., Loera, S., Xue, L., He, H., Zhang, K., Hu, S., Huang, Y. *et al.* (2013) Ribonucleotide reductase small subunit M2 serves as a prognostic biomarker and predicts poor survival of colorectal cancers. *Clin Sci (Lond)*, **124**, 567-578.
66. Fujita, H., Ohuchida, K., Mizumoto, K., Itaba, S., Ito, T., Nakata, K., Yu, J., Kayashima, T., Souzaki, R., Tajiri, T. *et al.* (2010) Gene expression levels as predictive markers of outcome in pancreatic cancer after gemcitabine-based adjuvant chemotherapy. *Neoplasia*, **12**, 807-817.
67. Hsieh, Y.Y., Chou, C.J., Lo, H.L. and Yang, P.M. (2016) Repositioning of a cyclin-dependent kinase inhibitor GW8510 as a ribonucleotide reductase M2 inhibitor to treat human colorectal cancer. *Cell Death Discov*, **2**, 16027.
68. Zhang, C., Aldrees, M., Arif, M., Li, X., Mardinoglu, A. and Aziz, M.A. (2019) Elucidating the Reprograming of Colorectal Cancer Metabolism Using Genome-Scale Metabolic Modeling. *Front Oncol*, **9**, 681.
69. Tomita, H., Tanaka, K., Tanaka, T. and Hara, A. (2016) Aldehyde dehydrogenase 1A1 in stem cells and cancer. *Oncotarget*, **7**, 11018-11032.
70. Krause, K., Caution, K., Badr, A., Hamilton, K., Saleh, A., Patel, K., Seveau, S., Hall-Stoodley, L., Hegazi, R., Zhang, X. *et al.* (2018) CASP4/caspase-11 promotes autophagosome formation in response to bacterial infection. *Autophagy*, **14**, 1928-1942.
71. Fink, S.P., Myeroff, L.L., Kariv, R., Platzer, P., Xin, B., Mikkola, D., Lawrence, E., Morris, N., Nosrati, A., Willson, J.K. *et al.* (2015) Induction of KIAA1199/CEMIP is associated with colon cancer phenotype and poor patient survival. *Oncotarget*, **6**, 30500-30515.
72. Avogaro, L., Querido, E., Dalachi, M., Jantsch, M.F., Chartrand, P. and Cusanelli, E. (2018) Live-cell imaging reveals the dynamics and function of single-telomere TERRA molecules in cancer cells. *RNA Biol*, 1-10.
73. Hong, M., Zhang, Z., Chen, Q., Lu, Y., Zhang, J., Lin, C., Zhang, F., Zhang, W., Li, X., Zhang, W. *et al.* (2019) IRF1 inhibits the proliferation and metastasis of colorectal cancer by suppressing the RAS-RAC1 pathway. *Cancer Manag Res*, **11**, 369-378.
74. Komatsu, Y., Christian, S.L., Ho, N., Pongnopparat, T., Licursi, M. and Hirasawa, K. (2015) Oncogenic Ras inhibits IRF1 to promote viral oncolysis. *Oncogene*, **34**, 3985-3993.
75. Ohsugi, T., Yamaguchi, K., Zhu, C., Ikenoue, T., Takane, K., Shinozaki, M., Tsurita, G., Yano, H. and Furukawa, Y. (2019) Anti-apoptotic effect by the suppression of IRF1 as a downstream of Wnt/beta-catenin signaling in colorectal cancer cells. *Oncogene*, **38**, 6051-6064.
76. Shao, L., Hou, W., Scharping, N.E., Vendetti, F.P., Srivastava, R., Roy, C.N., Menk, A.V., Wang, Y., Chauvin, J.M., Karukonda, P. *et al.* (2019) IRF1 Inhibits Antitumor Immunity through the Upregulation of PD-L1 in the Tumor Cell. *Cancer Immunol Res*, **7**, 1258-1266.
77. Duong, H.Q., Yi, Y.W., Kang, H.J., Bae, I., Jang, Y.J., Kwak, S.J. and Seong, Y.S. (2014) Combination of dasatinib and gemcitabine reduces the ALDH1A1 expression and the proliferation of gemcitabine-resistant pancreatic cancer MIA PaCa-2 cells. *Int J Oncol*, **44**, 2132-2138.

78. Ozden, O., Park, S.H., Kim, H.S., Jiang, H., Coleman, M.C., Spitz, D.R. and Gius, D. (2011) Acetylation of MnSOD directs enzymatic activity responding to cellular nutrient status or oxidative stress. *Aging (Albany NY)*, **3**, 102-107.
79. Grasmann, G., Smolle, E., Olschewski, H. and Leithner, K. (2019) Gluconeogenesis in cancer cells - Repurposing of a starvation-induced metabolic pathway? *Biochim Biophys Acta Rev Cancer*, **1872**, 24-36.
80. Xiang, J., Zhang, Y., Tuo, L., Liu, R., Gou, D., Liang, L., Chen, C., Xia, J., Tang, N. and Wang, K. (2019) Transcriptomic changes associated with PCK1 overexpression in hepatocellular carcinoma cells detected by RNA-seq. *Genes & Diseases*.
81. Leclerc, D., Pham, D.N., Levesque, N., Truongcao, M., Foulkes, W.D., Sapienza, C. and Rozen, R. (2017) Oncogenic role of PDK4 in human colon cancer cells. *Br J Cancer*, **116**, 930-936.
82. Aguirre-Portoles, C., Feliu, J., Reglero, G. and Ramirez de Molina, A. (2018) ABCA1 overexpression worsens colorectal cancer prognosis by facilitating tumour growth and caveolin-1-dependent invasiveness, and these effects can be ameliorated using the BET inhibitor apabetalone. *Mol Oncol*, **12**, 1735-1752.
83. Basu, N.K., Kole, L., Basu, M., Chakraborty, K., Mitra, P.S. and Owens, I.S. (2008) The major chemical-detoxifying system of UDP-glucuronosyltransferases requires regulated phosphorylation supported by protein kinase C. *J Biol Chem*, **283**, 23048-23061.
