## Supplementary Information for "Immunity Depletion, Telomere Imbalance, and Cancer-associated Metabolism Pathway Aberrations in Intestinal Mucosa upon Caloric Restriction"

**Supplementary Figure 1** | Epithelial and immune system subcellular network

**Supplementary Figure 2** | NK, dendritic, phagocytic, and antigen presenting cell functional annotations network

**Supplementary Figure 3** | Immune system, epithelial, and cancer-associated subset interaction networks

**Supplementary Figure 4** | Interferon-inducible GTPase paralogs

**Supplementary Figure 5** | A model of CR-induced mucosal cells and pathway responses in DM

**Supplementary Table 1** | Functional enrichments common between human and mouse

**Supplementary Table 2** | Cell cycle periodic genes regulated upon CR (CycleBase)

**Supplementary Table 3** | The 171 Immune system genes characterized using the Disease and Biological Function annotation tool from IPA

**Supplementary Table 4** | CR-responded cancer stem cell related genes

**Supplementary Table 5** | The 121 cancer-associated genes characterized using the Disease and Biological annotation tool from IPA

**Supplementary Table 6** | Selected up-regulated oncogenes and down-regulated tumor suppressors upon CR

**Supplementary Table 7** | Experimental validation targets

**Supplementary Table 8** | IPA generated enriched canonical pathways

**Supplementary Table 9** | IPA canonical pathway: Sirtuin Signaling Pathway

**Supplementary Table 10** | CR DEGs immune cell-type classification

**Supplementary Table 11** | Gene symbols associated with multiple probe sets

**Supplementary Figure 1. Epithelial and immune system subcellular network**

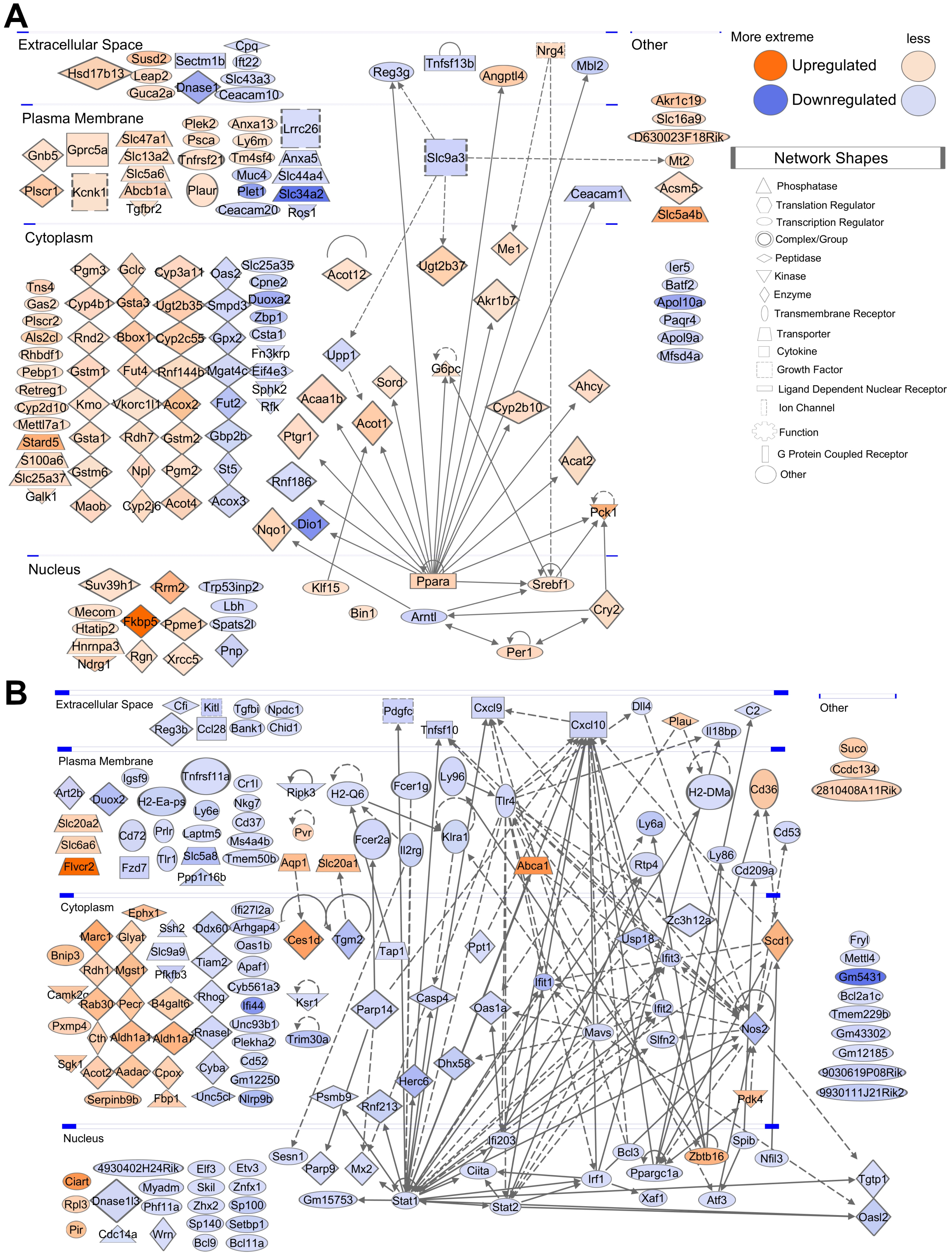


**(A)** Epithelial subcellular network including 29 connected and 118 non-connected ECGs from DM from DM distributed across four cellular compartments: extracellular space, plasma membrane, cytoplasm, nucleus. The ECG network contained 57 edges (protein interactions), average number of neighbors 2.28, clustering coefficient 0.06, network density 0.04, and protein-protein interaction (PPI) network enrichment $p < 1.00\ 10^{-16}$. The strongest hub for this network was Ppara (26 downstream, 4 upstream), which accounted for 26.3% (30/114) of the edges. **(B)** Immune system subcellular network including 68 connected and 103 non-connected ISGs. The ISG network contained 173 edges, average number of neighbors 3.72, clustering coefficient 0.18, network density 0.03, and PPI network enrichment $p < 1.00\ 10^{-16}$.The strongest hubs included Stat1 (36 downstream targets, 21 upstream targets), Tlr4 (19 downstream, 4 upstream), Stat2 (11 downstream, 5 upstream), Nos2 (10 downstream, 15 upstream), Ppargc1a (8 downstream, 6 upstream), and Cxcl10 (1 downstream, 16 upstream). Cellular locations of the gene hubs indicate Cxcl10 in the extracellular space, Tlr4 in the plasma membrane, Nos2 in the cytoplasm, and Stat1, Stat2, and Ppargc1a in the nucleus. In total, these genes account for 43.9% (152/356) of the edges.

**Supplementary Figure 2. NK, dendritic, phagocytic, and antigen presenting cell functional annotations network**

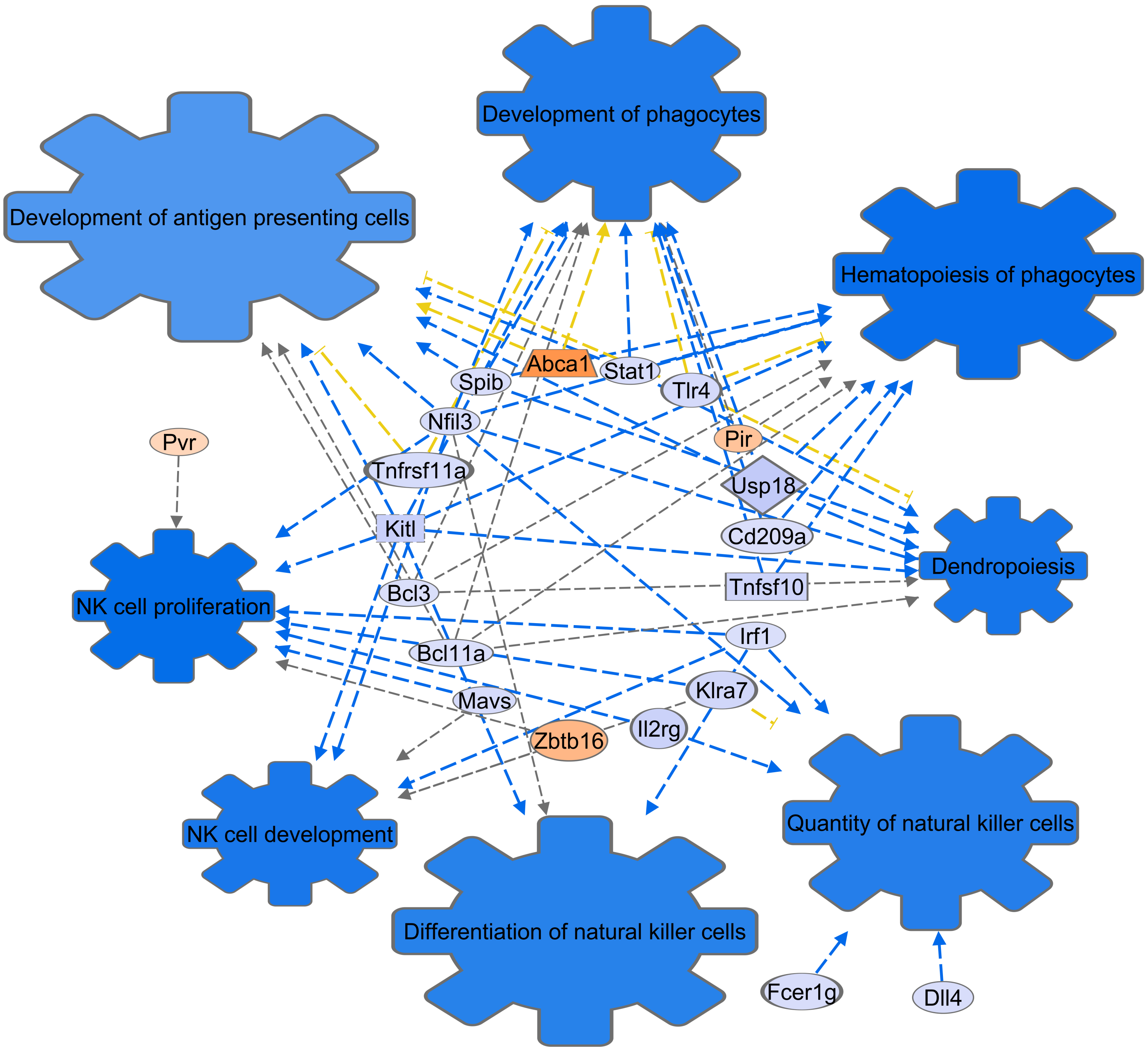


A network of 21 genes based on the functional annotations of the 171 Immune System Genes. NK, dendritic, phagocytic, and antigen presenting cell associations within the Lymphoid Structure and Development IPA category were selected including development of phagocytes (p = 1.29 $10^{-08}$, *n* = 13), hematopoiesis of phagocytes (p = 2.1 $10^{-08}$, *n* = 11), development of antigen presenting cells (p = 7.18 $10^{-07}$, *n* = 10), dendropoiesis (p = 2.19 $10^{-06}$, *n* = 8), NK cell proliferation (p = 7.31 $10^{-06}$, *n* = 8), NK cell development (p = 5.86 $10^{-05}$, *n* = 6), quantity of natural killer cells (p = 1.56 $10^{-03}$, *n* = 6), and differentiation of natural killer cells (p = 2.39 $10^{-03}$, *n* = 3). Additional T lymphocyte annotations of the CR-induced immune DEGs included lack of T lymphocytes (p = 2.93 $10^{-6}$, *n* = 5), abnormal morphology of T lymphocytes (p = 3.53 $10^{-5}$, *n* = 8), abnormal morphology of Peyer's patches (p = 8.54 $10^{-4}$, *n* = 4), and lack of gamma-delta T lymphocytes (p = 1.25 $10^{-3}$, *n* = 2). Additional B lymphocyte annotations included: abnormal morphology of B lymphocytes (p = 1.04 $10^{-3}$, *n* = 5) and morphology of B-cell follicle (p = 2.04 $10^{-3}$, *n* = 4). These T and B cell annotations were not included in Figure 4 due to lack of IPA predicted z-score information for activation or inhibition.

**Supplementary Figure 3. Immune system, epithelial, and cancer-associated subset network interactions**

**(A)** The cancer-associated and immune network contained 93 genes, 245 edges, average number of neighbors 3.63, clustering coefficient 0.16, network density 0.02, and PPI network enrichment $p < 1\ 10^{-16}$. **(B)** The cancer and epithelial network contained 68 genes, 186 edges, average number of neighbors 3.62, clustering coefficient 0.12, network density 0.03, and and PPI network enrichment $p < 1\ 10^{-16}$. Non-connected genes are not displayed. Blue indicates downregulated genes and orange indicates upregulated genes. Color intensity is based on the expression fold changes of the genes. Understanding the tumor-immune, immune-epithelial, and tumor-epithelial microenvironment interactions could explain CR malignancy risks and inform therapeutic strategies.

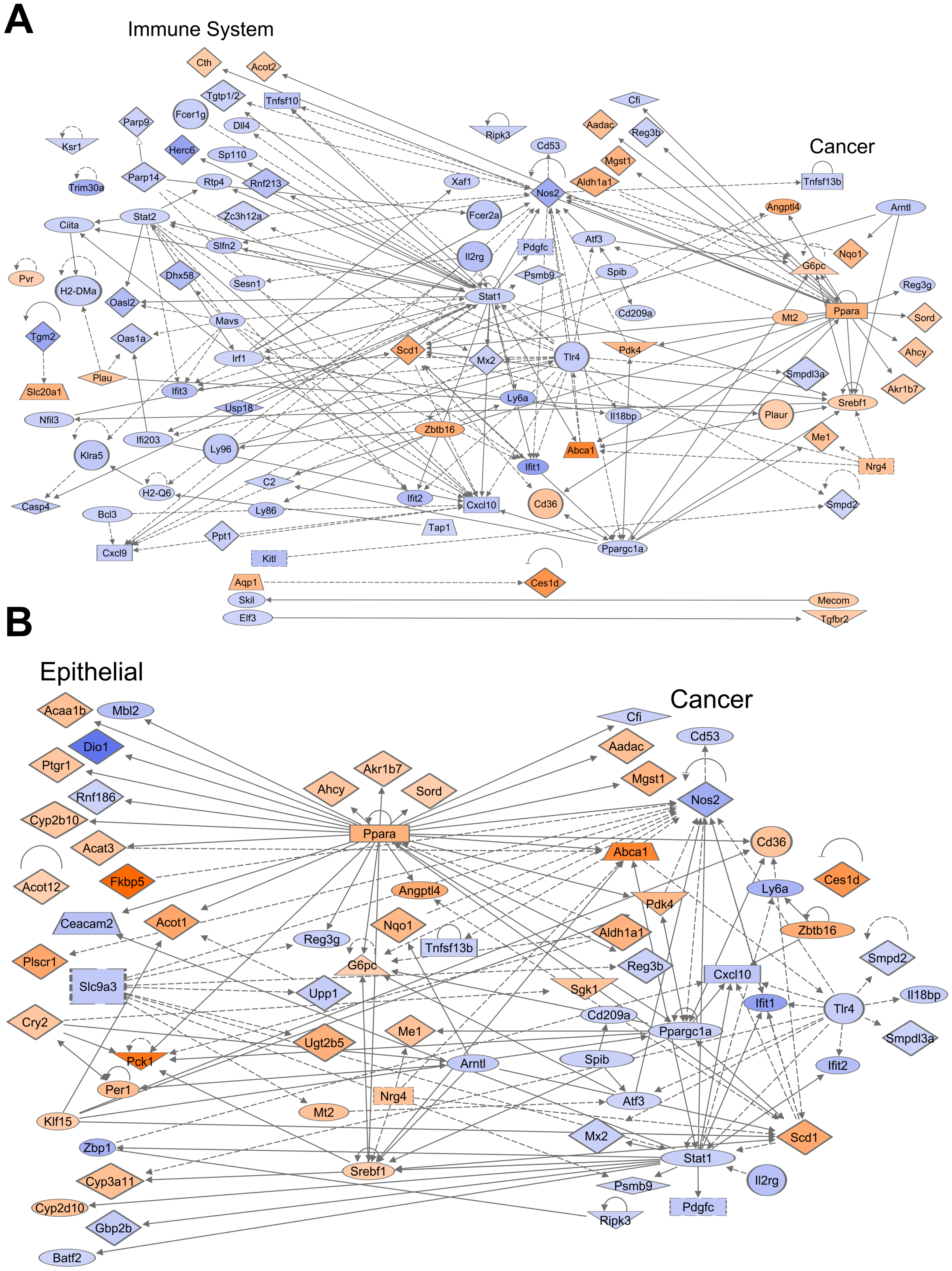

**Supplementary Figure 4. Interferon-inducible GTPase paralogs**

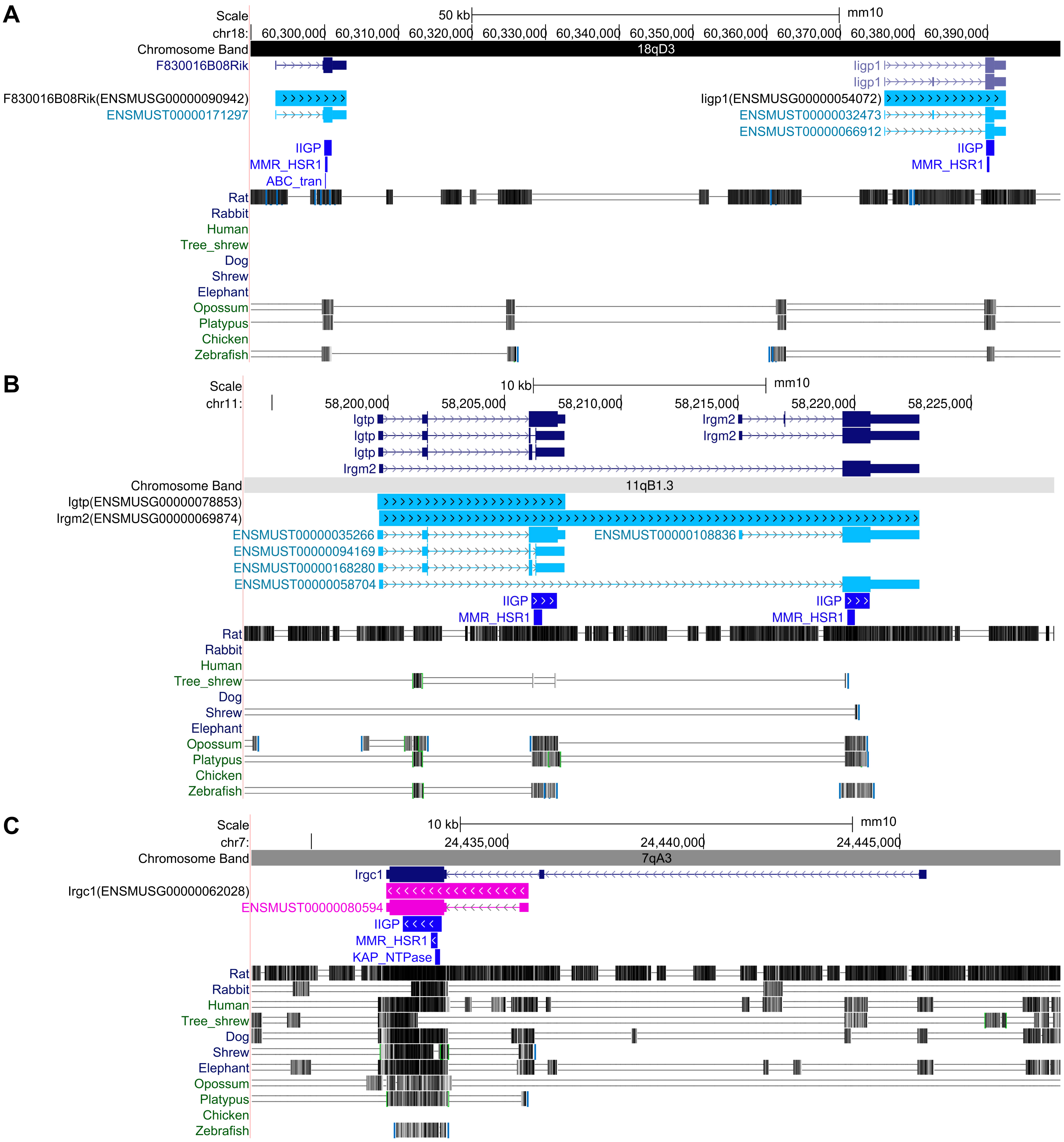


UCSC genome browser tracks defining four paralogs of the interferon-inducible GTPase family: **(A)** F830016B08Rik, Iigp1, **(B)** Irgm2, Igtp, **(C)** Irgc1. Annotation tracks are from Ensembl (Build 75) and RefSeq genome assembly (GRCm38.p5). Pfam-A domains are identified by software HMMER3: IIGP (Interferon-inducible GTPase; PF05049), MMR_HSR1(50S ribosome-binding GTPase; PF01926), ABC_tran (ABC transporter; PF00005), KAP_NTPase (KAP family P-loop domain; PF07693). The "Multiz alignments" track for mouse genome assembly (mm10) provided multiple alignment data combining PhyloP and PhastCons methods and shows the results of measurements of evolutionary conservation using these methods from the PHAST package for 60 vertebrates and three subsets (Glires, Euarchontoglires and placental mammal).

**Supplementary Figure 5. A model of CR-induced mucosal cells and pathway responses in DM**

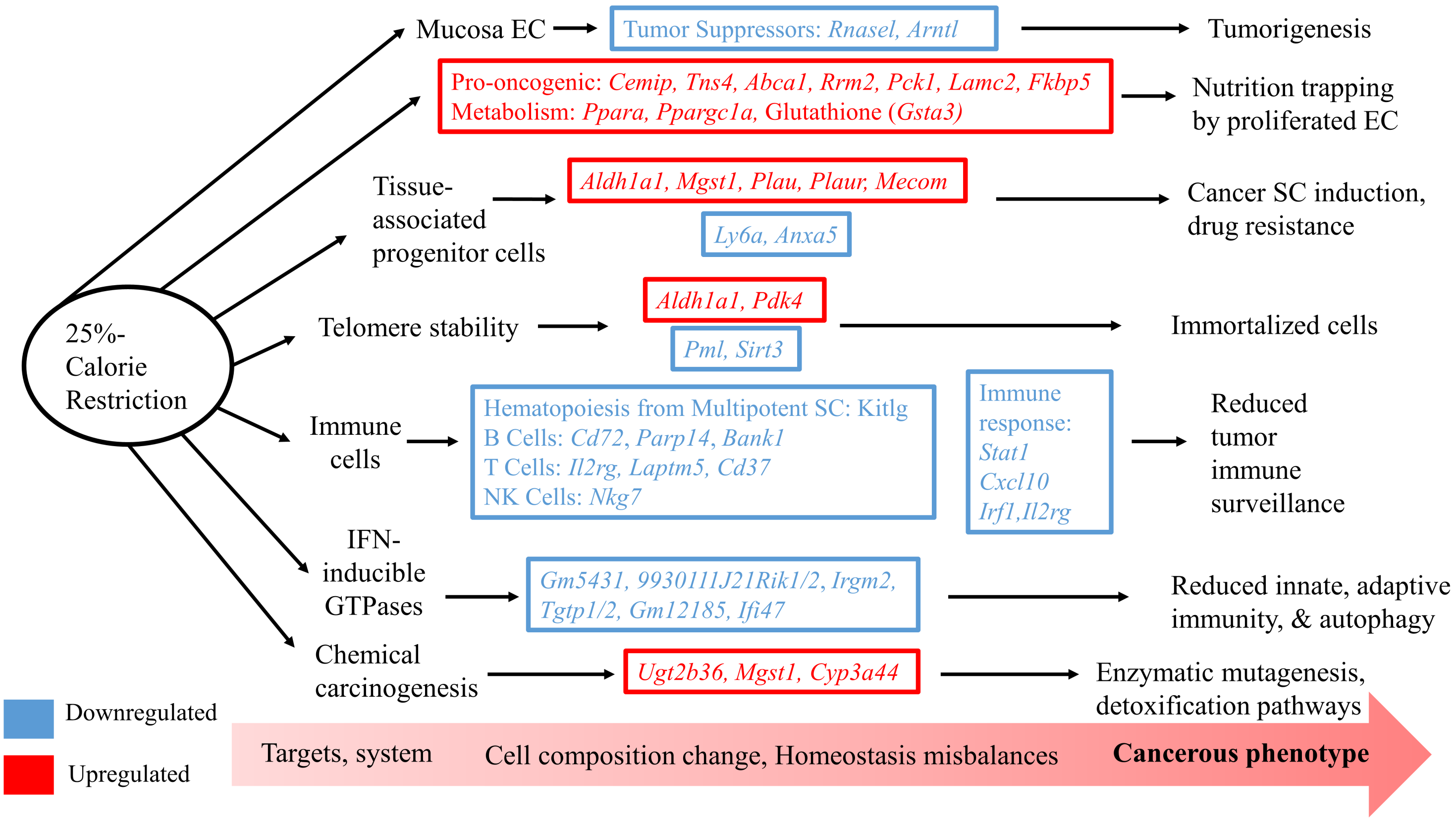


A more comprehensive proposed gene-specific model of CR-induced mucosal cells and pathway responses in metabolic reprogramming as illustrated in main text Figure 9. Note: Epithelial Cell (EC); Stem Cell (SC).

### Supplementary Table 1. Functional enrichments common between human and mouse

#### (A) GO Biological Process

| term ID | term description | human observed gene count | human background gene count | human FDR | mouse observed gene count | mouse background gene count | mouse FDR |
|---|---|---|---|---|---|---|---|
| GO:0009617 | response to bacterium | 34 | 555 | 9.41E-07 | 44 | 566 | 2.47E-13 |
| GO:0002376 | immune system process | 111 | 2370 | 1.01E-15 | 78 | 1703 | 1.42E-12 |
| GO:0006950 | response to stress | 112 | 3267 | 4.17E-08 | 103 | 2899 | 1.41E-10 |
| GO:0051607 | defense response to virus | 27 | 181 | 1.30E-12 | 21 | 152 | 5.17E-10 |
| GO:0051704 | multi-organism process | 80 | 2222 | 2.83E-06 | 74 | 1840 | 1.87E-09 |
| GO:0044281 | small molecule metabolic process | 79 | 1779 | 6.76E-10 | 61 | 1489 | 8.80E-08 |
| GO:0045087 | innate immune response | 44 | 676 | 1.32E-09 | 32 | 534 | 4.95E-07 |
| GO:0032787 | monocarboxylic acid metabolic process | 32 | 477 | 3.86E-07 | 26 | 426 | 9.48E-06 |
| GO:0071345 | cellular response to cytokine stimulus | 49 | 953 | 1.22E-07 | 33 | 676 | 2.12E-05 |
| GO:0006082 | organic acid metabolic process | 47 | 959 | 9.47E-07 | 36 | 794 | 2.84E-05 |
| GO:0005996 | monosaccharide metabolic process | 19 | 198 | 3.32E-06 | 14 | 148 | 8.03E-05 |
| GO:0031667 | response to nutrient levels | 27 | 455 | 3.88E-05 | 25 | 461 | 9.48E-05 |

#### (B) GO Molecular Function

| term ID | term description | human observed gene count | human background gene count | human FDR | mouse observed gene count | mouse background gene count | mouse FDR |
|---|---|---|---|---|---|---|---|
| GO:0043167 | ion binding | 154 | 6066 | 1.70E-03 | 141 | 5302 | 1.45E-06 |
| GO:0052689 | carboxylic ester hydrolase activity | 10 | 139 | 1.41E-02 | 13 | 132 | 1.00E-04 |
| GO:0015081 | sodium ion transmembrane transporter activity | 13 | 164 | 2.20E-03 | 13 | 138 | 1.50E-04 |
| GO:0016788 | hydrolase activity, acting on ester bonds | 31 | 713 | 2.20E-03 | 30 | 660 | 1.70E-04 |
| GO:0005102 | signaling receptor binding | 54 | 1513 | 9.60E-04 | 51 | 1515 | 2.80E-04 |
| GO:0016290 | palmitoyl-CoA hydrolase activity | 4 | 10 | 4.30E-03 | 5 | 14 | 8.40E-04 |
| GO:0004364 | glutathione transferase activity | 10 | 25 | 2.49E-07 | 6 | 28 | 1.20E-03 |
| GO:0000166 | nucleotide binding | 63 | 2097 | 7.60E-03 | 58 | 2006 | 2.60E-03 |
| GO:0008144 | drug binding | 53 | 1710 | 1.09E-02 | 48 | 1630 | 5.60E-03 |
| GO:0016740 | transferase activity | 68 | 2250 | 4.30E-03 | 56 | 2110 | 1.76E-02 |
| GO:0030246 | carbohydrate binding | 13 | 270 | 4.24E-02 | 13 | 265 | 1.91E-02 |

#### (C) KEGG Pathways

| human term ID | mouse term ID | term description | human observed gene count | human background gene count | human FDR | mouse observed gene count | mouse background gene count | mouse FDR |
|---|---|---|---|---|---|---|---|---|
| hsa00983 | mmu00983 | Drug metabolism - other enzymes | 17 | 76 | 3.08E-10 | 19 | 86 | 2.64E-12 |
| hsa01100 | mmu01100 | Metabolic pathways | 56 | 1250 | 1.71E-07 | 60 | 1296 | 5.33E-10 |
| hsa05204 | mmu05204 | Chemical carcinogenesis | 15 | 76 | 7.05E-09 | 15 | 92 | 2.53E-08 |
| hsa00480 | mmu00480 | Glutathione metabolism | 13 | 50 | 7.05E-09 | 11 | 61 | 1.20E-06 |
| hsa00830 | mmu00830 | Retinol metabolism | 7 | 62 | 3.80E-03 | 11 | 89 | 3.18E-05 |
| hsa01040 | mmu01040 | Biosynthesis of unsaturated fatty acids | 7 | 23 | 3.03E-05 | 7 | 27 | 3.78E-05 |
| hsa03320 | mmu03320 | PPAR signaling pathway | 10 | 72 | 7.57E-05 | 10 | 85 | 1.20E-04 |
| hsa05200 | mmu05200 | Pathways in cancer | 26 | 515 | 2.70E-04 | 25 | 522 | 1.50E-04 |
| hsa01524 | mmu01524 | Platinum drug resistance | 11 | 70 | 1.03E-05 | 9 | 76 | 2.70E-04 |

#### (D) Reactome Pathways

| human term ID | mouse term ID | term description | human observed gene count | human background gene count | human FDR | mouse observed gene count | mouse background gene count | mouse FDR |
|---|---|---|---|---|---|---|---|---|
| HSA-1430728 | MMU-1430728 | Metabolism | 99 | 2032 | 2.48E-15 | 79 | 1685 | 2.16E-13 |
| HSA-211859 | MMU-211859 | Biological oxidations | 28 | 214 | 5.36E-12 | 25 | 237 | 6.97E-10 |
| HSA-168256 | MMU-156590 | Immune System | 81 | 1925 | 2.87E-09 | 7 | 31 | 2.60E-04 |
| HSA-156590 | MMU-168256 | Glutathione conjugation | 11 | 33 | 5.80E-08 | 48 | 1523 | 1.90E-03 |
| HSA-168249 | MMU-168249 | Innate Immune System | 38 | 1012 | 4.30E-03 | 30 | 879 | 1.10E-02 |

Tables are ordered by individual mouse FDR (STRING v11, FDR <0.05).

### Supplementary Table 2. (A) Upregulated cell cycle periodic genes (CycleBase)

<table>
<tr><th>Human Gene Symbol</th><th>Mouse Gene Symbol</th><th>Ensemble ID</th><th>Gene Description</th><th>log(FC)</th><th>adj. P-value</th><th>Cell Cycle Phase</th><th>Literature (PubMed PMID)</th></tr>
<tr><td>FKBP5</td><td>Fkbp5</td><td>ENSMUSG00000024222</td><td>fk506 binding protein 5</td><td>8.54</td><td>9.35E-04</td><td>S</td><td>23936393, 21119664, 21498116</td></tr>
<tr><td colspan="8">FKBP5 could be a biomarker for tumourigenesis (through the AKT singalling pathway) and chemoresistance. High FKBP51 levels decreased AKT phosphorylation and increased chemosensitivity, whereas low FKBP51 levels increased AKT phosphorylation and decreased chemosensitivity. Downregulation of FKBP5 desensitized pancreatic and breast cancer cell lines to several different classes of chemotherapeutic agents. FKBP5 genetic variation influences response to gemcitabine treatment of pancreatic cancer.</td></tr>
<tr><td>RRM2</td><td>Rrm2</td><td>ENSMUSG00000020649</td><td>ribonucleotide reductase M2</td><td>4.02</td><td>2.05E-04</td><td>S</td><td>31363169, 20122995, 23113760, 20927319, 31417867, 17404105, 27551518</td></tr>
<tr><td colspan="8">Ribonucleoside-diphosphate reductase subunit M2, Rrm2, in mouse (RRM2 in human) encodes one of the small subunits of ribonucleotide reductase (RR), the rate limiting enzyme for production of deoxyribonucleotides. In both mouse and human cells this gene provides regulation of cell cycle processes and is activated by E2F1 and EZH2 DNA binding in gene promoter regions. RRM2 associates with RRM1 forming an active RR. Its accumulation plays a central role in providing precursors necessary for DNA synthesis catalyzing the biosynthesis of deoxyribonucleotides. RRM2 plays active roles in tumorigenesis and is a poor prognostic factor for cancers, such as colon, breast and pancreatic. It is involved in colorectal cancer metabolic reprogramming and siRNA knockdown of RRM2 in vitro and in vivo reduced cell proliferation.</td></tr>
<tr><td>ACSL3</td><td>Acsl3</td><td>ENSMUSG00000032883</td><td>acyl-CoA synthetase long-chain family member 3</td><td>2.11</td><td>6.33E-03</td><td>G1</td><td>30008815</td></tr>
<tr><td colspan="8">ACSL3 is associated with disease and especially with several cancers, can promote cancer cell survival through amplified fatty acid β-oxidation and increased arachidonic acid-dependent prostaglandin synthesis, both of which can drive tumor growth.</td></tr>
<tr><td>GAS2L3</td><td>Gas2l3</td><td>ENSMUSG00000074802</td><td>growth arrest-specific 2 like 3</td><td>1.81</td><td>2.16E-02</td><td>G2</td><td>22344256, 23469016, 24571573, 19139817</td></tr>
<tr><td colspan="8">Cytotoxic immunoconjugates which target and eliminate gastric cancer cells showed consistent downregulation responses of GAS2L3 in whole genome microarray expression profiling (validated using RTQ-PCR). GAS2L3 is necessary for proper cytokinesis and for abscission, the final step of cytokinesis that results in separation of the daughter cells. GAS2L3 is specifically expressed in mitosis and localizes to the spindle midzone/midbody and the constriction zone during cytokinesis. It mediates interactions with components of the so-called chromosome passenger complex consisting of the mitotic kinase Aurora B and associated proteins.</td></tr>
<tr><td>OPN3</td><td>Opn3</td><td>ENSMUSG00000026525</td><td>opsin 3</td><td>1.74</td><td>9.67E-03</td><td>G1</td><td>31802643, 29863164</td></tr>
<tr><td colspan="8">OPN3 acted as an oncogene through enhancing metastasis in lung adenocarcinoma via its overexpression promoting epithelial-mesenchymal transition. Blue light-emitting diodes (LED) irradiation and reductive effect on colon cancer cell viability was reversed by Opn3 siRNA knockdown.</td></tr>
<tr><td>RBBP8</td><td>Rbbp8</td><td>ENSMUSG00000041238</td><td>retinoblastoma binding protein 8</td><td>1.70</td><td>2.25E-02</td><td>S</td><td>31636387, 16249056</td></tr>
<tr><td colspan="8">RBBP8 facilitates the G1/S transition promoting Cyclin D1 and CDK4 levels and G2/M cycle checkpoints in double-stranded break repair processes. RBB8 is overexpressed in both gastric cancer and high-grade intraepithelial neoplasia tissues (HGIEN). Knockdown of RBBP8 inhibited cell proliferation and colony formation in gastric cancer. CtIP (also known as RBBP8) interacts with tumor suppressors, such as BRCA1. Ctip(-/-) embryo cells are arrested in G1 and do not enter S phase.</td></tr>
<tr><td>PUS7</td><td>Pus7</td><td>ENSMUSG00000057541</td><td>pseudouridylate synthase 7 homolog (S. cerevisiae)</td><td>1.64</td><td>2.43E-02</td><td>G1</td><td>31451225, 31433208, 30008265</td></tr>
<tr><td colspan="8">PUS7 may be a potential biomarker of glioma. It is involved in transcriptome reprogramming by cancer exosomes, which could lead to cancer-associated pathologies, immune evasion/modulation, and cell fate alteration/metastasis. PUS7 is an interactor of SIRT1, involved in the sirtuin signaling pathway.</td></tr>
<tr><td>ABCC5</td><td>Abcc5</td><td>ENSMUSG00000022822</td><td>ATP-binding cassette, sub-family C (CFTR/MRP), member 5</td><td>1.62</td><td>6.93E-03</td><td>S</td><td>15897250, 25640272</td></tr>
<tr><td colspan="8">Multidrug resistance proteins (MRPs) of the ABCC subfamily (including ABCC5) are involved in increased cellular efflux and resistance to fluoropyrimidine-based therapy. ABCC5 confers resistance to 5'-Fluorouracil (5-FU), used in the treatment of colon and breast cancers. Membrane transporters prevent anticancer drugs from reaching target intracellular concentrations. ABCC5 is an efflux transporter for cyclic nucleotides and nucleic acid analogs. Endogenous ABCC5 siRNA-mediated silencing increased 5-FU cellular cytotoxicity and enhanced accumulation of its metabolites.</td></tr>
<tr><td>KMO</td><td>Kmo</td><td>ENSMUSG00000039783</td><td>kynurenine 3-monooxygenase (kynurenine 3-hydroxylase)</td><td>1.51</td><td>4.32E-02</td><td>G1</td><td>32339939, 26099564</td></tr>
<tr><td colspan="8">Kynurenine 3-monooxygenase (KMO) is a pivotal enzyme of the kynerenine pathway (KP) involved in tryptophan (Trp) metabolism, which generates several toxic metabolites responsible for inflammatory disorders. KP enzymes are involved in cancer progression and may be a key biomarker for liver cancer. KMO positively regulated proliferation, migration, and invasion of human hepatocellular carcinoma (HCC). Kynurenin from cancer cells binds to the aryl hydrocarbon receptor (AHR) on T cells and suppresses T cell proliferation and oncolytic activities, attenuating anticancer immunity.</td></tr>
</table>

**Supplementary Table 2. (B) Downregulated cell cycle periodic genes (CycleBase)**

<table>
<tr><th>Human Gene Symbol</th><th>Mouse Gene Symbol</th><th>Ensemble ID</th><th>Gene Description</th><th>log(FC)</th><th>adj. P-value</th><th>Cell Cycle Phase</th><th>Literature (PubMed PMID)</th></tr>
<tr><td>IFIT1</td><td>Ifit1</td><td>ENSMUSG00000034459</td><td>interferon-induced protein with tetratricopeptide repeats 1</td><td>-3.36</td><td>2.57E-03</td><td>G1</td><td>31372503</td></tr>
<tr><td colspan="8">Functions in antiviral response. Ifit1 knockout promoted viral replication in murine norovirus infected cells.</td></tr>
<tr><td>TGM2</td><td>Tgm2</td><td>ENSMUSG00000037820</td><td>transglutaminase 2, C polypeptide</td><td>-3.10</td><td>4.48E-03</td><td>G1</td><td>31010374, 28754668, 31570702</td></tr>
<tr><td colspan="8">TGM2 is an apoptosis attenuator and related to cancer stem cell survival/tumor formation in multiple cancers. Its inhibition reversed mesenchymal transdifferentiation in glioma stem cells. TGM2 levels are upregulated in CRC and CRC patients with high levels of TGM2 had lower survival. TGM2-siRNA interference inhibited wnt3a/β-catenin/cyclin D1 pathway in colorectal cancer cells, attenuating tumor growth in nude mice. TGM2 is transcriptionally activated by ETS1, inhibits apoptosis, and is associated with chemotherapy stress in CRC via activation of Wnt/β-catenin signaling.</td></tr>
<tr><td>UNC5CL</td><td>Unc5cl</td><td>ENSMUSG00000043592</td><td>unc-5 homolog C (C. elegans)-like</td><td>-2.36</td><td>2.27E-04</td><td>G2</td><td>22158417, 30071397</td></tr>
<tr><td colspan="8">Unc5CL is a factor in epithelial inflammation and immunity. It a candidate gene involved in mucosal diseases, such as inflammatory bowel disease. Many chemokines (IL-8, CXCL1 and CCL20) are downstream targets of Unc5CL. These pro-inflammatory markers orchestrate the initial recruitment of immune cells and Unc5CL could be involved in epithelial danger responses. Unc5CL is a member of the family of death domain (DD)-containing proteins involved in many cellular processes, including apoptosis, inflammation, and development. Unc5CL is an inducer of proinflammatory signaling cascades leading to activation of NF-κB and JNK. It also has high specific tissue distribution in mucosal epithelia (including the intestine), and the protein is sorted to the apical face of these cells.</td></tr>
<tr><td>IFIT2</td><td>Ifit2</td><td>ENSMUSG00000045932</td><td>interferon-induced protein with tetratricopeptide repeats 2</td><td>-2.30</td><td>6.83E-03</td><td>G1</td><td>29316541</td></tr>
<tr><td colspan="8">IFIT2 knockdown significantly increased proliferation, migration, and invasion of human gastric cancer cell lines.</td></tr>
<tr><td>PAQR4</td><td>Paqr4</td><td>ENSMUSG00000023909</td><td>progestin and adipoQ receptor family member IV</td><td>-1.76</td><td>4.09E-04</td><td>S</td><td>29228296, 30322804, 28992327</td></tr>
<tr><td colspan="8">PAQR4 (Progestin and AdipoQ Receptor 4) expression is closely associated with progression of many cancers and microRNA (miRNA) processing. miR-370 inhibited cell proliferation, invasion and epithelial-to-mesenchymal transition (EMT) of gastric cancer (GC) cells by directly down-regulating PAQR4 expression. Knockdown of PAQR4 supressed cell proliferation in human breast cancer cells. PAQR4 is highly expressed in human breast cancers. The steady state of CDK4 (crucial for G1-to-S transition) is controlled by PAQR4. PAQR4 knockdown reduces cell proliferation accompanied by decreased CDK4 levels. PAQR4-deleted mice are resistant to chemical carcinogen-induced tumor formation.</td></tr>
<tr><td>STAT1</td><td>Stat1</td><td>ENSMUSG00000026104</td><td>signal transducer and activator of transcription 1</td><td>-1.66</td><td>2.37E-03</td><td>G2</td><td>22190859, 11323675, 12093877, 15891579, 30235866</td></tr>
<tr><td colspan="8">STAT1 can suppress tumor formation. The elimination of interferon (IFN)-γ, or STAT1 genes (thus lacking interferon-mediated pathways) in mice resulted in increased incidence and growth of spontaneous and chemically-induced tumors. Deficiency in STAT1 signaling predisposes gut inflammation and prompts colorectal cancer development</td></tr>
<tr><td>CDKL5</td><td>Cdkl5</td><td>ENSMUSG00000031292</td><td>cyclin-dependent kinase-like 5</td><td>-1.66</td><td>1.57E-02</td><td>S</td><td>31858726, 17589290, 28740074</td></tr>
<tr><td colspan="8">Cyclin-dependent kinase-like 5 (CDKL5) is highly expressed in gliomas, and CDKL5 overexpression promotes invasion, proliferation, migration and drug resistance of glioma cells. CDKL5 acts through the phosphoinositide 3-kinase (PI3K)/AKT axis. CDKL5 is also a potential target for immuotherapy in adult T-cell leukemia. CDKL5 localizes at the centrosome and at the midbody in proliferating cells and is required for faithful cell division. Acute inactivation of CDKL5 by RNA interference (RNAi) leads to multipolar spindle formation, cytokinesis failure and centrosome accumulation.</td></tr>
<tr><td>IER5</td><td>Ier5</td><td>ENSMUSG00000056708</td><td>immediate early response 5</td><td>-1.54</td><td>1.28E-03</td><td>S</td><td>28430589, 29104487, 26754925</td></tr>
<tr><td colspan="8">The immediate early response gene 5 (IER5) is a radiation response gene involved in DNA damage and repair. IER5 induced by radiation dose enhanced apoptosis of cervical cancer, was inversely associated with tumor size. IER5 is upregulated in several cancers and contributes to proliferation of cells under stressed conditions.</td></tr>
</table>

**Supplementary Table 3. The 171 Immune system genes characterized using the Disease and Biological Function annotation tool from IPA**

**(A)**

| Category | p-value | Molecules |
|---|---|---|
| Endocrine System Disorders | 1.1E-38-2.66E-04 | 59 |
| Gastrointestinal Disease | 1.1E-38-2.9E-03 | 80 |
| Immunological Disease | 1.1E-38-2.48E-03 | 73 |
| Metabolic Disease | 1.1E-38-2.7E-03 | 60 |
| Cellular Growth and Proliferation | 4.07E-19-2.39E-03 | 64 |
| **Lymphoid Tissue Structure and Development** | 4.07E-19-2.94E-03 | 66 |
| Infectious Diseases | 1.26E-17-2.94E-03 | 37 |
| Cell-To-Cell Signaling and Interaction | 2.47E-16-2.57E-03 | 47 |
| Immune Cell Trafficking | 2.47E-16-2.94E-03 | 51 |
| **Inflammatory Response** | 2.47E-16-2.94E-03 | 77 |
| Antimicrobial Response | 1.62E-14-6.6E-04 | 21 |
| Tissue Morphology | 7.65E-13-2.92E-03 | 58 |
| Cellular Function and Maintenance | 1.87E-12-2.39E-03 | 55 |
| Cell Death and Survival | 8.81E-12-2.94E-03 | 68 |
| Connective Tissue Disorders | 1.19E-09-1.23E-03 | 24 |
| Inflammatory Disease | 1.19E-09-2.3E-03 | 46 |

**(A)** Disease and Biological category annotations for the 171 Immune system gene subset. P-values indicate category dynamical range (all p<0.01), calculated by Fisher's Exact Test. **(B)** Disease annotations for the IPA Lymphoid Tissue Structure and Development category. **(C)** Disease annotations for the IPA Inflammatory Response category.

**(B)**

| Diseases or Functions Annotation | p-value | Activation z-score | # Molecules |
|---|---|---|---|
| **B cell annotations** | | | |
| Proliferation of B lymphocytes | 9.52E-10 | -1.033 | 18 |
| Quantity of B lymphocytes | 3.58E-06 | -0.257 | 17 |
| Proliferation of pre-B lymphocytes | 4.99E-04 | -0.837 | 4 |
| Abnormal morphology of B lymphocytes | 1.04E-03 | | 5 |
| Proliferation of pro-B lymphocytes | 1.73E-03 | | 3 |
| Differentiation of B lymphocytes | 1.90E-03 | | 8 |
| Morphology of B-cell follicle | 2.04E-03 | | 4 |
| **T cell annotations** | | | |
| Cell proliferation of T lymphocytes | 1.20E-09 | 0.88 | 25 |
| T cell development | 5.12E-09 | -1.255 | 24 |
| Quantity of T lymphocytes | 3.66E-07 | -0.084 | 23 |
| Lack of T lymphocytes | 2.93E-06 | | 5 |
| Differentiation of T lymphocytes | 3.94E-06 | -1.545 | 16 |
| Quantity of CD8+ T lymphocyte | 1.53E-05 | -1.156 | 10 |
| Abnormal morphology of T lymphocytes | 3.53E-05 | | 8 |
| Quantity of helper T lymphocytes | 1.99E-04 | 1.379 | 7 |
| Quantity of CD4+ T-lymphocytes | 2.14E-04 | -0.042 | 9 |
| Quantity of intraepithelial T lymphocytes | 3.99E-04 | | 3 |
| Differentiation of Th2 cells | 6.16E-04 | 1.4 | 5 |
| Quantity of double-negative T lymphocyte | 6.96E-04 | -1.387 | 5 |
| Abnormal morphology of Peyer's patches | 8.54E-04 | | 4 |
| Lack of gamma-delta T lymphocytes | 1.25E-03 | | 2 |
| Quantity of natural killer T lymphocytes | 2.92E-03 | | 4 |
| **NK, phagocytic, dendritic, and antigen presenting cell annotations** | | | |
| Development of phagocytes | 1.29E-08 | -1.576 | 13 |
| Hematopoiesis of phagocytes | 2.10E-08 | -2.208 | 11 |
| Development of antigen presenting cells | 7.18E-07 | -1.035 | 10 |
| Dendropoiesis | 2.19E-06 | -1.756 | 8 |
| NK cell proliferation | 7.31E-06 | -2.393 | 8 |
| NK cell development | 5.86E-05 | | 6 |
| Quantity of natural killer cells | 1.56E-03 | -1.467 | 6 |
| Differentiation of natural killer cells | 2.39E-03 | | 3 |

B Cell genes:

Bank1,Bcl11a,Bcl3,Ccl28,Cd209a, Cd36,Cd72,Dll4,Fcer1g,Fcer2a,Il2rg, Kitl,Ly86,Ly96,Mavs,Nfil3,Nos2, Parp14,Plekha2,Rhog,Ripk3,Spib, Stat1,Tgfbi,Tlr4,Tnfrsf11a,Unc93b1, Zc3h12a

T Cell genes:

Apaf1,Art2b,Bcl11a,Bcl3,Camk2g, Ccdc134,Cd209a,Cd37,Ciita,Cr1l, Cth,Cxcl10,Dll4,Elf3,Fcer1g,Fzd7, H2-DMa,H2-Q6,Il2rg,Irf1,Kitl,Klra7, Ksr1, Laptm5, Ly6a,Mavs,Nfil3, Nos2,Plau,Psmb9,Pvr,Ripk3,Skil, Slc6a6,Spib, Stat1,Stat2,Tap1,Tlr4, Tnfrsf11a,Tnfsf10,Trim30a,Usp18, Zbtb16,Zc3h12a

NK genes:

Dll4,Fcer1g,Il2rg,Irf1,Kitl,Klra7, Mavs, Nfil3,Pvr,Zbtb16

**(C)**

| Diseases or Functions Annotation | p-value | Activation z-score | # Molecules |
|---|---|---|---|
| Activation of leukocytes | 2.5E-16 | -0.544 | 32 |
| Antimicrobial response | 1.6E-14 | | 20 |
| Antiviral response | 5.3E-14 | | 16 |
| Inflammation of absolute anatomical region | 1.3E-11 | 1.594 | 37 |
| Inflammatory response | 3.7E-09 | -1.326 | 26 |
| Activation of macrophages | 8.4E-08 | -0.336 | 12 |
| Activation of phagocytes | 1.1E-07 | -0.342 | 14 |

Inflammatory response genes:

Atf3,Ccl28,Cd209a,Cd36,Cd37,Ciita, Cr1l,Cxcl10,Cxcl9,Fcer1g,Kitl,Mavs, Nos2,Pdk4,Pfkfb3,Plau,Ppt1,Reg3a, Ripk3,Rnasel,Stat1,Tgm2,Tlr4, Tnfrsf11a,Usp18,Zbtb16

**Supplementary Table 4. CR-responded cancer stem cell related genes**

**(A)**

| Ensemble ID | Gene Symbol | Gene Description | log(FC) | adj. P-value |
|---|---|---|---|---|
| ENSMUSG00000020826 | Nos2 | nitric oxide synthase 2, inducible | -2.92 | 1.39E-02 |
| ENSMUSG00000075602 | Ly6a | lymphocyte antigen 6 complex, locus A | -2.67 | 1.05E-03 |
| ENSMUSG00000019966 | Kitl | kit ligand | -2.25 | 8.00E-03 |
| ENSMUSG00000031304 | Il2rg | interleukin 2 receptor, gamma chain | -2.18 | 1.62E-04 |
| ENSMUSG00000028019 | Pdgfc | platelet-derived growth factor, C polypeptide | -1.96 | 3.73E-03 |
| ENSMUSG00000027712 | Anxa5 | annexin A5 | -1.80 | 3.84E-02 |
| ENSMUSG00000039005 | Tlr4 | toll-like receptor 4 | -1.77 | 1.23E-03 |
| ENSMUSG00000027314 | Dll4 | delta-like 4 (Drosophila) | -1.61 | 1.31E-02 |
| ENSMUSG00000024371 | C2 | complement component 2 (within H-2S) | -1.59 | 1.99E-02 |
| ENSMUSG00000071757 | Zhx2 | zinc fingers and homeoboxes 2 | -1.59 | 1.56E-03 |

**(B)**

| Ensemble ID | Gene Symbol | Gene Description | log(FC) | adj. P-value |
|---|---|---|---|---|
| ENSMUSG00000046223 | Plaur | plasminogen activator, urokinase receptor | 1.55 | 2.57E-02 |
| ENSMUSG00000027684 | Mecom | MDS1 and EVI1 complex locus | 1.67 | 2.99E-02 |
| ENSMUSG00000021822 | Plau | plasminogen activator, urokinase | 1.78 | 1.48E-03 |
| ENSMUSG00000078566 | Bnip3 | BCL2/adenovirus E1B interacting protein 3 | 2.20 | 8.95E-03 |
| ENSMUSG00000040584 | Abcb1a | ATP-binding cassette, sub-family B (MDR/TAP), member 1A | 2.34 | 2.27E-04 |
| ENSMUSG00000008540 | Mgst1 | microsomal glutathione S-transferase 1 | 2.37 | 3.45E-04 |
| ENSMUSG00000053279 | Aldh1a1 | aldehyde dehydrogenase family 1, subfamily A1 | 2.45 | 1.76E-04 |
| ENSMUSG00000005125 | Ndrg1 | N-myc downstream regulated gene 1 | 2.65 | 1.48E-03 |

**(A)** Downregulated cancer stem cell related genes differentially regulated upon CR. **(B)** Upregulated cancer stem cell related genes differentially regulated upon CR. *Aldh1a1* was identified as both a cancer stem cell related gene and marker.

**Supplementary Table 5. The 121 cancer-associated genes characterized using the Disease and Biological annotation tool from IPA**

**(A)**

| Category | p-value | Molecules |
|---|---|---|
| Metabolic Disease | 4.17E-13-1.21E-03 | 38 |
| **Cancer** | 7.03E-12-2.64E-03 | 48 |
| Humoral Immune Response | 4.79E-11-2.98E-03 | 24 |
| Gastrointestinal Disease | 1.63E-10-2.86E-03 | 55 |
| Endocrine System Disorders | 2.1E-10-1.21E-03 | 32 |
| Digestive System Development and Function | 4.47E-10-1.6E-03 | 28 |
| Immunological Disease | 5.39E-10-2.86E-03 | 33 |
| Lipid Metabolism | 1.47E-09-2.73E-03 | 33 |
| Small Molecule Biochemistry | 1.47E-09-2.73E-03 | 37 |
| Lymphoid Tissue Structure and Development | 9.41E-09-2.49E-03 | 34 |
| Immune Cell Trafficking | 1.08E-08-2.64E-03 | 32 |
| Carbohydrate Metabolism | 1.25E-08-1.18E-03 | 24 |
| Inflammatory Response | 4.82E-08-2.64E-03 | 53 |
| Cellular Function and Maintenance | 2.75E-07-2.29E-03 | 40 |
| Infectious Diseases | 5.68E-07-1.18E-03 | 16 |

Metabolic Disease:

Abca1,Akr1b7,Angptl4,Arntl,Atf3,Cd209a,Cd36, Cxcl10,Ddx60,Fcer2a,G6pc,Gsta1,Ifit1,Ifit2, Igsf9,Il18bp,Ly6a,Maob,Me1,Mgst1,Nos2,Nqo1, Nrg4,Pdgfc,Ppara,Ppargc1a,Psmb9,Scd1,Spib, Srebf1,Stat1,Tap1,Tgfbr2,Tlr4,Tnfrsf21, Tnfsf13b,Upp1,Xaf1

Cancer:

Abca1,Angptl4,Atf3,Cd36,Cyba,Cyp2c55,Fut4, Galk1,Guca2a,Htatip2,Ifit1,Il18bp,Il2rg,Lamc2, Ly6a,Mecom,Mt2,Muc4,Ndrg1,Nos2,Nqo1, Nuak2,Pdgfc,Pdk4,Plaur,Ppara,Ppargc1a,Psmb9, Reg3a,Ripk3,Rpl3,Rrm2,Scd1,Setbp1,Slc6a6, Smpd3,Spib,Srebf1,Stat1,Tap1,Tff1,Tgfbr2,Tlr4, Tnfrsf21, Tnfsf13b,Xaf1,Xrcc5,Zbtb16

**(B)**

| Diseases or Functions Annotation | p-value | Activation z-score | Molecules | # Molecules |
|---|---|---|---|---|
| Abdominal neoplasm | 7.03E-12 | -0.594 | Atf3,Cyp2c55,Fut4,Guca2a,Htatip2,Ly6a,Mt2,Muc4,Ndrg1,Nos2, Nuak2,Pdgfc,Ppara,Psmb9,Ripk3,Rrm2,Scd1,Tap1,Tff1,Tgfbr2,Tlr4, Xaf1,Xrcc5 | 23 |
| Digestive organ tumor | 1.63E-10 | -0.943 | Cyp2c55,Guca2a,Htatip2,Ly6a,Mt2,Muc4,Ndrg1,Nos2,Nuak2,Pdgfc, Ppara,Psmb9,Ripk3,Rrm2,Scd1,Stat1,Tap1,Tff1,Tgfbr2,Tlr4 | 21 |
| Abdominal cancer | 3.00E-10 | -0.184 | Atf3,Cyp2c55,Fut4,Htatip2,Ly6a,Mt2,Muc4,Ndrg1,Nos2,Pdgfc,Ppara, Psmb9,Ripk3,Rrm2,Scd1,Tgfbr2,Xaf1 | 17 |
| Liver cancer | 7.57E-08 | -0.512 | Cyp2c55,Htatip2,Mt2,Ndrg1,Nos2,Pdgfc,Ppara,Psmb9,Rrm2,Scd1, Tgfbr2 | 11 |
| Abdominal carcinoma | 7.96E-08 | 0.152 | Tgfbr2,Scd1,Rrm2,Ripk3,Psmb9,Ppara,Pdgfc,Nos2,Mt2,Ly6a,Htatip2, Cyp2c55 | 12 |
| Epithelial neoplasm | 2.40E-07 | 0.585 | Angptl4,Atf3,Cyp2c55,Htatip2,Ifit1,Ly6a,Mt2,Nos2,Nqo1,Nuak2, Pdgfc,Ppara,Psmb9,Reg3a,Ripk3,Rrm2,Scd1,Stat1,Tff1,Tgfbr2 | 20 |
| Tumorigenesis of epithelial neoplasm | 1.08E-04 | 0.317 | Htatip2,Ifit1,Ly6a,Mt2,Nqo1,Nuak2,Pdgfc,Ppara,Psmb9,Rrm2,Stat1, Tff1,Tgfbr2 | 13 |

**(A)** Disease and Biological category annotations for the 121 cancer-associated gene subset. P-values indicate category dynamical range (all p<0.01), calculated by Fisher's Exact Test. **(B)** Disease annotations for the IPA Cancer category.

**Supplementary Table 6. Selected up-regulated oncogenes and down-regulated tumor suppressors upon CR**

| Gene Symbol | log(FC) | Adjusted P-value | Cancer relevance |
|---|---|---|---|
| Pck1 | 4.39 | 1.62E-04 | 1, 2 |
| Aldh1a1 | 2.45 | 1.76E-04 | 3 |
| Rrm2 | 4.02 | 2.05E-04 | 4 |
| Cemip | 4.25 | 2.27E-04 | 5 |
| Tns4 | 2.27 | 1.28E-03 | 6, 7, 8 |
| Abca1 | 3.90 | 1.66E-03 | 9 |
| Gsta3 | 3.00 | 2.04E-03 | 10, 11 |
| Rab30 | 2.32 | 1.22E-02 | 12, 13 |
| Sgk1 | 1.86 | 3.90E-02 | 14 |
| Rnasel | -1.74 | 2.37E-03 | 15, 16, 17, 18, 19, 20, 21, 22 |
| Irf1 | -1.54 | 2.43E-03 | 23, 24, 25, 26, 27 |
| Arntl | -1.87 | 1.56E-02 | 28, 29, 30, 31 |


1. Grasmann G, Smolle E, Olschewski H, Leithner K. Gluconeogenesis in cancer cells - Repurposing of a starvation-induced metabolic pathway? Biochim Biophys Acta Rev Cancer 1872, 24-36 (2019).
2. Xiang J, et al. Transcriptomic changes associated with PCK1 overexpression in hepatocellular carcinoma cells detected by RNA-seq. Genes & Diseases, (2019).
3. Tomita H, Tanaka K, Tanaka T, Hara A. Aldehyde dehydrogenase 1A1 in stem cells and cancer. Oncotarget 7, 11018-11032 (2016).
4. Liu X, et al. Ribonucleotide reductase small subunit M2 serves as a prognostic biomarker and predicts poor survival of colorectal cancers. Clin Sci (Lond) 124, 567-578 (2013).
5. Fink SP, et al. Induction of KIAA1199/CEMIP is associated with colon cancer phenotype and poor patient survival. Oncotarget 6, 30500-30515 (2015).
6. Tang X, et al. A mechanically-induced colon cancer cell population shows increased metastatic potential. Mol Cancer 13, 131 (2014).
7. Kim S, Kim N, Kang K, Kim W, Won J, Cho J. Whole Transcriptome Analysis Identifies TNS4 as a Key Effector of Cetuximab and a Regulator of the Oncogenic Activity of KRAS Mutant Colorectal Cancer Cell Lines. Cells 8, (2019).
8. Sawazaki S, et al. Clinical Significance of Tensin 4 Gene Expression in Patients with Gastric Cancer. In Vivo 31, 1065-1071 (2017).
9. Aguirre-Portoles C, Feliu J, Reglero G, Ramirez de Molina A. ABCA1 overexpression worsens colorectal cancer prognosis by facilitating tumour growth and caveolin-1-dependent invasiveness, and these effects can be ameliorated using the BET inhibitor apabetalone. Mol Oncol 12, 1735-1752 (2018).
10. Xiao Y, Meierhofer D. Glutathione Metabolism in Renal Cell Carcinoma Progression and Implications for Therapies. Int J Mol Sci 20, (2019).
11. Al Ahmad A, et al. Papillary Renal Cell Carcinomas Rewire Glutathione Metabolism and Are Deficient in Both Anabolic Glucose Synthesis and Oxidative Phosphorylation. Cancers (Basel) 11, (2019).
12. Regnier M, et al. Insights into the role of hepatocyte PPARalpha activity in response to fasting. Mol Cell Endocrinol 471, 75-88 (2018).
13. Vidyasekar P, et al. Genome Wide Expression Profiling of Cancer Cell Lines Cultured in Microgravity Reveals Significant Dysregulation of Cell Cycle and MicroRNA Gene Networks. PLoS One 10, e0135958 (2015).
14. Lang F, Perrotti N, Stournaras C. Colorectal carcinoma cells--regulation of survival and growth by SGK1. Int J Biochem Cell Biol 42, 1571-1575 (2010).
15. Long TM, et al. RNase-L deficiency exacerbates experimental colitis and colitis-associated cancer. Inflamm Bowel Dis 19, 1295-1305 (2013).
16. Burke JM, Lester ET, Tauber D, Parker R. RNase L promotes the formation of unique ribonucleoprotein granules distinct from stress granules. J Biol Chem 295, 1426-1438 (2020).
17. Burke JM, Moon SL, Matheny T, Parker R. RNase L Reprograms Translation by Widespread mRNA Turnover Escaped by Antiviral mRNAs. Mol Cell 75, 1203-1217 e1205 (2019).
18. Kruger S, et al. The additive effect of p53 Arg72Pro and RNASEL Arg462Gln genotypes on age of disease onset in Lynch syndrome patients with pathogenic germline mutations in MSH2 or MLH1. Cancer Lett 252, 55-64 (2007).
19. Kruger S, et al. Arg462Gln sequence variation in the prostate-cancer-susceptibility gene RNASEL and age of onset of hereditary non-polyposis colorectal cancer: a case-control study. Lancet Oncol 6, 566-572 (2005).
20. Batman G, Oliver AW, Zehbe I, Richard C, Hampson L, Hampson IN. Lopinavir up-regulates expression of the antiviral protein ribonuclease L in human papillomavirus-positive cervical carcinoma cells. Antivir Ther 16, 515-525 (2011).
21. Banerjee S, et al. OAS-RNase L innate immune pathway mediates the cytotoxicity of a DNA-demethylating drug. Proc Natl Acad Sci U S A 116, 5071-5076 (2019).
22. Banerjee S, et al. RNase L is a negative regulator of cell migration. Oncotarget 6, 44360-44372 (2015).
23. Zenke K, Muroi M, Tanamoto KI. IRF1 supports DNA binding of STAT1 by promoting its phosphorylation. Immunol Cell Biol 96, 1095-1103 (2018).
24. Hong M, et al. IRF1 inhibits the proliferation and metastasis of colorectal cancer by suppressing the RAS-RAC1 pathway. Cancer Manag Res 11, 369-378 (2019).
25. Komatsu Y, Christian SL, Ho N, Pongnopparat T, Licursi M, Hirasawa K. Oncogenic Ras inhibits IRF1 to promote viral oncolysis. Oncogene 34, 3985-3993 (2015).
26. Ohsugi T, et al. Anti-apoptotic effect by the suppression of IRF1 as a downstream of Wnt/beta-catenin signaling in colorectal cancer cells. Oncogene 38, 6051-6064 (2019).
27. Shao L, et al. IRF1 Inhibits Antitumor Immunity through the Upregulation of PD-L1 in the Tumor Cell. Cancer Immunol Res 7, 1258-1266 (2019).
28. Gwon DH, et al. BMAL1 Suppresses Proliferation, Migration, and Invasion of U87MG Cells by Downregulating Cyclin B1, Phospho-AKT, and Metalloproteinase-9. Int J Mol Sci 21, (2020).
29. Tang Q, et al. Circadian Clock Gene Bmal1 Inhibits Tumorigenesis and Increases Paclitaxel Sensitivity in Tongue Squamous Cell Carcinoma. Cancer Res 77, 532-544 (2017).
30. Fekry B, et al. Incompatibility of the circadian protein BMAL1 and HNF4alpha in hepatocellular carcinoma. Nat Commun 9, 4349 (2018).
31. Yeh CM, et al. Epigenetic silencing of ARNTL, a circadian gene and potential tumor suppressor in ovarian cancer. Int J Oncol 45, 2101-2107 (2014).

### Supplementary Table 7. Experimental validation targets

| Ensembl ID | Gene Symbol | Gene Description | Categorization | Potential Therapeutic Target | log(FC) | adj. P-value |
|---|---|---|---|---|---|---|
| ENSMUSG00000020826 | Nos2 | nitric oxide synthase 2, inducible | cancer-associated/Sirtuin Signaling Pathway | yes | -2.92 | 1.39E-02 |
| ENSMUSG00000039005 | Tlr4 | toll-like receptor 4 | cancer-associated | yes | -1.77 | 1.23E-03 |
| ENSMUSG00000034855 | Cxcl10 | chemokine (C-X-C motif) ligand 10 | cancer-associated | yes | -2.03 | 3.80E-03 |
| ENSMUSG00000036986 | Pml | promyelocytic leukemia | telomeric | | -1.43 | 3.57E-02 |
| ENSMUSG00000029167 | Ppargc1a | peroxisome proliferative activated receptor, gamma, coactivator 1 alpha | metabolism/cancer-associated/Sirtuin Signaling Pathway | | -1.65 | 1.22E-02 |
| ENSMUSG00000027513 | Pck1 | phosphoenolpyruvate carboxykinase 1, cytosolic | Sirtuin Signaling Pathway | | 4.39 | 1.62E-04 |
| ENSMUSG00000015243 | Abca1 | ATP-binding cassette, sub-family A (ABC1), member 1 | Sirtuin Signaling Pathway/cancer cell metabolism | | 3.90 | 1.66E-03 |
| ENSMUSG00000025934 | Gsta3 | glutathione S-transferase, alpha 3 | glutathione metabolism | | 3.00 | 2.04E-03 |
| ENSMUSG00000020649 | Rrm2 | ribonucleotide reductase M2 | cancer-associated, cell cycle, glutathione metabolism | yes | 4.02 | 2..05E-04 |
| ENSMUSG00000026479 | Lamc2 | laminin, gamma 2 | cancer-associated | | 1.75 | 6.78E-03 |
| ENSMUSG00000021822 | Plau | plasminogen activator, urokinase | cancer-associated | | 1.78 | 1.48E-03 |

Indicates top genes for future qPCR validation.

**Supplementary Table 8. IPA generated enriched canonical pathways**

**(A)** 467 Unique Duodenal CR DEGs ($p < 10^{-4}$)

| Ingenuity Canonical Pathways | Molecules[a] | p-value |
|---|---|---|
| LPS/IL-1 Mediated Inhibition of RXR Function | 20 | 4.17E-08 |
| Xenobiotic Metabolism Signaling | 23 | 6.76E-08 |
| PXR/RXR Activation | 11 | 7.59E-08 |
| Glutathione-mediated Detoxification | 6 | 1.20E-05 |
| Superpathway of Melatonin Degradation | 8 | 1.74E-05 |
| Estrogen Biosynthesis | 7 | 2.14E-05 |
| Antigen Presentation Pathway | 6 | 3.24E-05 |
| Nicotine Degradation III | 7 | 3.63E-05 |
| Triacylglycerol Degradation | 7 | 5.13E-05 |
| Interferon Signaling | 6 | 6.03E-05 |
| Aryl Hydrocarbon Receptor Signaling | 12 | 6.31E-05 |
| Melatonin Degradation I | 7 | 6.92E-05 |
| Acyl-CoA Hydrolysis | 4 | 8.71E-05 |

[a]Number of 467 Unique Duodenal DEGs in Category

**(B)** Cancer-associated Genes ($p < 10^{-4}$)

| Ingenuity Canonical Pathways | Molecules[a] | p-value |
|---|---|---|
| LPS/IL-1 Mediated Inhibition of RXR Function | 10 | 6.92E-07 |
| Sirtuin Signaling Pathway | 11 | 1.32E-06 |
| PXR/RXR Activation | 6 | 2.00E-06 |
| Xenobiotic Metabolism Signaling | 10 | 5.37E-06 |
| Sphingomyelin Metabolism | 3 | 1.55E-05 |
| LXR/RXR Activation | 6 | 8.13E-05 |

[a]Number of Cancer-associated Genes in Category

**(C)** Telomere Genes ($p < 0.01$)

| Ingenuity Canonical Pathways | Molecules[a] | p-value |
|---|---|---|
| Sirtuin Signaling Pathway | 5 | 9.77E-05 |
| UDP-D-xylose and UDP-D-glucuronate Biosynthesis | 1 | 3.63E-03 |
| RAR Activation | 3 | 4.37E-03 |
| PXR/RXR Activation | 2 | 4.79E-03 |
| Cell Cycle: G1/S Checkpoint Regulation | 2 | 6.03E-03 |
| Cyclins and Cell Cycle Regulation | 2 | 8.71E-03 |

[a]Number of Telomere Genes in Category

**(D)** Epithelial Cell Genes ($p < 10^{-4}$)

| Ingenuity Canonical Pathways | Molecules[a] | p-value |
|---|---|---|
| Superpathway of Melatonin Degradation | 8 | 1.95E-09 |
| Nicotine Degradation III | 7 | 1.12E-08 |
| LPS/IL-1 Mediated Inhibition of RXR Function | 12 | 2.69E-08 |
| Melatonin Degradation I | 7 | 2.88E-08 |
| Nicotine Degradation II | 7 | 3.39E-08 |
| Estrogen Biosynthesis | 6 | 2.45E-07 |
| Bupropion Degradation | 5 | 4.57E-07 |
| Xenobiotic Metabolism Signaling | 12 | 7.24E-07 |
| Acetone Degradation I (to Methylglyoxal) | 5 | 1.41E-06 |
| PXR/RXR Activation | 6 | 4.79E-06 |
| Glutathione-mediated Detoxification | 4 | 1.78E-05 |
| Glucose and Glucose-1-phosphate Degradation | 3 | 5.13E-05 |

[a]Number of Epithelial Cell Genes in Category

**(E)** Immune System Genes ($p < 10^{-4}$)

| Ingenuity Canonical Pathways | Molecules[a] | p-value |
|---|---|---|
| Antigen Presentation Pathway | 6 | 1.45E-07 |
| Interferon Signaling | 6 | 2.82E-07 |
| Type I Diabetes Mellitus Signaling | 8 | 3.24E-06 |
| Communication between Innate and Adaptive Immune Cells | 6 | 2.29E-05 |
| T Helper Cell Differentiation | 6 | 2.51E-05 |
| Retinoic acid Mediated Apoptosis Signaling | 5 | 3.63E-05 |
| Crosstalk between Dendritic Cells and Natural Killer Cells | 6 | 3.89E-05 |
| iCOS-iCOSL Signaling in T Helper Cells | 7 | 3.98E-05 |
| Th1 Pathway | 7 | 4.57E-05 |
| iNOS Signaling | 5 | 5.25E-05 |

[a]Number of Immune System Genes in Category

**Supplementary Table 9. IPA canonical pathway: Sirtuin Signaling Pathway**

**(A)**

| Category | Molecules[a] | p-value |
|---|---|---|
| Cancer-associated | 11 | 1.32E-06 |
| Telomeres | 5 | 9.77E-05 |
| Epithelial | 8 | 8.51E-04 |
| 467 Unique | 15 | 1.38E-03 |

[a]Number of Genes in Category

**(B)**

| Ensemble ID | Gene Symbol | Gene Description | log(FC) | adj. P-value |
|---|---|---|---|---|
| ENSMUSG00000027513 | Pck1 | phosphoenolpyruvate carboxykinase 1, cytosolic | 4.39 | 1.62E-04 |
| ENSMUSG00000003849 | Nqo1 | NAD(P)H dehydrogenase, quinone 1 | 2.26 | 7.34E-04 |
| ENSMUSG00000026773 | Pfkfb3 | 6-phosphofructo-2-kinase/fructose-2,6-biphosphatase 3 | -2.19 | 1.01E-03 |
| ENSMUSG00000022383 | Ppara | peroxisome proliferator activated receptor alpha | 2.49 | 1.34E-03 |
| ENSMUSG00000005125 | Ndrg1 | N-myc downstream regulated gene 1 | 2.65 | 1.48E-03 |
| ENSMUSG00000015243 | Abca1 | ATP-binding cassette, sub-family A (ABC1), member 1 | 3.90 | 1.66E-03 |
| ENSMUSG00000042688 | Mapk6 | mitogen-activated protein kinase 6 | 1.51 | 1.98E-03 |
| ENSMUSG00000031583 | Wrn | Werner syndrome homolog (human) | -1.59 | 2.36E-03 |
| ENSMUSG00000020538 | Srebf1 | sterol regulatory element binding transcription factor 1 | 1.57 | 3.30E-03 |
| ENSMUSG00000039231 | Suv39h1 | suppressor of variegation 3-9 homolog 1 (Drosophila) | 1.65 | 5.63E-03 |
| ENSMUSG00000029167 | Ppargc1a | peroxisome proliferative activated receptor, gamma, coactivator 1 alpha | -1.65 | 1.22E-02 |
| ENSMUSG00000020826 | Nos2 | nitric oxide synthase 2, inducible | -2.92 | 1.39E-02 |
| ENSMUSG00000055116 | Arntl | aryl hydrocarbon receptor nuclear translocator-like | -1.87 | 1.56E-02 |
| ENSMUSG00000041238 | Rbbp8 | retinoblastoma binding protein 8 | 1.70 | 2.25E-02 |
| ENSMUSG00000026187 | Xrcc5 | X-ray repair complementing defective repair in Chinese hamster cells 5 | 1.58 | 3.01E-02 |

**(A)** Number of common genes in the enriched canonical pathway, the Sirtuin Signaling Pathway, per gene category subset. **(B)** 15 significant genes in the enriched ingenuity canonical pathway, the Sirtuin Signaling Pathway, for the duodenum mucosa samples of caloric restriction vs. *ad libitum* control mice (adj. p-value < 0.05 at |FC|>1.5). Fischer's exact test is used to calculate p-values in IPA. Sirtuin 3 was also downregulated in the telomere subset, but had FC = -1.38 and thus not counted in the 467 Unique DEGs.

**Supplementary Table 10. CR DEGs immune cell-type classification**

**(A)** human T cells

| Gene Symbol | Immune Cell Type | log(FC) | adj. P-value |
|---|---|---|---|
| Cth | Human T Regulatory Cells | 1.62 | 7.85E-03 |
| H2-Ea-ps | Human T Regulatory Cells | -2.08 | 2.32E-02 |
| Mfsd7c | Human T Regulatory Cells | 5.57 | 2.00E-04 |
| Npdc1 | Human All T cells | -1.55 | 2.98E-03 |
| Sesn1 | Human T Regulatory Cells | -1.58 | 6.33E-03 |
| Slc6a6 | Human T Regulatory Cells | 1.55 | 1.29E-02 |
| Zbtb16 | Human CD8+T Cells | 2.63 | 3.63E-03 |

**(B)** murine macrophages

| Gene Symbol | Immune Cell Type | log(FC) | adj. P-value |
|---|---|---|---|
| Cxcl10 | Murine M1 Macrophages | -2.03 | 3.80E-03 |
| Ddx60 | Murine M1 Macrophages | -2.13 | 1.49E-02 |
| Gbp6 | Murine M1 Macrophages | -2.15 | 1.81E-04 |
| H2-Q6 | Murine M1 Macrophages | -1.56 | 1.33E-02 |
| Herc6 | Murine M1 Macrophages | -3.23 | 1.01E-03 |
| Ifi44 | Murine M1 Macrophages | -4.94 | 3.04E-04 |
| Ifit1 | Murine M1 Macrophages | -3.36 | 2.57E-03 |
| Ifit2 | Murine M1 Macrophages | -2.30 | 6.83E-03 |
| Ms4a4c | Murine M1 Macrophages | -1.53 | 4.01E-02 |
| Mx2 | Murine M1 Macrophages | -1.62 | 1.64E-02 |
| Pecr | Murine Macrophages | 1.97 | 1.48E-02 |
| Stat1 | Murine M1 Macrophages | -1.66 | 2.37E-03 |
| Stat2 | Murine M1 Macrophages | -1.51 | 3.36E-03 |
| Tlr4 | Murine Macrophages | -1.77 | 1.23E-03 |
| Xaf1 | Murine M1 Macrophages | -1.71 | 1.22E-02 |

**(C)** human monocytes

| Gene Symbol | Immune Cell Type | log(FC) | adj. P-value |
|---|---|---|---|
| Aldh1a1 | Human Classical Monocytes | 2.45 | 1.76E-04 |
| Aldh1a7 | Human Classical Monocytes | 2.63 | 4.24E-04 |
| Fbp1 | Human Intermediate Monocytes | 1.67 | 4.50E-03 |
| Gbp6 | Human Intermediate Monocytes | -2.15 | 1.81E-04 |
| Mgst1 | Human Classical Monocytes | 2.37 | 3.45E-04 |
| Mosc1/Marc1 | Human Classical Monocytes | 3.07 | 1.06E-03 |
| Nkg7 | Human Intermediate Monocytes | -1.63 | 2.21E-03 |
| Rdh16 | Human Intermediate Monocytes | -1.54 | 6.18E-03 |
| Rdh9 | Human Intermediate Monocytes | 1.60 | 7.16E-03 |
| Scd1 | Human Intermediate Monocytes | 2.72 | 9.51E-03 |
| Setbp1 | Human Nonclassical Monocytes | -2.14 | 1.10E-03 |
| Tgm2 | Human Intermediate Monocytes | -3.10 | 4.48E-03 |

**(D)** human dendritic cells

| Gene Symbol | Immune Cell Type | log(FC) | adj. P-value |
|---|---|---|---|
| Cd36 | Human Monocyte Derived DC's | 1.86 | 2.21E-03 |
| Cxcl10 | Human Mature DC's | -2.03 | 3.80E-03 |
| Fcer2a | Human Monocyte Derived DC's | -1.90 | 7.30E-03 |
| Gbp6 | Human Mature DC's | -2.15 | 1.81E-04 |
| Ifit1 | Human Mature DC's | -3.36 | 2.57E-03 |
| Ifit3 | Human Mature DC's | -2.22 | 2.26E-02 |
| Mx2 | Human Mature DC's | -1.62 | 1.64E-02 |
| Oasl2 | Human Mature DC's | -2.37 | 6.78E-03 |
| Tnfsf10 | Human Mature DC's | -2.06 | 1.48E-03 |
| Usp18 | Human Mature DC's | -2.55 | 6.24E-03 |

**(E)** human neutrophils

| Gene Symbol | Immune Cell Type | log(FC) | adj. P-value |
|---|---|---|---|
| Sema3c | Human Mature Neutrophils | 1.51 | 1.03E-02 |

**(F)** 8 genes with multiple cell specific classifications

| Gene Symbol | Immune Cell Types |
|---|---|
| Nkg7 | Human B-cell and Human Intermediate Monocyte |
| Setbp1 | Human B-cell and Human Nonclassicial Monocyte |
| Cxcl10 | Murine M1 Macrophage and Human Mature DC |
| Gbp6 | Murine M1 Macrophage, Human Intermediate Monocyte, and Human Mature DC |
| Ifit1 | Murine M1 Macrophage and Human Mature DC |
| Ifit3 | Human B-cell and Human Mature DC |
| Mx2 | Murine M1 Macrophage and Human Mature DC |
| H2-Ea-ps | Human T Regulatory and Human B-cell |

**(G)** human B cells

| Gene Symbol | Immune Cell Type | log(FC) | adj. P-value |
|---|---|---|---|
| Abca1 | Human B-cells | 3.90 | 1.66E-03 |
| Bank1 | Human B-cells | -1.89 | 4.98E-02 |
| Bcl11a | Human B-cells | -1.54 | 3.74E-02 |
| Cd72 | Human B-cells | -1.80 | 5.71E-03 |
| Ciita | Human B-cells | -1.55 | 2.59E-02 |
| Cr1l | Human B-cells | -1.51 | 1.35E-02 |
| Ephx1 | Human B-cells | 2.24 | 3.59E-04 |
| Etv3 | Human B-cells | -1.52 | 8.55E-03 |
| H2-Dma | Human B-cells | -1.62 | 8.87E-04 |
| H2-Ea-ps | Human B-cells | -2.08 | 2.32E-02 |
| Ifi27l2a | Human B-cells | -1.54 | 1.93E-03 |
| Ifit3 | Human B-cells | -2.22 | 2.26E-02 |
| Ly86 | Human B-cells | -2.04 | 5.78E-03 |
| Nkg7 | Human B-cells | -1.63 | 2.21E-03 |
| Parp14 | Human B-cells | -1.84 | 3.93E-03 |
| Rab30 | Human B-cells | 2.32 | 1.22E-02 |
| Setbp1 | Human B-cells | -2.14 | 1.10E-03 |
| Sp140 | Human B-cells | -1.88 | 7.34E-04 |
| Spib | Human B-cells | -1.62 | 2.90E-02 |

Enriched genes in immune cells (>2-fold difference) for human and mouse expression profiles when immune cell types were compared. Human genes were converted to mouse orthologs using MGI Batch Query.

**Supplementary Table 11. Gene symbols associated with multiple probe sets**

| Ensemble ID | Gene Symbol | # of probe sets | Gene Accession | Gene Description | log(FC) | adj. P-value |
|---|---|---|---|---|---|---|
| ENSMUSG00000052392 | Acot4 | 2 | NM_134247 | acyl-CoA thioesterase 4 | 2.34 | 2.87E-03 |
| ENSMUSG00000027597 | Ahcy | 3 | NM_016661 | S-adenosylhomocysteine hydrolase | 1.76 | 1.17E-02 |
| ENSMUSG00000028028 | Alpk1 | 3 | NM_027808 | alpha-kinase 1 | -2.64 | 4.88E-04 |
| ENSMUSG00000071324 | Armc2 *isoform B | 2 | NM_001034858 | armadillo repeat containing 2 | -1.88 | 8.52E-04 |
| ENSMUSG00000071324 | Armc2 *isoform A | | | armadillo repeat containing 2 | 1.65 | 5.01E-03 |
| ENSMUSG00000078566 | Bnip3 | 2 | NM_009760 | BCL2/adenovirus E1B interacting protein 3 | 2.20 | 8.95E-03 |
| ENSMUSG00000059005 | Hnrnpa3 | 9 | NM_053263, NM_146130, ENSMUST00000111964 | heterogeneous nuclear ribonucleoprotein A3 | 1.69 | 2.21E-03 |
| ENSMUSG00000028184 | Lphn2 | 5 | NM_001081298 | latrophilin 2 | -1.64 | 2.29E-03 |
| ENSMUSG00000032418 | Me1 | 2 | NM_008615 | malic enzyme 1, NADP(+)-dependent, cytosolic | 1.92 | 7.02E-04 |
| ENSMUSG00000066595 | Mfsd7b | 3 | BC010797, NM_001081259 | major facilitator superfamily domain containing 7B | 1.84 | 2.43E-03 |
| ENSMUSG00000032959 | Pebp1 | 2 | ENSMUST00000036951, NM_018858 | phosphatidylethanolamine binding protein 1 | 1.62 | 1.03E-02 |
| ENSMUSG00000032369 | Plscr1 | 2 | NM_011636 | phospholipid scramblase 1 | 2.81 | 1.54E-04 |
| ENSMUSG00000029167 | Ppargc1a | 2 | NM_008904, NR_027710 | peroxisome proliferative activated receptor, gamma, coactivator 1 alpha | -1.65 | 1.22E-02 |
| ENSMUSG00000070327 | Rnf213 | 9 | AK173199, ENSMUST00000131035 | ring finger protein 213 | -2.19 | 3.88E-03 |
| ENSMUSG00000027227 | Sord | 2 | NM_146126 | sorbitol dehydrogenase | 1.56 | 9.32E-03 |
| ENSMUSG00000070034 | Sp110 | 4 | NM_175397 | Sp110 nuclear body protein | -1.88 | 1.62E-04 |
| ENSMUSG00000037926 | Ssh2 | 2 | NM_177710 | slingshot homolog 2 (Drosophila) | -1.54 | 9.45E-03 |

Annotation of Armc2 isoforms:

| | | |
|---|---|---|
| Armc2 *isoform A | pos. regulated | GPL6246, 10368881 (ID_REF), NM_001034858, chr10:42008653-42008725 (SPOT ID), 2nd exon longest isoform |
| Armc2 *isoform B | neg. regulated | GPL6246, 10368859 (ID_REF), NM_001034858, XM_006512653, BC172168, chr10:41914993-42007709, (SPOT ID), longest isoform |

One gene, *Armc2*, had both upregulation and downregulation data, leading to the distinction of isoform A (upregulated) and isoform B (downregulated). The probe set with the lowest adj. p-value was chosen when a specific gene symbol had all upregulated or downregulated expression data for all associated probe sets.
